# Synergistic targeting of EP300/CBP and EYA co-activators collapses the rhabdomyosarcoma core regulatory circuit

**DOI:** 10.64898/2026.09.02.748955

**Authors:** AL Gustafson, S Nance, BE Gryder, NAM Shendy, L Wick, G McKay-Corkum, KE Ritter, SC Purdy, AR Wolin, SR Rosenbaum, S Mathavarajah, NA Demelfi, Y Wang, Y Zhang, MW Zimmerman, AM Kavariyani, EA Citarella, VJ Ebegboni, JW Hardin, A LaVeck, X Wang, NV Dharia, AL Hong, G Kugener, JS Boehm, JA Roth, J Khan, F Vazquez, KB Artinger, R Zhao, DM Langenau, J Qi, K Stegmaier, BJ Abraham, HL Ford, AD Durbin

## Abstract

Rhabdomyosarcoma (RMS) is a multi-subtype, high-risk pediatric sarcoma with a low mutational burden. The mutations found in RMS often alter genes involved in transcriptional control. Approaches to target dysregulated RMS transcription have remained elusive. Here, we develop a novel approach to target RMS transcription comprising simultaneous targeting of two distinctly acting transcriptional co-activators. We discover a common identity-controlling pan-RMS core regulatory circuit (CRC) composed of oncogenic and lineage-specific myogenic master transcription factors (mTFs). Using a super-enhancer-based reporter screen, we identify the EP300/CBP inhibitor A485 as a potent inhibitor of the pan-RMS CRC, though with efficacy-limiting toxicities. To enhance efficacy, we identify the mTF-binding co-activator EYA2 as a co-factor of this pan-RMS CRC and exploit a new second-generation EYA1/2 inhibitor, LG1-34, to disrupt its function. Combined co-activator inhibition inactivates the CRC and synergistically reduces RMS growth. This strategy dually targets CRC-associated co-activators to cooperatively suppress the RMS transcriptome and enforce cell death.

## Introduction

Rhabdomyosarcoma (RMS) is the most common pediatric soft tissue sarcoma^1^. Despite the use of aggressive multimodal therapies, including radiation, chemotherapy and surgery, children with metastatic or relapsed RMS continue to display poor survival^2,3^, and outcomes for these patients have not improved in decades^4^. Thus, there is a clear need for targeted agents in this pediatric population.

Pediatric RMS is subdivided molecularly into fusion-positive (FP-RMS) and fusion-negative RMS (FN-RMS) based on the presence of an oncogenic chromosomal translocation, most commonly between *PAX3* or *PAX7* and *FOXO1*^5^. FN-RMS commonly harbors mutations in receptor tyrosine kinases and p53 pathways^6,7^. Despite these differences, both FN- and FP-RMS share molecular features like dysregulation of *MYC* family transcription factor oncogenes, including amplification of *MYCN* in FP-RMS and trisomy of chromosome 8, which harbors the *MYC* locus^6,8^. MYC family transcription factors are known drivers of RMS^6,8–11^. Critically, regardless of mutational status or subtype classification, RMS tumors resemble embryonic skeletal muscle, an identity emphasized by elevated expression of myogenic master transcription factors (mTFs), such as MYOD1 and MYOG, TFs required for normal muscle differentiation^12^. Additional myogenic and cooperating mTFs, such as SIX1, SOX8, MYF5, MEF2D and the chimeric PAX3-FOXO1 fusion proteins have been proposed as drivers of RMS. However, the mechanisms by which these mTFs, many of which are required for normal myogenic differentiation, cooperate to establish and maintain the malignant RMS transcriptome remain poorly understood. Moreover, mechanisms to target these transcriptional drivers of RMS remain elusive.

Aberrant activation of transcription, driven by a small number of mTFs and their association with activating *cis-*regulatory elements, i.e. gene enhancers and promoters, is common in cancer^13–17^. Enhancers, which control gene expression during development, often become dysregulated in cancer^10,18,19^. In many tumor types, super-enhancers (SEs), large clusters of enhancers usually identified by H3K27ac-marked nucleosomes, are densely occupied by transcription factors, co-activators, and epigenetic regulatory machinery. These SEs regulate transcription of a small group of mTFs that bind to each other’s, as well as their own, SEs, creating an autoregulatory positive feedback loop, termed a core regulatory circuit (CRC), which drives a persistent proliferative cell state^17–20^. Overexpressed MYC family TFs further invade these enhancers, resulting in dramatically amplified transcriptional output and elevated expression of both oncogenes and cell-state-specific genes^13,18,19^. MYCN has been previously identified as a member of a proposed CRC driving FP-RMS proliferation, along with the lineage-specific transcription factors SOX8 and MYOD1^18^. Since *c-MYC* has increased copy number in RMS, we hypothesized that both c-MYC and MYCN drive SE activation and corresponding upregulation of myogenic and oncogenic mTFs in RMS, resulting in the formation of a CRC that drives a hyperproliferative state.

Here, we combine exome-scale functional genomics with targeted chromatin immunoprecipitation studies in nine RMS cell lines and a cohort of patient-derived xenografts (PDXs) to identify a pan-RMS core regulatory circuit. We identify a group of mTFs commonly required in both FP- and FN-RMS, including c-MYC, MYCN, ZEB2, SOX8, TCF12, MYOD1 and SIX1. Leveraging a super-enhancer-based reporter compound screen of 147 small molecules targeting epigenetic regulatory proteins, we identify A485, a histone acetyltransferase domain inhibitor of the co-activator proteins EP300/CBP, as a mechanism to destabilize CRC transcription, resulting in G1 cell cycle arrest and apoptosis, *in vitro* and *in vivo*. Unfortunately, A485 has efficacy-limiting toxicities *in vivo*. We therefore identify the tyrosine phosphatase EYA2 as a key co-factor of the mTF SIX1 and utilize a novel EYA1/2 tyrosine phosphatase inhibitor, LG1-34, to independently destabilize RMS CRC transcription and induce apoptosis. Combination treatment with A485 and LG1-34 causes synergistic inhibition of RMS growth *in vitro* and *in vivo*, significantly prolonging survival in a mouse model of RMS. These observations provide proof-of-concept that combined targeting of multiple enhancer-binding CRC co-factors can control the RMS transcriptome and induce RMS cell death, while also illuminating common transcriptional regulatory mechanisms spanning RMS subtypes.

## Results

### The selectively enriched dependency landscape of rhabdomyosarcoma includes members of the core regulatory circuitry

To identify which mTFs are required for transcriptional output in both FP and FN-RMS and thus may be members of a pan-RMS CRC, we analyzed the Broad Institute’s Cancer Dependency Map (DepMap 25Q2 release) exome-wide CRISPR-cas9/cas12 dropout screen. Screening was performed in 1183 distinct cancer cell lines, including 12 RMS lines: 8 containing *PAX-FOXO1* gene fusions (7 *PAX3-FOXO1*, 1 *PAX7-FOXO1*) and 4 without gene fusions. Comparing all RMS cell lines with all other cancer cells, we identified 140 genes that were selectively enriched for genetic dependency in RMS cells, compared to the remainder of cancer cell lines (effect size < 0, q-value < 0.05) **(Figure 1A, Supplementary Table 1)**. Curated gene ontology classification of these genes revealed an array of functions, with a striking enrichment for transcription factors (25/140, 18%, Fisher’s exact test compared to whole genome p-value = 0.0008) **(Supplementary Table 2)**. The majority of the most enriched and lineage-specific dependencies in RMS were transcription factors (TFs) **(Figure 1A, TFs noted in red)**, including *MYOD1*, *SOX8* and *MYCN*, which have previously been described as critical TFs in fusion-positive RMS tumors^18,21–24^.

**Figure 1.**
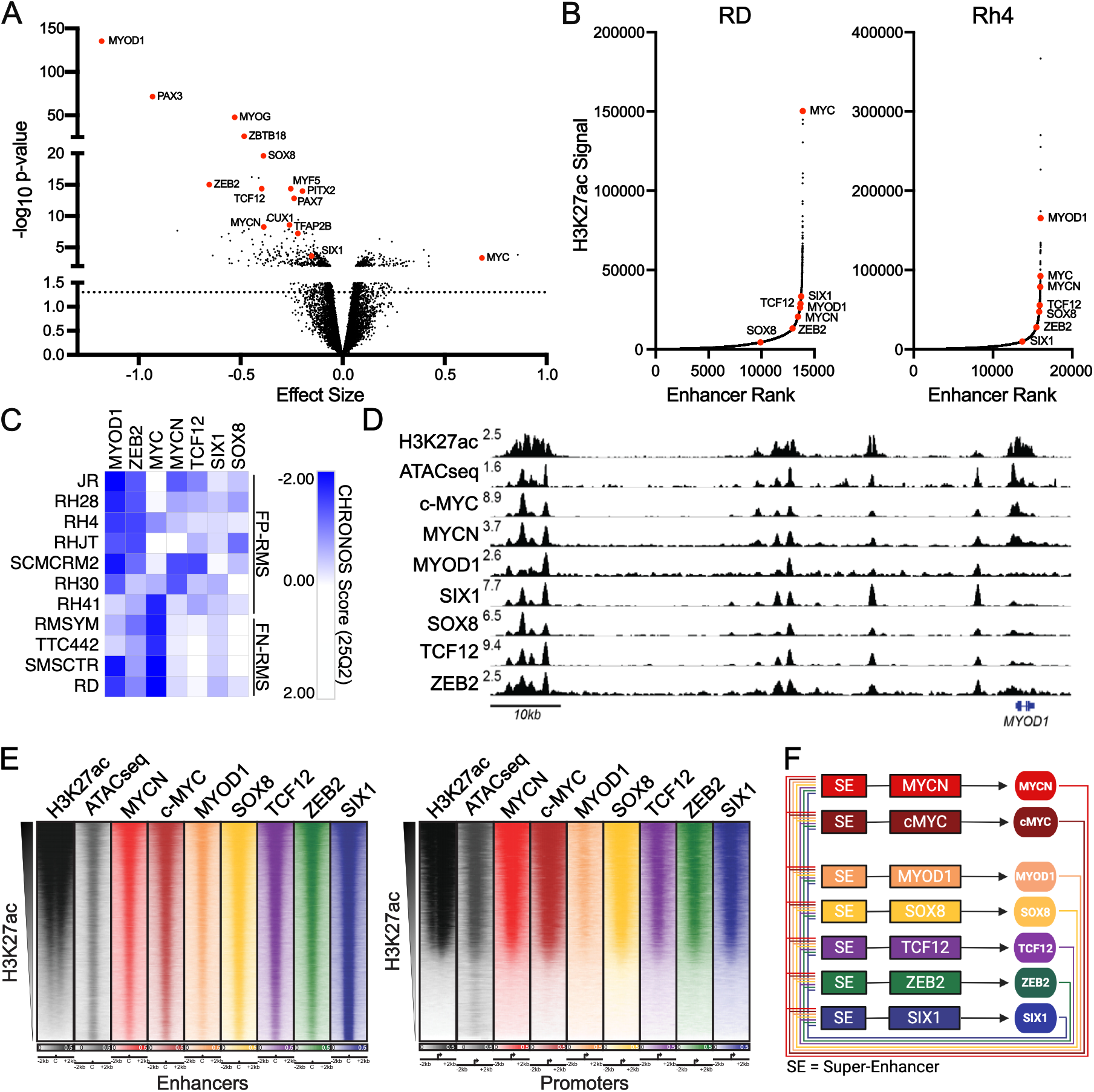
The selectively enriched dependency landscape of rhabdomyosarcoma includes members of the core regulatory circuitry. **A.** Volcano plot of enriched genes in 12 rhabdomyosarcoma (RMS) cell lines, compared to 1,171 non-rhabdomyosarcoma cancer cell lines. Data retrieved from www.depmap.org, 25Q2 release. Transcription factors enriched in RMS, and c-MYC are highlighted. **B.** Plots of H3K27ac signal, compared to enhancer rank, in RD and Rh4 cells. Highlighted are common transcription factors identified in >6/9 cell lines as being super-enhancer-associated. **C.** Dependency score (CHRONOS) in 11 RMS cell lines. Genes shown are genes regulated by super-enhancers in >6/9 RMS cell lines and are dependencies (CHRONOS Score < -0.5) in >2/11 RMS cell lines. **D.** ChIP-seq gene tracks at the *MYOD1* locus in RD FN-RMS cells, demonstrating co-binding of c-MYC, MYCN, MYOD1, SIX1, SOX8, TCF12, and ZEB2. Binding occurs in regions of open chromatin resolved by ATAC-sequencing, and within a super-enhancer, resolved by H3K27ac ChIP-seq. Data is representative of four independent RMS cell lines and all CRC loci. **E.** Genome-wide heatmaps ranked by H3K27ac signal at enhancers (left) and promoters (right) for the union of peaks bound by MYOD1, SIX1, SOX8, TCF12, and ZEB2. Binding of c-MYC and MYCN are also shown, and accessible chromatin is shown by ATACseq. Data are representative of four independent RMS cell lines. **F.** Model of core-regulatory circuitry in RMS cell lines showing super-enhancers (SE) controlling expression of CRC master transcription factor genes (boxes), resulting in autoregulatory transcription factor production (circles). Figure created with Biorender.com

To ascertain which TFs identified using DepMap data were super-enhancer associated, a standard heuristic for CRC members, we performed chromatin immunoprecipitation coupled to massively parallel high throughput sequencing (ChIP-seq). We used antibodies against H3K27ac in 10 RMS cell lines (3 fusion negative, 6 PAX3-FOXO1 positive, and 1 PAX7-FOXO1 positive). We used the Rank Ordering of Super-Enhancers (ROSE) algorithm to identify transcription factor genes associated with SEs (Methods, ChIPseq Analysis). We observed several genes, including *MYCN*, *MYOD1,* and *SOX8* in addition to *c-MYC*, *ZEB2, SIX1* and *TCF12* that were commonly marked by SEs in the majority of RMS cell lines **(Figure 1B, Supplementary Figure 1, Supplementary Table 3).** Next, we integrated DepMap data of genetic dependencies with this cohort of SE-associated TFs to identify potential CRC mTFs. Because CW9019 was the sole PAX7-FOXO1 cell line in our cohort, and any SE calls unique to this line could not be distinguished from cell-line-specific idiosyncrasies as opposed to PAX7-FOXO1-specific biology; we excluded it and identified all TFs regulated by SEs in >6/9 RMS cell lines. We then integrated these observations with DepMap data (CHRONOS Score < -0.5 in at least two RMS cell lines). These criteria narrowed the list of TFs to 7 candidate pan-RMS core regulatory circuit members: MYOD1, ZEB2, c-MYC, TCF12, MYCN, SIX1, and SOX8 (**Figure 1C**). Surprisingly, MYOG, a TF identified as a candidate mTF in FP-RMS previously^21^ and a TF that is highly enriched in RMS, was SE-regulated in only 3 RMS cell lines, suggesting it may play a role as a mTF in a more restricted subset of RMS. This is further supported by published scRNA-seq data in FN-RMS showing that *MYOG* is primarily expressed in less proliferative, “myocyte-like” tumor cells rather than in the tumor-propagating cell compartment^25^. We therefore excluded MYOG from the pan-RMS CRC, though MYOG likely plays a key role in FP-RMS specifically. We similarly considered whether the fusion oncoprotein PAX3-FOXO1 should be included as a lineage-specific mTF, but we found that *PAX3* was SE-associated in only 3/9 RMS cell lines, and FOXO1 was SE-associated in 5/9, excluding both based on our stringent selection criteria. Furthermore, CRISPR screening data did not distinguish deletion of *PAX3* or *FOXO1* from *PAX3-FOXO1*, confounding analyses of its essentiality. Previous work has shown that PAX3-FOXO1 is important for the initiation of a FP-RMS CRC, but that the fusion protein does not occupy the majority of active enhancers, suggesting that it does not operate as a full CRC member^21^. In contrast to *MYOG* and *PAX3-FOXO1*, *c-MYC* was negatively enriched in RMS compared with the rest of cancer cell lines (**Figure 1A**), focused examination of SEs and dependency scores across RMS cell lines using the criteria above demonstrates that *c-MYC* was also commonly a dependency gene and SE-associated in RMS, suggesting it is a potential mTF in RMS. The broad relevance of *c-MYC* across tumor types likely explains why it is not observed to be an RMS-specific dependency. Notably, *c-MYC* was observed as a dependency across both FP- and FN-RMS cell lines, while *MYCN* dependency was more prominent in FP-RMS lines (**Figure 1C**), consistent with subtype-specific differences in MYC-family-member-specific genetic events. Given that the DepMap RMS cohort includes only four FN-RMS cell lines, formal subtype-stratified analysis is not sufficiently powered to be informative. Together these analyses defined a final pan-RMS CRC comprising the 7 mTFs, *MYOD1, ZEB2, c-MYC, TCF12, MYCN, SIX1,* and *SOX8*.

Next, we sought to determine whether the 7 candidate mTFs bind at their own and each other’s super-enhancers to form a CRC. To do so, we performed ChIP-seq using commercially available and validated antibodies recognizing MYOD1, MYCN, c-MYC, SOX8, SIX1, TCF12 and ZEB2 in four RMS cell lines, including three fusion-positive (Rh4, RH28, and JR) and one fusion-negative (RD). Each factor bound in clustered regions within their own and each other’s SEs, in addition to gene promoters **(Figure 1D, Supplementary Figure 2)**. This binding occurred in sites of open chromatin, resolved by the Assay for Transposase Accessible Chromatin-sequencing (ATAC-seq) **(Figure 1D, Supplementary Figure 2)**. Consistent with our *in vitro* data, ChIP-seq data from a panel of RMS PDXs similarly demonstrated extensive stretches of H3K27ac at these same pan-RMS CRC associated loci in all samples (**Supplementary Figure 3**). These data are consistent with these factors forming a CRC, resembling CRCs identified in other tumors and untransformed cells^15–18,21,26–29^. In addition to locus-specific effects, CRC transcription factors colocalize at H3K27ac-marked, accessible chromatin across the genome to broadly regulate the transcriptome **(Figure 1E, Supplementary Figure 4)**. We also used publicly available ChIP-seq data generated using an antibody against the chimeric transcription factor PAX3-FOXO1 in the Rh4 cell line^30^ which demonstrated that this fusion oncoprotein colocalized with other members of the CRC at certain active enhancers **(Supplementary Figure 2)**. Together, these results indicate that transcription factors identified as pan-RMS CRC candidates bind shared regions genome-wide at H3K27ac-enriched sites, consistent with them acting as mTFs. Their binding at each other’s regulatory elements further suggests that they form a pan-RMS CRC (**Figure 1F**). Intriguingly, these data also indicate that both of the MYC family oncogenic transcription factors c-MYC and MYCN share properties of CRC mTFs in RMS cells.

### Inhibition of EP300/CBP-driven histone acetylation drives RMS death by inhibiting the RMS CRC

Next, we aimed to disrupt transcription of CRC members in RMS cells. Inhibitors that target epigenetic processes such as histone reader proteins, transcriptional cyclin-dependent kinases, as well as histone deacetylases and histone acetyltransferases, have previously been used to preferentially disrupt CRC transcription in a variety of diseases^17,18,21,31^. Since disruption of co-activator function has been shown to disrupt SE-mediated transcription in other settings^21,32^, we sought to inhibit CRC-mediated transcription in RMS by targeting epigenetic or transcriptional regulatory enzymes. We performed a screen of 147 epigenetic and transcription-targeted compounds (**Figure 2A, Supplementary Figure 5**) using validated Rh4 RMS reporter cell lines in which a portion of an intronic super-enhancer at the *ALK* locus was linked to luciferase (*ALK-SE-Luc*), and compared to a control Rh4 cell line expressing luciferase driven by a constitutively active CMV promoter^32^. The ALK SE reporter was determined to be an appropriate readout of CRC activity given that interrogation of our ChIP-seq data identified that all RMS CRC members bind to the intronic ALK enhancer in Rh4 cells (**Figure 2B)**. Previous work on the role of PAX3-FOXO1 in FP-RMS demonstrated that the ALK intronic SE is a readout of PAX3-FOXO1 transcription, and our ChIP-seq data demonstrates that PAX3-FOXO1 co-binds at the ALK intronic SE with the rest of the RMS CRC mTFs in Rh4 cells, supporting use of this reporter system to measure CRC-driven transcription (**Figure 2B**). We used DMSO as a negative control compound and actinomycin D as a positive control compound to generally inhibit transcription (**Supplementary Figure 5**). The most potent compound specifically suppressing *ALK-SE-Luc* activity was the inhibitor A485 **(Figure 2C, Supplementary Figure 6A)**. A485 is a highly selective spiro-oxazolidinedione competitive inhibitor of the histone acetyltransferase (HAT) co-activator proteins EP300 and CBP^33,34^. Consistent with literature reports, several inhibitors of histone deacetylases and bromodomains also displayed selectivity for *ALK-SE-Luc* activity, compared to CMV-Luc controls, illustrating that maintenance of the RMS CRC is regulated by numerous epigenetic regulatory enzymes, and is not solely regulated by EP300/CBP **(Supplementary Figure 6B)**^18,21,32^. To confirm that A485’s effects reflect on-target EP300/CBP HAT inhibition rather than off-target activity, we tested an expanded panel of CBP/EP300-directed compounds, including additional HAT inhibitors, bromodomain inhibitors, and PROTAC degraders, using the Rh4 ALK-SE reporter cell line. Two EP300/CBP HAT inhibitors, IHK-44^35^ and DS-9300^36^, showed markedly lower IC_50_ values than other EP300/CBP inhibitors. Two EP300/CBP bromodomain inhibitors (CCS1477^37^, GNE-049^38^) and three CBP/EP300 PROTAC degraders (JQAD1^39^, CBPD-409^40^, dCBP1^7^) showed reduced efficacy relative to the HAT inhibitors (**Supplementary Figure 7**). Together, these data indicate that suppression of the pan-RMS CRC is mediated by EP300/CBP HAT activity rather than bromodomain engagement or off-target effects, consistent with our prior work showing that the mechanism of EP300/CBP targeting shapes the resulting transcriptional phenotype^41^.

**Figure 2.**
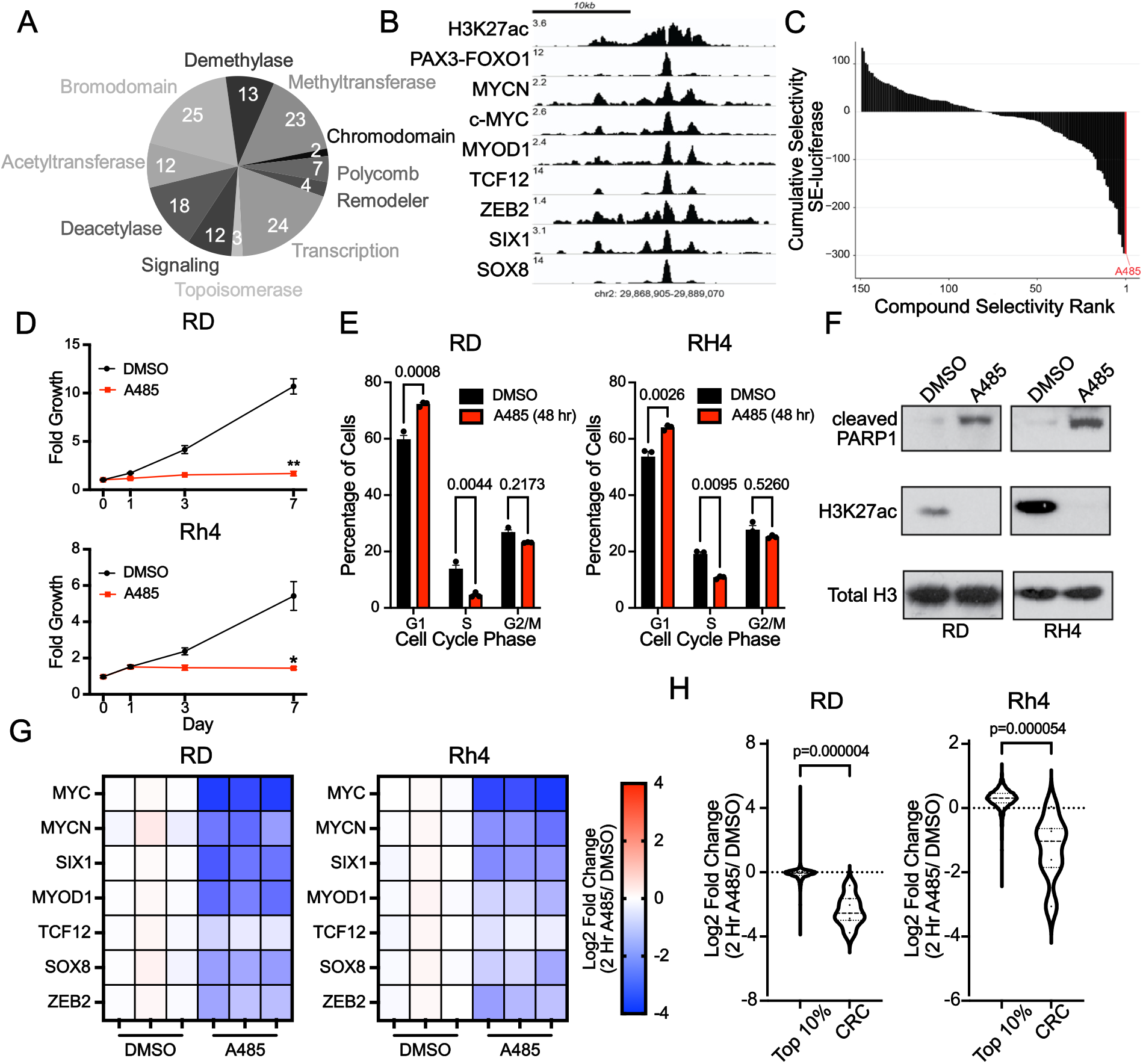
Inhibition of EP300/CBP-driven histone acetylation drives RMS death by inhibiting the RMS CRC. **A.** Pie chart demonstrating the functions targeted by each of the 147 epigenetically targeted compounds tested in an enhancer reporter luciferase compound screen. **B.** Super-enhancer reporter as in Gryder et al. *Nature Communications* 2019. Gene tracks demonstrated show the reporter locus in the *ALK* gene, bound by all RMS CRC proteins, which was cloned upstream of luciferase. **C.** Rh4 luciferase reporter cells were treated with 147 epigenetically targeted compounds, and selectivity for super-enhancer-driven luciferase signal downregulation, compared with non-SE-luciferase signal, was identified. The most potent compound identified causing selective downregulation of super-enhancer-driven signal was the EP300/CBP inhibitor A485. **D.** RD (top) and Rh4 (bottom) cell lines were treated with one dose of either 2μM (RD) or 500 nM (Rh4) A485 for 7 days, and relative cell growth quantified at day 0, 1, 3, and 7 by CellTiter-Glo assay. N=3, bars=S.E.M. Statistical difference was determined by Welch’s t-test. **E.** RD and Rh4 RMS cells were treated as in D for 48h and cell cycle was determined by propidium-iodide flow cytometry. n=3, bars=S.E.M. Statistics were determined by two-way ANOVA followed by a Sidak’s multiple comparison test. Data is representative of 4 independent cell lines. **F.** RD and Rh4 cells were treated as in D for 48h prior to protein extraction for western blotting. Data are representative of three independent protein isolations, treatments and western blots. **G.** RD and Rh4 cells were treated with 2μM (RD) or 500 nM (Rh4) A485 for 2h prior to RNA extraction and external spike-in normalized RNA-seq analysis. Shown are heatmaps demonstrating log_2_(fold change) at 2h, compared with DMSO treatment, in CRC mTF transcript abundance following treatment with A485. **H.** Analysis of RNAseq data from 2G, examining the response of the top 10% highest expressed genes in vehicle controls after 2h A485 treatment, compared to CRC gene response at the same time point. Statistical significance was assessed using a two-sample Kolmogorov-Smirnov test where p < 0.05 was considered significant. n=3 treatments and RNA isolation/sequencing per timepoint, treatment and cell line.

EP300 and CBP catalyze the acetylation of H3K27, which marks super-enhancers^42^. Prior evidence indicates that RMS cells are sensitive to levels of histone acetylation controlled by histone deacetylases^43^, and the results of our drug screen demonstrate that A485 inhibits SE-regulated transcription above baseline transcription. These findings led us to hypothesize that disruption of EP300/CBP activity may be a tractable method to selectively target SEs, suppressing the RMS CRC and, by proxy, its downstream targets. To test this hypothesis, we treated both FP-RMS (Rh4) and FN-RMS (RD) cells over 7 days with 5 µM A485, a concentration chosen because of its observed SE-luciferase suppression in our epigenetic compound screen **(Supplementary Figure 6A)**. Over 7 days of treatment, we observed potent growth suppression in both RMS cell lines (**Figure 2D**). We next determined the cell-line-specific IC_50_ doses of A485 in four RMS cell lines following 7 days of treatment (**Supplementary Figure 8A**). To determine the mechanism underlying the observed growth suppression, we treated four RMS cell lines with A485 at their IC_50_ dose for 48 hours and examined the effect on the cell cycle. A485 treatment induced a marked G1 cell cycle arrest and loss of the EP300/CBP target mark, H3K27ac (**Figure 2E,F, Supplementary Figure 8B,C**). A485 treatment also induced apoptosis, marked by cleaved PARP-1 (**Figure 2F, Supplementary Figure 8C**), a phenotype not mutually exclusive with the G1 cell cycle arrest observed in other cells within the same population. These data indicate that A485 potently inhibits EP300/CBP catalytic activity, reducing the H3K27ac mark, and leading to cell death.

Next, we sought to determine the acute effects of A485 on the RMS transcriptome. We hypothesized that disruption of EP300/CBP catalytic activity would result in changes in *cis*-regulatory SE elements, bound by and regulating the CRC genes. We treated 4 cell lines (JR, Rh4, Rh28, RD) with DMSO or A485 at the cell-line-specific IC_50_ value for 2 hours and then lysed cells for External RNA Controls Consortium (ERCC)-controlled RNA-sequencing. Comparing the normalized RNA-seq results after 2 hours of A485 treatment against their DMSO controls, we observed that all CRC genes were downregulated by A485 in each cell line, including both c-*MYC* and *MYCN* **(Figure 2G, Supplementary Figure 9A)**. Because highly expressed genes have greater basal transcriptional output, we controlled for the possibility that the 2-hour A485 treatment simply reduced total transcription globally, rather than selectively affecting the pan-RMS CRC, by comparing the changes in CRC expression to that of the top 10% highest-expressed genes in each cell line, a group that necessarily included all CRC mTFs. These data demonstrated that CRC genes were selectively downregulated by A485 treatment, as compared with the top 10% of expressed genes (**Figure 2H**, **Supplementary Figure 9B**). To characterize the highest-confidence common effects of A485 on gene expression across all RMS cell lines, we intersected all downregulated genes (log_2_ fold-change < -0.5, adjusted p < 0.05) found in >3/4 cell lines, after 2 hours of A485 treatment (n=370 genes). DepMap analysis identified that 61 (16.4%) of these downregulated genes were dependencies in >1 RMS cell line. Next, we performed a network analysis of these 61 genes using the STRING database to examine for potential interconnectivity. This analysis demonstrated a highly interlinked network of protein interactors, including all CRC members **(Supplementary Figure 9C)**. Enrichment analysis demonstrated the majority of these genes were involved in control of RNA polymerase II-based transcription in RMS cells (GO:0006357; FDR 1.07×10^-15^), and included other transcription factors with known roles in controlling the RMS malignant phenotype, such as *MYOG*^10,44^, *SNAI2*^45^, and JUN^46^. Gene set enrichment analysis of the Hallmark sets of the Molecular Signatures Database (MSigDB) demonstrated loss of myogenesis and MYC target gene sets in A485-treated cells, consistent with the observed effects (**Supplementary Figure 9D**). Taken together, these data indicate that treatment of RMS cells with A485 induces RMS cell cycle arrest and apoptosis, associated with preferential downregulation of the RMS CRC, including decreased expression of both c-*MYC* and *MYCN*.

### Re-expression of c-MYC stabilizes the effects of A485 on the pan-RMS CRC

MYCN and c-MYC control distinct but overlapping gene expression programs in different cancer types^47^. Importantly, in some contexts, these proteins may also functionally complement each other^48^. MYC family proteins have been implicated in stabilizing and driving CRCs in other tumor models^21,39^, and our epigenetic and dependency data indicate that both *MYCN* and *c-MYC* are relevant to RMS biology. We therefore hypothesized that A485-mediated downregulation of *c-MYC* and *MYCN* is a dominant driver of A485’s effect on RMS cells, and that this effect may be blunted by stable re-expression of MYC family proteins.

To test this hypothesis, we first established the expression patterns of both MYCN and c-MYC in RMS. We performed western blot analysis on a panel of RMS cell lines using antibodies specific to either MYCN or c-MYC and found that both proteins were detectable in the majority of RMS cell lines **(Figure 3A)**. These data were further supported by analyses of primary RMS tumor data, which showed that most tumors express both c-*MYC* and *MYCN*, in addition to pan-cancer analyses of cell line expression by RNAseq, which demonstrated that RMS cell lines commonly display expression of both c-*MYC* and *MYCN* **(Figure 3B, Supplementary Figure 10A).** To investigate expression of c-MYC and MYCN in RMS compared to normal skeletal muscle development we queried RNA-sequencing data taken from human skeletal muscle myoblasts (HSMM)^49^ and observed elevated c-MYC expression in myoblasts that declines with differentiation. In contrast, we observed increasing MYH3 expression associated with differentiation, and negligible expression of MYCN (**Supplementary Figure 10B**). This observation contrasts with the sustained co-expression of both c-MYC and MYCN we observed in RMS, suggesting that RMS cells aberrantly activate MYCN expression, and may retain developmentally expressed c-MYC.

**Figure 3.**
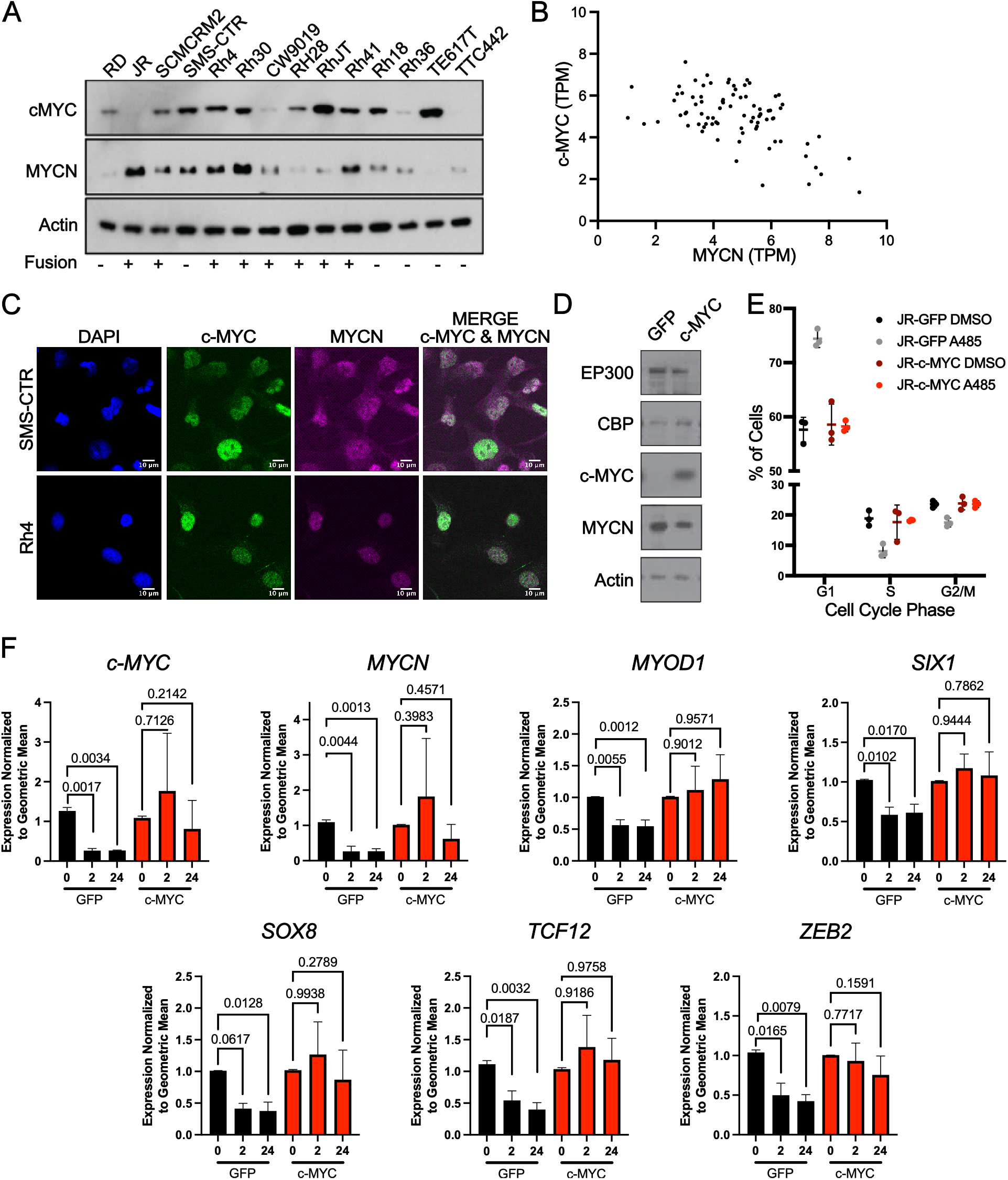
Re-expression of *c-MYC* stabilizes the effects of A485 on the pan-RMS CRC. **A.** Western blotting of RMS cell lines for c-MYC and MYCN protein expression. Actin is shown as a loading control. Fusion status is demonstrated. Data are representative of three independent lysates and blots. **B.** mRNA expression of *c-MYC* and *MYCN* in 84 primary RMS tumors. Data retrieved from sequence read archive accession SRP048220. **C.** Immunofluorescent detection of c-MYC and MYCN was performed on RMS cell lines (Rh4 and SMS-CTR). C-MYC is stained in green, MYCN in magenta, and DAPI in blue. Images are representative of ≥ 3 images and independent staining reactions. Bar = 10μm. **D.** JR RMS cells were stably transduced with GFP or c-MYC expressing lentiviruses. Western blotting of protein recovered from JR-GFP or JR-c-MYC cells are demonstrated. Data are representative of three independent lysates and blots. **E.** JR-GFP and JR-cMYC cells were treated with A485 or DMSO at 1 μM for 24h, prior to cell cycle analysis by propidium-iodide flow cytometry. n=3 independent replicates. **F.** JR-GFP and JR-c-MYC cells were treated with A485 or DMSO at 1μM for the noted timepoints, prior to RNA extraction and qRT-PCR. n=3 independent treatments and qRT-PCR reactions per sample. Target expression was normalized to and 0 hr treatment per cell line. Data are normalized to the geometric mean calculated from loading controls *β-actin, HPRT,* and *GAPDH* (mean ± S.E.M., statistical significance measured by one-way ANOVA followed by post hoc Dunnett’s multiple comparisons test.

Taking into account the intratumoral heterogeneity of RMS cell lines^25,50–52^, we next sought to identify if c-MYC and MYCN proteins were expressed in different cells within a heterogeneous population, or alternatively in the same cell. To do so, we performed immunofluorescence analysis of the RMS cell lines Rh4 (FP-RMS) and SMS-CTR (FN-RMS), using antibodies specific to either c-MYC or MYCN. As controls, we performed similar analyses on the *MYCN-*amplified neuroblastoma cell line Kelly, which expresses solely MYCN, and the *c-MYC-*amplified osteosarcoma cell line G292cloneA141B1, which expresses solely c-MYC. We observed that c-MYC and MYCN are co-expressed in the same cells of Rh4 and SMS-CTR (**Figure 3C, Supplementary Figure 10C**). Leveraging an orthogonal approach to immunofluorescence, we performed RNAscope assays using probes specific to *c-MYC* or *MYCN,* each of which were marked by a specific chromogen. Using SMS-CTR cells, we performed positive and negative assay controls, staining for the positive control probe *PP1B* and the negative control *dapB* (**Supplementary Figure 10D**). To assess the specificity of probes for *c-MYC* or *MYCN*, we stained for *c-MYC* and *MYCN* in the control G292cloneA141B1 and Kelly cell lines used in previous immunofluorescence experiments. We observed positive staining for *c-MYC* in G292cloneA141B1 (teal chromogen) and no positive staining for *MYCN* (**Supplementary Figure 10E**). Conversely, *MYCN* was strongly positive in Kelly cells (pink chromogen) with little positive staining for c-*MYC* (**Supplementary Figure 10E**). After establishing low background signal in the RNAscope protocol, and the specificity of the *c-MYC* and *MYCN* probes, we performed RNAscope in two FN-RMS cell lines, RD and SMS-CTR, and five FP-RMS cell lines, Rh4, Rh41, Rh28, RhJT, and JR (**Supplementary Figure 10F**). We found that in agreement with our findings in immunofluorescence, *c-MYC* and *MYCN* are co-expressed in individual RMS cells. Further, analysis of ChIP-seq data demonstrated largely overlapping binding patterns for these proteins (**Figure 1D,E**), suggesting that MYC proteins bind overlapping loci within the RMS CRC. To understand whether c-MYC or MYCN have selectivity for certain *cis*-regulatory elements genome-wide, we queried our RMS cell line c-MYC and MYCN ChIP-seq data. We found by comparing c-MYC and MYCN binding at enhancers and promoters in RMS cell lines that there was no preference in binding levels. We observed a strong correlation (Spearman’s rank correlation coefficient for JR = 0.985, RD = 0.963, Rh28 = 0.960, Rh4 = 0.979) between c-MYC and MYCN signal at all annotated promoters in these RMS cell lines (**Supplementary Figure 10G**). This correlation held for enhancers, where, in these RMS cell lines, both MYCN and c-MYC demonstrate a strong correlation in binding at sites co-bound by the remaining CRC mTFs (Spearman’s rank correlation coefficient for JR = 0.974, RD = 0.938, Rh28 = 0.911, Rh4 = 0.967) (**Supplementary Figure 10H**). These analyses support a model wherein c-MYC and MYCN do not have preferences for specific binding sites but instead bind at the same activating *cis*-regulatory elements genome-wide.

To further investigate the hypothesis that c-MYC and MYCN are mechanistically interchangeable in the context of the pan-RMS CRC, we leveraged JR *MYCN-*amplified RMS cells which express primarily *MYCN* **(Figure 3A)**. We established JR cells that stably express either GFP or c-MYC, under the control of a retroviral promoter **(Figure 3D)**. As with parental cells, JR-GFP cells underwent G1 cell cycle arrest in response to A485 treatment **(Figure 3E)**. In contrast, JR-c-MYC cells were resistant to the effects of A485 on cell cycle **(Figure 3E)**. Importantly, expression of c-MYC did not change the expression of EP300 or CBP, indicating that any effects on response to A485 are not due to altered co-activator expression (**Figure 3D**). To evaluate the direct effects of A485 on CRC gene expression in these cells, we next performed qRT-PCR for each CRC member, in both GFP and c-MYC overexpressing cells, after treatment with DMSO or A485. While GFP-expressing control cells demonstrated loss of CRC mTF gene expression, this effect was severely blunted in c-MYC overexpressing cells **(Figure 3F)**. To understand if the effects of c-MYC on stabilizing CRC gene expression correlate with changes in histone acetylation, we performed CUT&RUN for H3K27ac in JR-GFP and JR-MYC cells treated for 2 hours with A485. Paralleling our qRT-PCR findings, A485 caused profound loss of H3K27ac at this timepoint, which was blunted by the overexpression of c-MYC (**Supplementary Figure 11**). These findings indicate that MYC overexpression is associated with both stabilized CRC gene expression and histone acetylation. These data support a model in which c-MYC expression stabilizes CRC gene expression and can substitute for reduced MYCN, permitting continued oncogenic transcription and cell cycle progression in RMS cells.

### A485 reduces tumor growth in multiple *in vivo* RMS models

Since A485 potently induces G1 cell cycle arrest and apoptosis, driving growth suppression *in vitro*, we next sought to test its effects *in vivo.* To do so, we took advantage of an immunodeficient zebrafish model for xenotransplantation of RD FN-RMS and Rh41 FP-RMS cell lines^53^. To establish if A485 functions similarly *in vivo* as *in vitro*, we used cell lines containing the fluorescent ubiquitin-based cell cycle indicator (FUCCI) system^54^ (**Supplementary Figure 12A**). FUCCI4 stably transfected RD and Rh41 cell lines were treated *in vitro* with 2 µM and 1 µM A485 respectively for 24 hours and monitored for cell cycle dynamics by fluorescent imaging (**Supplementary Figure 12B**). A485 treatment led to a pronounced accumulation of cells in the G1 phase, as indicated by a strong increase in mKO2-Cdt1 signal and a reduction in Clover-Geminin signal. Quantitative analysis revealed a significant increase in the percentage of G1-phase cells compared to vehicle-treated controls, confirming that A485 induces G1 arrest in RD and Rh41 FUCCI4 cells (**Supplementary Figure 12C**). Next, we periorbitally transplanted FUCCI4-expressing RD and Rh41 cells into *rag2^-/-^;il2rga^-/-^* immune-compromised zebrafish. Tumor-bearing animals were orally gavaged with A485 at 5mg/kg once per day after tumor transplantation. Tumor-bearing zebrafish were then monitored by live fluorescent microscopy (**Figure 4A**). As *in vitro*, RD and Rh41 xenografts showed a G1 growth arrest within 18-24 hours following A485 treatment **(Figure 4B,C, Supplementary Figure 12B,C)**. Moreover, fish treated with A485 displayed significant reductions in the average number of tumor cells over time. Quantification of fluorescent cells in each image demonstrated a significant decrease in tumor volume in A485-treated RMS cells compared to PBS-treated controls (**Figure 4D**). These data strongly indicate that A485 has similar effects on tumor cell growth *in vitro* and *in vivo.* To further explore these findings, we xenotransplanted Rh4 FP-RMS cells into the flank of NSG mice. After allowing the tumors to reach 100mm^3^, we treated the mice with either vehicle or with 75 mg/kg A485 daily by i.p. injection. This lower A485 dose (75 mg/kg) was chosen following observed toxicity in NSG mice with prolonged treatment of the previously reported maximally tolerated dose of 100mg/kg i.p. twice daily^34^. Mice were treated with vehicle or A485, tumor growth curves were fit to an exponential growth model, and growth rate constants were compared between vehicle and A485 treated mice using an extra sum-of-squares F-test (p-value = 4.8×10^-7^) (**Figure 4E)**. Mice were euthanized when tumors reached 2000 mm^3^. We observed that treatment with A485 decreased tumor growth and prolonged animal survival (log-rank p-value = 0.0060) (**Figure 4E,F**). For three animals per group, we treated for 10 days with A485 or vehicle control and then sacrificed the animal and performed immunohistochemistry on the engrafted tumor. Consistent with A485’s effect on growth, examination of tumor tissue demonstrated reduction of the Ki67 marker of proliferation, and a concomitant increase in cleaved caspase-3, a marker of apoptosis **(Supplementary Figure 12D)**. Thus, A485 effectively reduces tumor growth *in vivo,* though is insufficient to cure animals as a single agent, likely due to dose-limiting toxicities.

**Figure 4.**
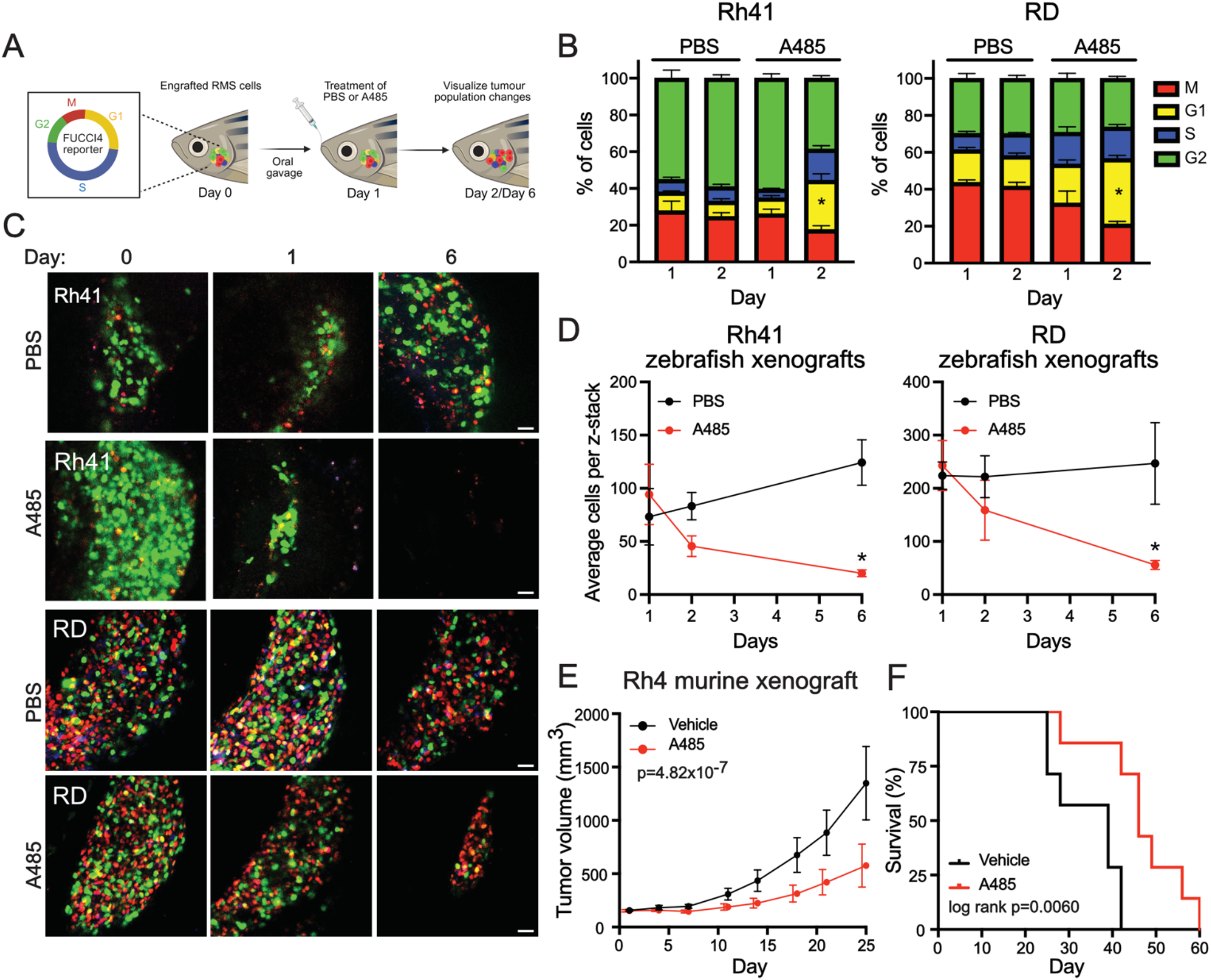
A485 reduces tumor growth in multiple *in vivo* RMS models. **A.** Diagram illustrating experimental design used for FUCCI4 reporter system in zebrafish RMS xenograft models. **B.** Quantification of cell cycle phases from Rh41 (left) and RD (right) cells stably expressing the FUCCI4 cell cycle reporter system xenografted into the retro-orbital space of *Rag2^-/-^IL2^-/-^* immune compromised zebrafish and orally gavaged with either PBS or A485 (1mg/kg) one day after engraftment. **C.** Representative fluorescent images of Rh41 (top) and RD (bottom) FUCCI4 reporter cells described in B, images are at cell engraftment (d=0), treatment (d=1), and 6 days post-engraftment. Images are representative of n≥3 animals per treatment group. **D.** Tumor growth measured by fluorescent imaging and then quantified by average cells per z-stack in Rh41 (left) and RD (right) xenografted zebrafish. N ≥ 3 animals per treatment group. * p<0.05. **E.** Growth curves of Rh4 xenografts in NSG mice treated with vehicle or A485 (75 mg/kg i.p. daily). Curves were fit to an exponential growth model, and growth rate constants were compared using an extra sum-of-squares F-test, p-value = 4.82×10^-7^. Bars = S.D. **F.** Percent survival of mice under each treatment condition, with euthanasia at tumor volume ≥ 1500 mm³. Log-rank p-value was calculated using Log rank (Mantel-Cox) test, Log-rank p = 0.0060.

### EYA proteins support the CRC through SIX1 interactions

Because the effects of A485 were profound *in vitro* but limited by toxicities *in vivo*, we sought to enhance its efficacy by identifying collaborators with the pan-RMS CRC to target in combination. To do so, we identified proteins that interact with pan-RMS CRC mTFs using the STRING database. We identified 107 interactors with different mTF members, and then determined that 29/107 were targetable using available small molecules. Of these targetable interacting proteins, many (13/29, 44.83%) were general epigenetic machinery, including the recurrently identified CBP/EP300 and HDAC1/2. Transcriptional regulators comprised 11/29 interactors, 4 of which were EYA family proteins (**Supplementary Figure 13A, Supplementary Table 4)**. The EYA transcriptional co-activators were of particular interest as interactors of SIX1, because SIX1 lacks a transactivation domain and so requires a co-activator, often an EYA family member, to recruit transcriptional regulatory machinery^55,56^. Further, EYA proteins contain intrinsic tyrosine phosphatase activity that can regulate SIX1 transcriptional activity^57^, which can be targeted using small molecule inhibitors^58^. EYA family members (EYA1-4) are typically expressed in the embryo and downregulated after completion of embryonic development. These proteins are often reactivated in cancer, therefore reducing the likelihood of unwanted side effects when therapeutically targeted^59,60^. Thus, we hypothesized that SIX1’s interaction with EYA proteins in the pan-RMS CRC may represent an alternative mechanism for disruption of CRC activities. To test this hypothesis, we first sought to establish which EYA proteins interacted with SIX1 in RMS cells. We performed co-immunoprecipitation (co-IP) followed by western blot analysis in three RMS cell lines (RD, SMS-CTR, and Rh4), which identified that EYAs1-3 interact with SIX1 in RMS cells (**Figure 5A, Supplementary Figure 13B**). While EYA4 was identified as an interactor in some cell lines (RD and SMS-CTR) the pull-down in Rh4 cells was inconclusive. To prioritize which EYA proteins may be most relevant *in vivo*, we re-analyzed published scRNAseq data from RMS patients^50^, which demonstrated that *EYA1*, *EYA2*, *EYA3*, and *EYA4* are expressed in RMS cells (**Supplementary Figure 13C**). Importantly, two of four EYA members were regulated by super-enhancers in at least 5 of the 9 RMS cell lines we investigated: EYA1, and EYA2 (**Supplementary Table 3**). Since prior medicinal chemistry experiments identified allosteric tyrosine phosphatase inhibitors with enhanced selectivity for EYA2 over other EYA proteins^58,61,62^, we prioritized EYA2 for investigation.

**Figure 5.**
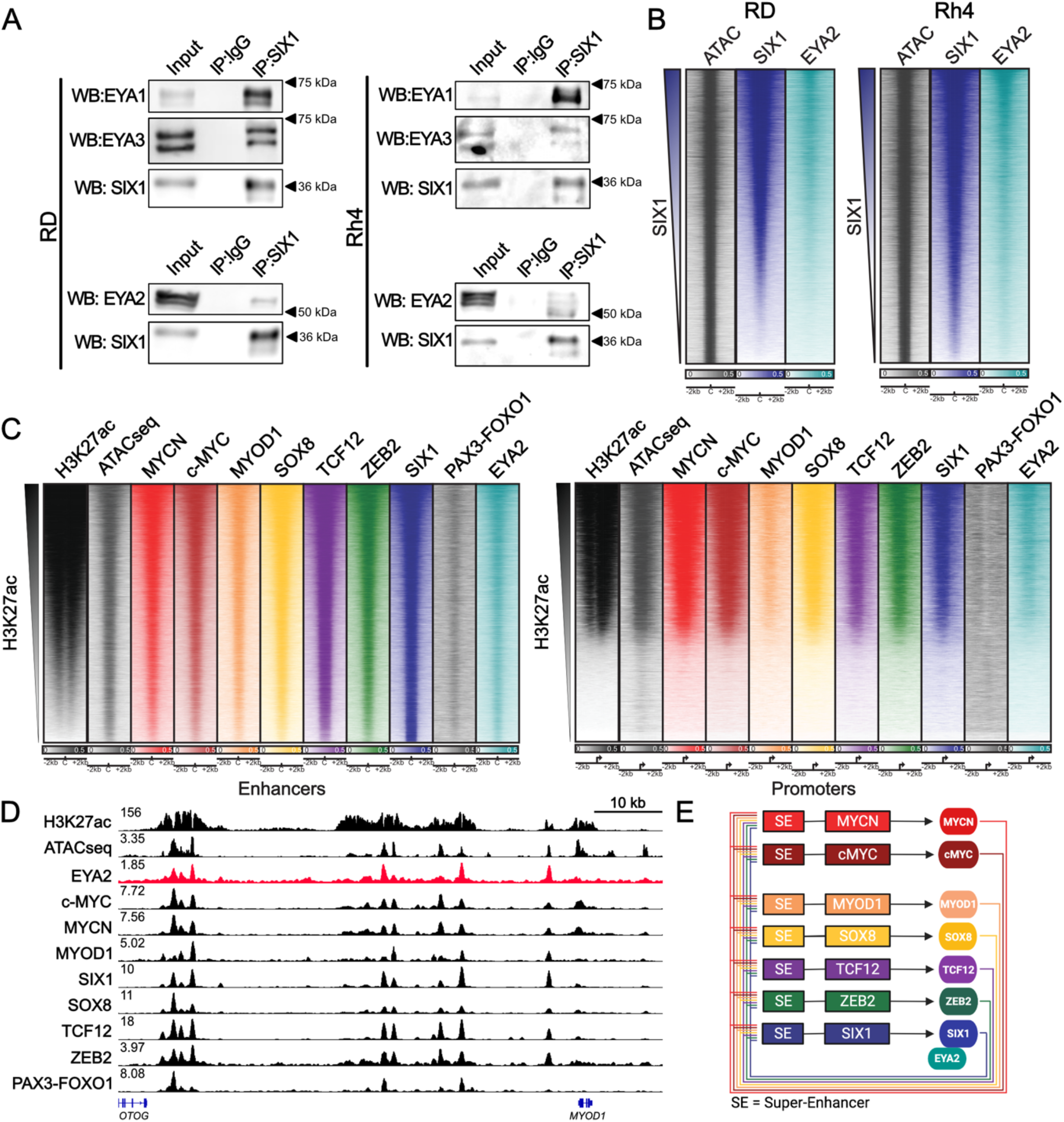
EYA proteins are SIX1-binding core regulatory circuitry co-factors. **A.** Co-immunoprecipitation for SIX1 in RD and Rh4 cells followed by western blot analysis where blots were probed for SIX1, EYA1, EYA2, and EYA3. Experiment was performed in 3 independent cell lines. **B.** Genome-wide heat maps of SIX1 and EYA2 binding in areas of open chromatin ranked by SIX1 signal in RD and Rh4 RMS cell lines. **C.** Genome-wide heatmaps ranked by H3K27ac signal at enhancers (left) and promoters (right) for the union of peaks bound by MYOD1, SIX1, SOX8, TCF12, and ZEB2. Binding of c-MYC, MYCN, PAX3-FOXO1 and EYA2 are also shown, and accessible chromatin is shown by ATACseq. Data are representative of two independent RMS cell lines (**Supplementary Figure 14B**). **D.** ChIP-seq coverage tracks at the *MYOD1* locus in Rh4 FP-RMS cells, demonstrating co-binding of EYA2, c-MYC, MYCN, MYOD1, SIX1, SOX8, TCF12, ZEB2, and PAX3-FOXO1. Binding occurs in regions of open chromatin resolved by ATAC-sequencing, and within a super-enhancer, resolved by H3K27ac ChIP-seq. Data are representative of two independent RMS cell lines. **E.** Model of core-regulatory circuitry in RMS cell lines with the addition of the CRC-associated co-factor EYA2, showing super-enhancers (SE) controlling expression of CRC master transcription factor genes (boxes), resulting in autoregulatory transcription factor production (circles). Figure created with Biorender.com

Since our co-IP experiments demonstrated an EYA2-SIX1 interaction (**Figure 5A**), we sought to understand if EYA2 bound coordinately alongside SIX1, genome-wide, when SIX1 is engaged with chromatin. Thus, we performed ChIP-seq analyses using antibodies against EYA2 in RD and Rh4 cells. Genome-wide, EYA2 and SIX1 demonstrated strong co-localization in RD and Rh4 cells in regions of open chromatin, identified by ATACseq **(Figure 5B)**. To determine the degree to which SIX1 and EYA2 bind shared sites genome-wide, we assessed all sites identified as significantly EYA2-bound or SIX1-bound in the ChIP-seq data produced in Rh4 cells. We observed that EYA2 and SIX1 largely bind the same genomic sites genome-wide and critically, we did not observe a population of high-confidence EYA2+/SIX1-sites or EYA2-/SIX1+ sites where signal was restricted to one factor alone (**Supplementary Figure 14A**). Next, we sought to understand if EYA2 similarly bound genome-wide with the CRC mTFs. We integrated our heatmap analysis of the CRC mTFs with binding of EYA2, which demonstrated that the interaction between SIX1 and EYA2 is found genome-wide alongside the pan-RMS CRC mTFs (**Figure 5C, Supplementary Figure 14A**). Examining all CRC loci, we observed that EYA2 is present at CRC mTF-bound peaks (**Figure 5D, *MYOD1* locus shown, Supplementary Figure 14B,C, Supplementary Figure 15)**. To confirm that the extensive co-localization among mTFs is not a general feature of ChIP-Seq datasets, we compared binding of the mTFs identified as CRC members in this study to previously performed ChIP-seq datasets for CTCF^63^ and analyzed all datasets at presumptive insulator *cis*-regulatory elements. We observed limited overlap between CRC associated ChIP-seq signal and CTCF ChIP-seq signal at activating *cis*-regulatory elements (**Supplementary Figure 16)**. Because EYA2 is encoded by a super-enhancer-associated gene, binds chromatin where CRC mTFs bind, but does not bind to DNA directly, and furthermore, because none of the EYAs individually are genetic dependencies in RMS, we term EYA2 a CRC-associated co-factor (**Figure 5E**). This property has been proposed before for the non-DNA-binding linker protein LMO1 in T-ALL and neuroblastoma^27^. Together, these data nominate EYA2 as a pan-RMS CRC co-factor that binds together with pan-RMS CRC mTFs.

### An allosteric EYA1/2 tyrosine phosphatase inhibitor targets the core regulatory circuit

We have previously identified EYA inhibitors that target EYA’s intrinsic tyrosine phosphatase activity^58,61,62,64^. Prior work from our group in other models using a novel allosteric <u>E</u>YA2 <u>T</u>yrosine <u>P</u>hosphatase Inhibitor (ETPi) has demonstrated that inhibition of EYA2’s intrinsic tyrosine phosphatase enzymatic activity decreases *c-MYC* mRNA and protein expression in other models^61^. Given the importance of MYC proteins in the RMS CRC, and co-binding of EYA2 with the CRC genome-wide, we hypothesized that use of an ETPi would be a targeted and potent method to suppress the RMS CRC. Our previous work showed that the compound 9987 is a potent allosteric inhibitor of the conserved EYA2 EYA domain (EYA2-ED) that sits at the interface between the enzyme’s haloacid dehalogenase (HAD)-core and helical cap subdomain (an α7–α8 loop region on the back face of the active site)^64^. The inhibitor is nestled directly behind the catalytic Asp residues and the Mg²⁺-binding site. By occupying this backside pocket, 9987 displaces the obligate Mg²⁺ through an induced conformational change of the EYA2 active site, effectively “locking” EYA2-ED in an inactive conformation that cannot coordinate and bind its obligate metal co-factor, thus inhibiting the tyrosine phosphatase activity^64^. Further, a phenylalanine (F290) in EYA2, close to the entrance of the 9987-binding pocket, is a tyrosine in all other EYAs. The larger size of the Tyr residue and a potential hydrogen bond between Y290 and residue T421 likely prevents 9987 from binding other EYA family members, contributing to the compound’s specificity towards EYA2 alone^64^. Based on the structure of 9987, we previously generated a congeneric series of analogs^58^. This effort yielded several improved inhibitors with greater potency against EYA2-ED tyrosine phosphatase activity by *in vitro* phosphatase assays^58^. We tested these different analogs and identified one, LG1-34, which contains an additional central thiophene (**Figure 6A**), exhibited enhanced potency (IC_50_ ∼ 0.18 μM) compared to the parental compound 9987 (IC_50_ ∼ 0.39 μM) in *in vitro* EYA2-ED phosphatase assays (**Figure 6B)**. Additionally, LG1-34 showed minimal off-target inhibition of endogenous cellular phosphatases, such as the HAD-like Ser/Thr phosphatase, SCP1, the metal-dependent protein tyrosine phosphatase, PPM1A, or the classical protein tyrosine phosphatase, PTP1B^58^. When comparing LG1-34 and 9987, 9987 shows efficacy against EYA2, but not other EYA family members, due to the reliance of the compound on a phenylalanine (F290) in EYA2 that is absent in other EYA family members^64^. In contrast, LG1-34 was active against three EYA paralogs, with biochemical IC_50_ values of 0.18 µM against EYA2 (compared to 0.39 µM for 9987), 2.79 µM against EYA1 (a >50-fold improvement over 9987’s ∼150.9 µM), and 34.63 µM against EYA3 (**Figure 6B, Supplementary Figure 17A**). LG1-34 exhibited no biochemical activity against EYA4, as previously reported^58^ (**Supplementary Figure 17B**). LG1-34’s increased spectrum of activity was particularly attractive, given that multiple EYA proteins interact with SIX1 in RMS, EYA1 and EYA2 are SE-regulated in RMS cells, and individual EYA proteins are not genetic dependencies in RMS, implying EYA proteins may play partially compensatory roles.

**Figure 6.**
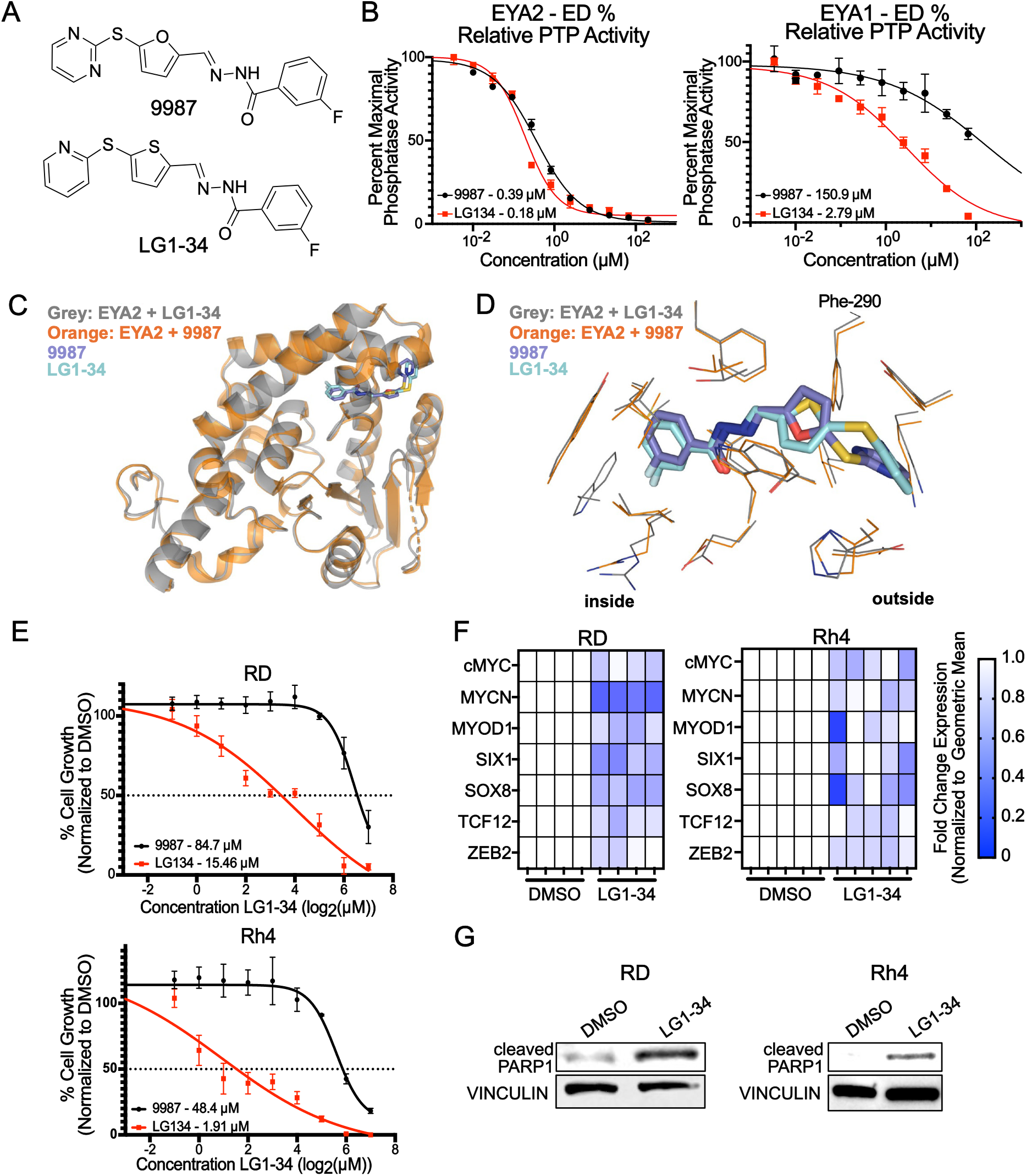
An allosteric EYA1/2 tyrosine phosphatase inhibitor targets the core regulatory circuit. **A.** Chemical structure of 9987 (top) and its analog, LG1-34 (bottom). **B.** Tyrosine phosphatase activity of purified EYA2-ED and EYA1-ED in the presence of increasing concentrations of 9987 (black) or LG1-34 (red). Enzymatic activity was measured by a fluorescent 3-O-methylfluorescein phosphate OMFP assay; data are plotted as percentage of activity relative to vehicle control. Dose–response curves (left) show that 9987 has minimal effect on EYA1 phosphatase activity, whereas LG1-34 potently inhibits both EYA1 and EYA2. Data are representative of at least three independent experiments (mean ± S.E.M). Statistical significance was determined by unpaired two-tailed t-test (P<0.05, *P<0.01). **C.** Ribbon diagram of the EYA2 C-terminal phosphatase domain (ED; gray) bound to LG1-34 (magenta stick model). The overall protein structure is shown superimposed on the previously solved EYA2–9987 structure (light blue, PDB: 5ZMA), indicating no global conformational changes upon analog binding. LG1-34 occupies an allosteric site distal to the active center (catalytic Asp residues in yellow; active-site Mg²⁺is not present in the inhibitor-bound structures). **D.** Close-up view of the allosteric pocket highlighting the distinct binding poses of LG1-34 versus 9987. EYA2 is shown as a surface and stick representation (key side chains Phe290 and Thr421 in green). LG1-34 (magenta) binds deeper and in a rotated orientation compared to 9987 (blue). Arrows denote the movement of the fluorobenzyl ring of the analog relative to the parent compound. **E.** RD (top) and Rh4 (bottom) were treated with a range of doses of LG1-34 (500nM – 128μM) for 72 hours. Cell growth was measured using CellTiter-Glo assay, values were normalized to DMSO control. Data points are representative of three independent experiments (mean ± SD). **F.** Heatmap of CRC gene expression in RD and Rh4 treated with LG1-34 at 15 μM (RD) and 2 μM (Rh4) (72h IC_50_ values) for 2 hours. Data are normalized to the geometric mean calculated from loading controls β-actin, HPRT, and GAPDH. Color represents the mean fold change of three technical replicates, and the results of >4 independent treatments and qRT-PCR reactions is shown. **G.** RD and Rh4 cells were treated with 15 μM (RD) and 2 μM (Rh4) LG1-34 for 48h prior to protein extraction for western blotting. Data are representative of three independent protein isolations, treatments and immunoblots.

To elucidate the differences in how 9987 and LG1-34 achieve their differing potency and specificity, we solved the co-crystal structure of EYA2-ED bound to LG1-34 (**Supplementary Figure 18, Supplementary Table 5)** and compared it with previously determined crystal structure of EYA2-ED bound to 9987^64^. LG1-34 binds to the same induced pocket in EYA2 as 9987 (**Figure 6C**). The global structure of EYA2 + 9987 compared to EYA2 + LG1-34 remains largely unchanged, with more noticeable differences localized to the residues within the allosteric pocket, active-site cleft, and the conformation of 9987 vs. LG1-34 (**Figure 6D**). Superimposition of the structures of LG1-34 or 9987 in complex with the EYA2-ED shows that LG1-34 adopts a different binding pose, marked by a rotation of the thiophene ring to pyridine ring and a slightly deeper insertion into the binding pocket, forming additional contacts with surrounding residues (**Figure 6D**). This altered compound binding pose is accompanied by a slight expansion of the binding pocket as side chains lining the pocket (including F290) shift to accommodate the compound (**Figure 6D**). In addition, LG1-34 contains a substituted sulfur in the thiophene ring, which is at a position that can potentially interact with the hydroxyl group of Y344 in EYA1 (equivalent to F290 in EYA2). These changes in the compound conformation and its interaction with surrounding residues (including potential interaction with Y344) may have contributed to the higher potency of LG1-34 against EYA1.

Given the broader spectrum of LG1-34, we next compared the effects of 9987 and LG1-34 across four RMS cell lines, RD, SMS-CTR, Rh4, and Rh30, where we observed a decrease in cell viability as measured by CellTiter-Glo assay after 3 days (**Figure 6E, Supplementary Figure 19A**). Consistent with the tighter binding of LG1-34 to EYA proteins, compared with 9987, we observed reduced IC_50_ values for LG1-34 compared to 9987 in all four RMS cell lines (**Supplementary Figure 19B**). Western blot analysis in parental RMS cell lines demonstrated a relationship between sensitivity to LG1-34 and expression of EYA2 (**Supplementary Figure 19C,D).** To examine if LG1-34 affected CRC expression, we next treated RD (low EYA2 expression) and Rh4 (high EYA2 expression) cells for 2 hours at the day-3 IC_50_ dose of LG1-34 and examined the effect on CRC gene expression by qRT-PCR (**Figure 6F, Supplementary Figure 19E**). LG1-34 treatment in RD and Rh4 cells reduced expression of all CRC mTFs **(Figure 6F, Supplementary Figure 19E)**. Consistent with these findings, LG1-34 treatment for 24 hours at the same dose caused reduced protein expression of CRC mTFs, without affecting EYA2 or a non-CRC protein control, vinculin (**Supplementary Figure 19F**). Continued treatment with LG1-34 for 48 hours resulted in activation of apoptosis, marked by PARP-1 cleavage in all four treated RMS cell lines (**Figure 6G, Supplementary Figure 19G**). These data suggest that the EYA2-biased tyrosine phosphatase inhibitor LG1-34 can disrupt CRC transcription in RD and Rh4 RMS cells, resulting in downregulation of the essential CRC and apoptotic cell death.

We next sought to investigate the mechanism through which disruption of the EYA1/2 tyrosine phosphatase activity inhibits the RMS CRC. We hypothesized that inhibition of EYA2’s tyrosine phosphatase activity inhibits SIX1’s ability to bind chromatin and function as a transcription factor. To investigate this, we treated Rh4 cells with 2 µM LG1-34 for 24 hours, isolated chromatin extracts, and performed western blot analysis for SIX1, EYA2 and H3K27ac. 24 hours of LG1-34 treatment reduced H3K27ac deposition as well as SIX1 and EYA2 chromatin binding (**Supplementary Figure 20A)**. To further evaluate the effect of LG1-34 treatment on SIX1/EYA2 binding, we performed ChIP-seq for SIX1 and EYA2 in Rh4 cells treated for 24 hours with 2 µM LG1-34. Treatment with LG1-34 for 24 hours resulted in a profound loss of genome-bound EYA2 and SIX1 (**Supplementary Figure 20B,C)**. Together these findings demonstrate that the EYA tyrosine phosphatase activity is necessary for SIX1/EYA2 chromatin occupancy, illustrating a potential mechanism through which LG1-34 disrupts RMS CRC maintenance.

### Combined targeting of EP300/CBP and EYA2 synergistically inhibits RMS growth

Given that A485 and LG1-34 are parallel inhibitors of the RMS CRC that operate in distinct fashions – either by inhibiting enhancer assembly or the tyrosine phosphatase activity of a CRC co-factor, we next sought to identify whether LG1-34 would enhance the effects of A485 on RMS growth suppression. We tested a dose-range of A485 and LG1-34 (2nM-20μM) alone and in combination in two RMS cell lines, RD and Rh4, and resolved their effects on cell growth by CellTiter-Glo assay at 1, 3, and 7 days of treatment. We first calculated the cell-based IC_50_ of each compound alone in RD and Rh4 cells, identifying suppression of growth in the micromolar range for both compounds **(Supplementary Figure 21A**). Next, we tested the effects of each compound alone or in combination across this dose range. Using an HSA synergy score metric^65^, the combination of A485 and LG1-34 synergistically reduced growth at doses below the single-agent IC_50_ for A485 and LG1-34 in both cell lines (**Figure 7A**). Further, while each individual compound caused suppression of cell growth over time, the combination enhanced these effects to a greater extent in high EYA2 Rh4 cells, as compared with low EYA2 RD cells **(Supplementary Figure 19C,D)**. To assess if the observed synergy between A485 and LG1-34 relies on on-target inhibition of EP300/CBP and EYA1/2, rather than off-target activity of the compound scaffolds we utilized structurally matched but catalytically inactive analogs of the parent compounds A485 (A486) and LG1-34 (LG1-37) in *in vitro* synergy experiments. We found that substitution with these inactive compounds abolished the synergy previously observed between A485 and LG1-34 (**Supplementary Figure 22, synergistic range shown with dotted black lines**). These data indicate that the combination of LG1-34 and A485 induces synergistic growth suppression, *in vitro*.

**Figure 7.**
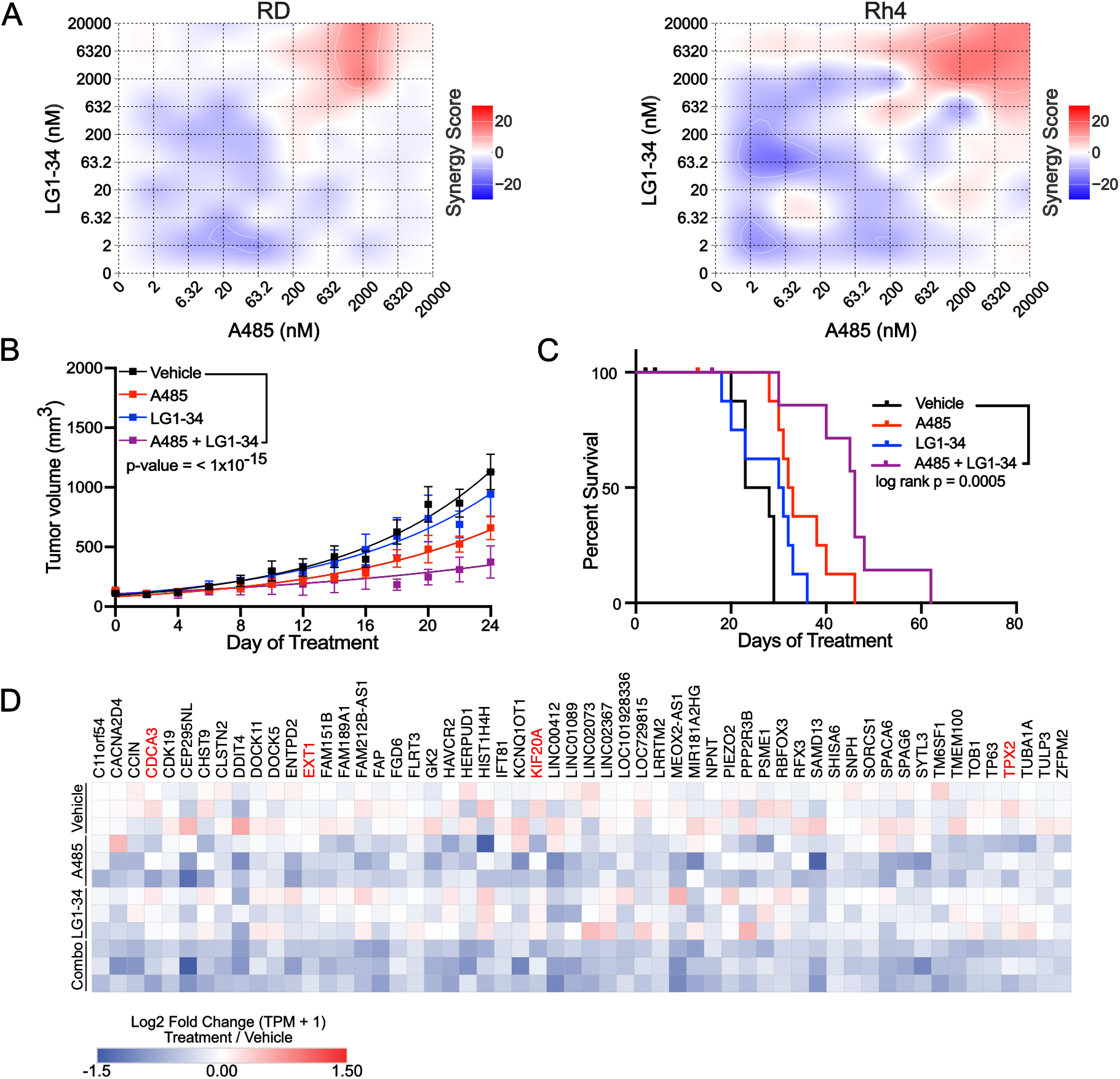
Combined targeting of EP300/CBP and EYA2 synergistically inhibits RMS growth. **A.** RD and Rh4 cells were treated with combinations of A485 (2nM – 20 μM) and LG1-34 (2nM - 20μM) for 72h (Rh4) and 6d (RD), prior to cell growth measurement by CellTiter-Glo assay. HSA synergy scores demonstrated. Data represent the average of three independent experiments. **B.** Growth curves of Rh4 xenografts in NSG mice treated with vehicle, A485 (75 mg/kg i.p. daily), LG1-34 (50 mg/kg oral daily), or A485 + LG1-34 at these doses. Curves were fit to an exponential growth model, and growth rate constants were compared using an extra sum-of-squares F-test. p-values: vehicle vs. A485, 3.116 × 10⁻^12^; vehicle vs. LG1-34, 0.1425; vehicle vs. combination, < 1 × 10⁻^15^; LG1-34 vs. combination, 4.157 × 10⁻^10^; A485 vs. combination, 1.136 × 10⁻^4^. **C.** Percent survival of mice under each treatment condition, with euthanasia at tumor volume ≥ 1500 mm³. Log-rank p-value was calculated for each condition using the Log rank (Mantel-Cox) test. Log rank p-values: vehicle vs. A485, 0.007; vehicle vs. LG1-34, 0.0682; vehicle vs. combination, 0.0005; A485 vs. combination, 0.0226; LG1-34 vs. combination, 0.0012. **D.** Genes that are down-regulated by combination treatment with A485 and LG1-34, as compared with A485 alone in vivo were identified by DESEQ2 analysis of RNAseq data. Dependency scores of these genes were identified in RMS cell lines (n=12) from DepMap (www.depmap.org, 25Q2 release). Genes that are dependencies in RMS cells (CHRONOS score <-0.5) are highlighted in red. Data shown represent RNAseq performed from tissue collected from three individual mouse tumors per treatment arm, treated for 7 days. Values represent log2 Fold Change TPM+1 in treatment groups normalized to vehicle controls.

Next, we sought to determine if LG1-34 could improve the tumor growth-suppression observed with A485 treatment *in vivo* in Rh4 xenografts. To do so, we first characterized the pharmacokinetic profile of LG1-34 in mice. At a single oral (PO) dose of 8 mg/kg, LG1-34 achieved a peak plasma concentration (Cmax) of 452 ng/mL (1.27 µM) at a Tmax of 0.25 h, with a plasma half-life (t1/2) of 1.63 h. Comparison with a 2 mg/kg intravenous (IV) reference cohort yielded an absolute oral bioavailability of 27.6% **(Supplementary Figure 23A).** Assuming dose-proportional pharmacokinetics, linear extrapolation to our efficacy dose of 50 mg/kg PO estimates a plasma Cmax of approximately 7.9 µM. This peak concentration exceeds the *in vitro* 3-day IC_50_ in Rh4 cells (2 µM) by roughly 4-fold, supporting this dose as sufficient to achieve transient target engagement despite the compound’s rapid clearance profile **(Supplementary Figure 23A). Therefore,** we engrafted Rh4 cells into the flank of NSG mice, and once tumors were >100mm^3^, treated tumor-bearing mice with vehicle alone, LG1-34 (50 mg/kg, oral, daily), A485 (75 mg/kg i.p. daily) or the combination of these agents **(Supplementary Figure 23B)**. Over the first 28 days of treatment, LG1-34 alone showed minor effects on tumor growth, while treatment with A485 demonstrated suppression of tumor growth, as previously observed **(Figure 4C,D).** The combination of LG1-34 and A485 led to enhanced growth suppression, compared with either agent alone or vehicle control (p-value < 1×10^-15^) (**Figure 7B**). Combined A485 and LG1-34 also led to prolongation of survival, as compared with single agent or vehicle controls (Log-rank p-value = 0.0005) (**Figure 7C**). As an assessment of compound combination toxicity, we examined mouse weights, which were equivalent across all groups **(Supplementary Figure 23C)**.

To examine the mechanism by which A485 and LG1-34 affect tumor growth *in vivo*, we performed an analogous experiment in a cohort of three mice per treatment arm. Tumors were collected from animals after one week of treatment and processed for immunohistochemistry (IHC) and RNAseq analyses. Consistent with the tumor growth curve results, A485 caused a suppression of Ki67^+^ proliferative cells that was enhanced in combination with LG1-34 (**Supplementary Figure 23D**). To confirm on-target specificity of A485, we performed IHC to the EP300/CBP mark H3K27ac, which showed reduced staining in A485 and A485 + LG1-34-treated tumors (**Supplementary Figure 23D**). Further, IHC for c-MYC demonstrated that combination of A485 and LG1-34 suppressed c-MYC protein expression (**Supplementary Figure 23D**). These data indicate that both A485 and LG1-34 drive synergistic growth suppression *in vivo*.

To more deeply examine the effects of A485, LG1-34 and the combination, we next performed RNAseq on the same tumors collected for IHC. We examined these data for changes in the transcriptome by DESEQ2^66^ analysis. Comparison of each group against vehicle control demonstrated, consistent with growth curve responses, profound changes in the transcriptome of A485 and combination-treated tumors, dominated by transcriptional downregulation **(Supplementary Figure 24A)**. In contrast, we observed very few changes in the tumors of mice treated with LG1-34 **(Supplementary Figure 24A)**. Focused examination of the CRC mTFs and EYA1-4 transcripts demonstrated that A485 alone and combination treated tumors displayed reduced expression of all CRC members **(Supplementary Figure 24B)**. Comparison of A485- and combination-treated tumors demonstrated generally correlated gene expression changes (Pearson’s R^2^=0.4495) **(Supplementary Figure 24C)**. To identify the changes in gene expression driven by the combination of A485 and LG1-34, compared with A485 alone, we performed DESEQ2 analysis of these conditions, and focused on genes selectively altered in each condition. Using an adjusted p-value cutoff of 0.05 and log_2_ fold change cutoff of <-0.5, we observed 566 overlapping genes between these treatments, with 449 genes uniquely downregulated by A485 alone and 56 genes uniquely downregulated by the combination of A485 and LG1-34 **(Supplementary Figure 24D)**. Consistent with our prior data, CRC genes were largely affected by both A485 and the combination of A485 and LG1-34 **(Supplementary Figure 24C)**. Examining the dependency scores for the 56 genes selectively downregulated by the combination of A485 and LG1-34, we observed loss of several RMS dependencies, including *CDCA3*, *EXT1*, *KIF20A*, and *TPX2*, which were unaffected in the absence of LG1-34 **(Figure 7D).** To identify pathways altered by combination treatment, we performed gene set enrichment analysis (GSEA) using Hallmark gene sets on genes differentially expressed with A485 + LG1-34 versus vehicle, identifying significant enrichment in several gene sets including HALLMARK_G2M_CHECKPOINT (p.adj = 1.55E-14, NES = -2.014), HALLMARK_MYC_TARGETS_V1 (p.adj = 3.36E-14, NES = -1.999) and HALLMARK_E2F_TARGETS (p.adj = 1.18E-13, NES = -1.965) (**Figure 7D, Supplementary Figure 24E**). These findings suggest that suppression of MYC target gene activation and inhibition of cell cycle progression and proliferation may contribute to the combinatorial efficacy of A485 and LG1-34. Analysis of our ChIPseq data for these 56 genes demonstrates binding of all CRC members at these sites, suggesting that a mechanism of transcriptional downregulation of these genes may rely on the effects of A485 and LG1-34 in decreasing the pan-RMS CRC’s enhancer activation at these sites (**Supplementary Figure 25A, example loci shown)**. To test this mechanism, we treated Rh4 cells with DMSO, A485, LG1-34 or the combination for 24 hours and performed CUT&RUN for H3K27ac. We found that H3K27ac deposition was lost significantly more at promoters and enhancers associated with these 56 downregulated genes in the combination treated samples than with LG1-34 or A485 alone (**Supplementary Figure 25A-D)**. These results indicate that combining epigenetic co-activator and transcriptional co-activator inhibition produces a mechanistically distinct effect on the RMS CRC, supporting further development of this co-targeting strategy for RMS.

## Discussion

Core regulatory circuitries (CRCs) sustain the malignant transcriptomes of cancer cells through the collective activity of a small number of lineage-defining or oncogenic master transcription factors^23^. While the component transcription factors (TFs) that comprise CRCs vary across different cell types, MYC family TFs recurrently appear in CRCs in numerous tumor types. MYC family TFs may play specialized roles in collaboration with tumor-specific mTFs in CRCs^17,21,27,67,68^ through their specialized functions as transcriptional amplifiers^69–71^. Because tumor CRCs are maintained by autoregulatory interactions between master transcription factors, enhancers, and promoters, they are particularly sensitive to perturbations of regulatory elements. However, directly targeting CRC mTFs has proven challenging, save for select cases, such as the degradation of IKZF1 in multiple myeloma using the molecular glue thalidomide^72^, or cases in which signaling transcription factors such as RARA or STAT proteins can be directly targeted^73–75^. Indirectly targeting CRCs through inhibition of general transcriptional or epigenetic machinery have been used effectively, but such treatments may be associated with toxicity because of off-target effects on pathways necessary for gene expression in untransformed cells. To this end, exome-wide CRISPR-knockout screening efforts have been useful to identify targets involved in epigenetic regulation of transcription beyond pan-lethal genes, that are instead required in distinct tumor states, such as the histone acetyltransferases EP300/CBP^39,41^, SAGA complexes^76^, or CDK8-containing complexes in the setting of RAS-mutant cancers^77^.

Here, we study rhabdomyosarcoma (RMS), a high-risk, lethal pediatric solid tumor, which is highly dependent on epigenetic regulation of transcription. RMS tumors are subclassified as either chromosomal fusion-positive (FP) or -negative (FN) subtypes. Presence of a chromosomal fusion often generates a chimeric transcription factor, PAX3-FOXO1 or PAX7-FOXO1, both of which dysregulate transcription^18,78,79^. Despite these differences, both FP- and FN-RMS resemble myogenic cells, with enhanced reliance on transcription factors such as MYOD1 and MYOG, which are critical proteins required for the regulation of normal myogenic differentiation^12^. By integrating functional genomics and epigenomic analyses, we identified a CRC that defines the transcriptome of RMS, regardless of fusion status, reflecting the myogenic state of these cells. Understanding the shared transcriptional dependencies between FP and FN-RMS therefore offers a unified therapeutic strategy despite differing driver mutations. Notably, several pan-RMS CRC mTFs, including SIX1 and MYCN, show oncofetal expression patterns, in that they are enriched during embryogenesis and largely absent from adult tissues, suggesting a favorable therapeutic window for targeting the pan-RMS CRC. This CRC is composed of several mTFs with key TFs significant during normal myogenic progression, including MYOD1, SIX1, TCF12, SOX8, c-MYC, and ZEB2. The RMS CRC also implicates both c-MYC and MYCN as critical, enhancer-invading, collaborating TFs. Capitalizing on the frequent observation that indirect inhibition of CRCs through targeting epigenetic and transcriptional regulatory enzymes has been a tractable method to disable CRCs, we use a high-throughput reporter assay-based screen to demonstrate that targeting enhancer regulation by the histone acetyltransferases (HATs) EP300 and CBP suppresses CRC-associated mTF expression in RMS. Building on this observation, we uncovered a newly characterized role for c-MYC in stabilizing the RMS CRC. Specifically, we show that c-MYC and MYCN proteins regularly bind the same CRC mTF-occupied loci and drive the malignant transcriptome. These findings reflect foundational studies in which targeted engineering of the MYCN locus to contain c-MYC sequences was sufficient to result in viable murine offspring, though some of the mice exhibited skeletal muscle defects, suggesting at least partially overlapping functions of these proteins^48^. To this end, we demonstrate that stable c-MYC expression is sufficient to rescue CRC mTF transcription following pharmacologic enhancer disruption with the EP300/CBP HAT inhibitor A485. These data underscore partially overlapping roles of MYC family oncoproteins in maintaining CRC activity through transcriptional amplification and enhancer invasion. They also provide an explanation for the observation of both c-MYC and MYCN expression in the same RMS cells, though in the absence of analogous rescue by *MYCN* expression, strictly redundant activities of c-MYC and MYCN cannot be ruled out. Intriguingly, c-MYC and MYCN proteins are known to counter-regulate each other in other settings, such that high MYCN expression results in transcriptional repression of MYC^47,80^. In both FP- and FN-RMS cells, chromosome 8, which bears the *MYC* locus, is commonly trisomic^5,6^, and FP tumor cells may also display focal *MYCN* amplification^11^. These observations indicate potential genetic mechanisms that may be responsible for disruption of these counter-regulatory mechanisms, permitting expression of both MYC and MYCN in the same cell, though further study is required to elucidate this point.

Importantly, targeting enhancer and promoter maintenance using the EP300/CBP HAT inhibitor A485 *in vitro* and *in vivo* was sufficient to induce a G1 cell cycle arrest and promote apoptosis in RMS cell lines. This observation is consistent with the concept of transcriptional addiction, wherein tumors with low mutational burden, such as RMS, become disproportionately dependent on *cis*-regulatory transcriptional programs and thus more vulnerable to their disruption than untransformed cells. While the effects of A485 were profound *in vitro* and in zebrafish xenografts, we observed dose-limiting toxicities in translating these findings to murine xenografts. Based on published A485 pharmacokinetics, estimated serum exposure at 75 mg/kg exceeds the Rh4 IC_50_ of 1 µM, though this serum exposure does not accurately reflect the concentration of A485 reaching tumor cells, given variables like protein binding and limited penetration of A485 from serum into the xenografted tumor. These data suggest, as previously indicated, that while EP300/CBP may have enhanced requirements in specific tumor settings, targeting the catalytic function of these proteins collectively may still produce toxicity in untransformed cells. To circumvent this issue, we present a novel therapeutic strategy that combines inhibition of master regulators of *cis-*regulatory elements with direct targeting of a CRC transcriptional co-activator. This strategy differs from previously reported strategies in which mTFs themselves are degraded, general enhancer co-activators inhibited, or signaling transcription factors targeted^39,72–75,81^, to instead target a new entity: mTF co-factors.

In the case of the RMS CRC, this dual targeting strategy was uniquely tractable, due to our identification of the SIX1 transcription factor as a critical CRC mTF. Previous work implicated SIX1 as required for restraining MYOD1 to stem/progenitor loci in RMS, with the loss of SIX1 resulting in MYOD1 relocalization and myogenic differentiation^82^. Here we extend the finding that SIX1 is necessary for RMS maintenance by demonstrating that SIX1 is an RMS CRC mTF and uncover EYA proteins as functional co-regulators of the pan-RMS CRC. Although EYA proteins are directly targetable, they exhibit functional redundancy, posing a therapeutic challenge. To overcome this issue, we used the novel compound LG1-34, which we demonstrate is an EYA2-biased, EYA1/2 tyrosine phosphatase inhibitor that disrupts RMS CRC function and leads to impaired growth and increased cell death *in vitro*. These effects were visible *in vitro* using this tool compound at micromolar level doses. Intriguingly, while LG1-34 alone showed limited efficacy *in vivo*, we observed enhanced tumor control and disruption of both the CRC and additional RMS-specific genetic dependencies when this agent was combined with A485. Consistent with on-target activity *in vivo*, LG1-34 treatment reduced Ki67, H3K27ac, and c-MYC by IHC, in line with the CRC disruption we observed *in vitro* and previously observed effects of the parental compound, 9987, on c-MYC^61^. The more modest transcriptional changes seen by bulk RNA-seq *in vivo* relative to *in vitro* data may reflect pulsatile LG1-34 exposure, and we note that direct biochemical confirmation of EYA1/2 phosphatase inhibition in tumor tissue was not performed, representing a limitation of our current pharmacodynamic characterization of LG1-34. Nonetheless, these findings are consistent with a model where combined targeting of CRC enhancer maintenance and the CRC-associated co-factor, EYA2, suppresses growth *in vitro* and *in vivo* by dysregulating the RMS CRC and subsequently the RMS malignant transcriptome. We note that combination treatment produced moderate tumor growth suppression rather than regression, and further compound optimization, of both A485 and LG1-34, will be required prior to any attempt at clinical translation.

Together, our findings define a novel epigenomic regulatory circuit driving RMS and provide a proof-of-concept for combinatorial therapeutic strategies that target general and tumor-specific CRC co-factors to overcome potential toxicities associated with targeting individual co-activators, while capitalizing on tumor cell dependency on core regulatory circuitry.

## Materials and Methods

### Cell Lines

All cell lines used for ChIPseq and in experiments investigating the effects of A485 alone (RD, JR, SCMCRM2, SMS-CTR, Rh4, Rh30, CW9019, RH28, RhJT, Rh41, Rh18, Rh36, TE617T, TTC442) were obtained from the Cancer Cell Line Encyclopedia through the Broad Institute of MIT and Harvard (RRIDs: CVCL_1649, CVCL_5916, CVCL_A770, CVCL_RT33, CVCL_8752, CVCL_A667, CVCL_0041, CVCL_N820, CVCL_VU81, CVCL_2176, CVCL_1659, CVCL_M599, CVCL_1755, CVCL_B255). Cell lines used to investigate the effect of LG1-34, and in mouse studies combining A485 and LG1-34 were gifted by Dr. Paul Jedlicka at the University of Colorado Anschutz Medical Campus (RRIDs: CVCL_0041, CVCL_5916, CVCL_1649, CVCL_A770). Prior to use in experiments, cell lines were short tandem repeat (STR)-tested to validate cell identity. RD (RRID:CVCL_1649) and SMS-CTR (RRID:CVCL_A667) cell lines were cultured in DMEM containing 10% heat-inactivated FBS. All other cell lines were cultured in RPMI containing 10% heat-inactivated FBS. All cell lines were screened routinely for *mycoplasma* contamination and only negative cells were used in experiments. Cell lines were used within 10 passages of thawing.

### DepMap Dependency Analysis and Gene Annotation

Analysis of dependency data was retrieved from the public DepMap portal (www.depmap.org) using the 25Q2 dataset. Two class comparisons of rhabdomyosarcoma (n=12: CW9019, JR, Rh28, Rh4, RhJT, SCMC-RM2, SMS-CTR, TTC442, Rh30, Rh41, RD, RMS-YM) vs. all other cell lines (n=1171) were performed using the custom analysis feature (Fig 1A). CHRONOS dependency scores were extracted for all genes and all cell lines (n =1183), and integrated with super-enhancer predicted gene loci to determine candidate master transcription factors (Fig 1C), or with RNAseq data to identify commonly downregulated dependency genes (Fig 7D) focusing on genes with dependency (CHRONOS Score <-0.5) in >1 RMS cell line. Details of individual cell lines are available at www.depmap.org. CCLE analyses of RNA expression were performed using the 25Q2 data release. Gene annotations were obtained from the PANTHER database annotations of molecular function^83^, followed by collapsing gene into curated processes as in^17^.

### RMS CRC Enhancer driven luciferase drug screening

A full description of creation and validation of SE construct was described previously^18^. Briefly, pGreenFire vector (with a minimal CMV promoter insufficient for basal transcription) from Systems Biosciences was modified by insertion of sequence from the CRC bound intronic SE within the ALK gene (shown in Figure 2B). RH4 cells were transduced with the lentiviral vector in a pooled fashion; cells were selected using puromycin for successful insertion of the construct. The counter-screen control vector, utilizing fully functional CMV promoter, was inserted into RH4 cells in the same fashion.

### Epigenetic Inhibitor Screen

Rh4 cells stably transfected with either the pALK-Luc construct (for reporting pan-RMS CRC activity) or a pCMV-Luc construct (reporter for impact on non-CRC related transcription) were used for assay development. A high throughput screening assay was then developed after optimization of cell seeding density, length of incubation of cells prior to and post treatment with test compounds, and effect of passaging among other factors. A pure compound library of 147 custom curated (and epigenetically focused) compounds was used in a medium throughput screening campaign for the identification of inhibitors of the pan-RMS CRC luciferase reporter activity in Rh4 ALK-Luc cells with minimal effect on activity in the Rh4 CMV-Luc reporter. DMSO solutions of screening library of compounds were thawed and used to prepare dose response concentration ranges (0.1 nM to 100 µM final concentrations) solutions in growth medium. Final DMSO concentration in assay wells was 0.2% or less. Each test concentration was assayed in quadruplicate. Rh4 ALK-Luc cells were seeded in white-walled and bottomed 384-well plates (Perkin Elmer, Cat# 6007658) for luciferase assays by transferring 27 µL of cell suspensions (seeding 3000 cells per well) into each well and transferred into an incubator for 18-20h. Rh4 CMV-Luc cells were similarly seeded into white plates and incubated. Screening compounds along with the positive (Actinomycin D, Sigma, Cat# A1410) and negative (DMSO) controls were added by transferring 3 µL of the prepared DMSO dilutions using an automated liquid handler (Agilent, Bravo). Treated plates were incubated for 24 h and allowed to equilibrate to room temperature for 30 minutes. SteadyLite Plus luciferase assay reagent powder (PerkinElmer, Cat# 6066759) was reconstituted in its buffer in parallel and was also allowed to equilibrate to room temperature. After transferring 30 mL of the luciferase assay reagent using a liquid handler, plates were further incubated at room temperature for 10 minutes. Finally, luminescence measurements were carried out using a multilabel microplate reader (BMG, Pherastar FSX) set in luminescence mode. Data was normalized to DMSO controls, and cumulative selectivity was calculated by taking the total difference in luciferase response across all concentrations (ie, ALK-luciferase signal minus CMV luciferase signal).

### CellTiter-Glo Growth Assays and Synergy Testing

CellTiter-Glo assays were performed following the manufacturer’s directions (Promega). In brief, 2,000 cells were plated in a 96-well format and treated with DMSO or varying compound concentrations diluted in culture medium. Cell growth was assessed after the reported treatment duration using luminescence measured by the CellTiter-Glo assay on a Molecular Devices SpectraMax iD3 Microplate Reader. For experiments testing the combination of A485 and LG1-34 *in vitro,* 1000 cells/well were plated in 384-well format and were treated with compounds dispensed using a TECAN D300e Digital Dispenser. Synergy was plotted using SynergyFinder^65^, using the HSA metric.

### Cell Cycle Analysis

RMS cells were treated with A485 or DMSO at the noted concentrations, then trypsinized and resuspend in a hypotonic citrate-PI solution for 30 minutes at 37°C, as described^84^. Nuclei were stabilized in 5 mol/L NaCl, and analyzed by flow-cytometry on a FACSAria II (BD Biosciences). Data analysis was performed using FlowJo v10.7 (BD Biosciences).

### Western Blotting

Cell lysates were collected by lysing cells in RIPA buffer (150mM NaCl, 1% NP-40, 0.5% DOC, 0.1% SDS, 50mM Tris pH 7.4) containing 1X protease inhibitor (Pierce Protease Inhibitor Tablets), or in Buffer “C” (10 mM HEPES, 3 mM MgCl_2_, 100 mM KCl, 0.01 mM EDTA, 10% glycerol). Protein content of samples was assessed using the Bio-Rad detergent-compatible (DC) protein assay. Samples were prepared to ensure equivalent total protein content and Laemmli Loading Buffer was added to a final concentration of 1X. Samples were resolved by polyacrylamide gel electrophoresis, followed by immunoblotting using primary antibodies targeting H3K27ac, Histone H3, c-MYC, MYCN, EP300, CBP, β-Actin, PARP1, MYOD1, SIX1, SOX8, TCF12, ZEB2, Vinculin, β-Tubulin, EYA1, EYA2, and EYA3 (RRIDs listed in **Supplementary Table 6**). Secondary antibodies were goat anti-rabbit or anti-mouse conjugated to horseradish-peroxidase (Licor). Chemiluminesence was detected using the Pierce ECL western blotting substrate and blots were imaged using an Azure 600 imaging instrument.

### Immunofluorescence

Immunofluorescence staining was performed with modifications as previously described^85^. Briefly, cells were seeded on sterile glass coverslips in six-well culture plates and incubated for 48 hours. Cells were fixed with 3% paraformaldehyde in PBS for 10 minutes at room temperature, permeabilized and blocked in 5% fetal bovine serum (FBS) in 0.3% Triton X-100 in PBS for 1 hour at room temperature, and incubated overnight at 4°C with primary antibodies against c-MYC (RRID:AB_1903938) and MYCN (RRID: AB_443533). Next, cells were washed and incubated at room temperature with DyLight anti-rabbit 488-conjugated secondary antibody (1:500; Thermo Fisher Scientific) and DyLight anti-mouse 526-conjugated secondary antibody (1:250; Thermo Fisher Scientific). Coverslips were mounted using ProLong Gold Antifade Mountant with DAPI (Thermo Fisher Scientific). Images were acquired using a Nikon A1 laser scanning confocal microscope equipped with a 40× Plan Fluor 1.3 NA oil immersion objective.

### RNAscope

Cell lines (Rh4, RD, JR, Rh41, Rh28, RhJT, SMS-CTR, Kelly, G292cloneA141B1) were pelleted and fixed in 10% neutral buffered formalin (StatLab, McKinney, TX). Fixed pellets were processed as tissue specimens using a HistoCore PEGASUS Tissue Processor (Leica Biosystems, Deer Park, IL), embedded in paraffin, and sectioned at 4 µm on a HistoCore AUTOCUT fully automated rotary microtome (Leica Biosystems). Duplex chromogenic in situ hybridization (ISH) for simultaneous detection of *MYC* and *MYCN* mRNA was performed using the mRNA Universal procedure on a Ventana Discovery Ultra automated stainer (Roche Diagnostics Corporation, Indianapolis, IN). RNAscope 2.5 VS probes (Advanced Cell Diagnostics, Newark, CA) targeting Hs-MYC (Cat. No. 311769) and Hs-MYCN (Cat. No. 417509) were applied, with Hs-PPIB (Cat. No. 313909) and dapB (Cat. No. 312039) serving as positive and negative control probes, respectively. Signal amplification and detection used the RNAscope VS Universal AP Reagent Kit (Cat. No. 323250) together with the DISCOVERY mRNA Sample Prep Kit (Cat. No. 08127166001), DISCOVERY mRNA Teal Detection Kit RUO (Cat. No. 8352941001), and DISCOVERY mRNA RED Detection Kit RUO (Cat. No. 07099037001; all Roche Diagnostics). *MYC* transcripts were visualized with the teal chromogen and *MYCN* transcripts with the red chromogen.

### Lentiviral Infection

All vectors were purchased from Addgene.org (pWZL-c-MYC-blast RRID:Addgene_10674, pWZL-GFP-blast RRID:Addgene_12269, psPAX2 RRID:Addgene_12260, pMD2.G RRID:Addgene_12259. Lentiviral particles were produced in HEK293T cells using lipofectamine 2000 (Thermo-Fisher Scientific) according to the manufacturer’s protocol. Viral supernatants were harvested and then used for infection with polybrene at a final concentration of 1 µg/mL (Sigma-Aldrich). Cells were selected using 5 µg/mL blasticidin (Gibco) and overexpression of c-MYC determined by western blotting. For FUCCI cell-cycle reporter lines, pLL3.7m-CloverGeminin (1-110)-IRES-mKO2-Cdt (30-120) (RRID:Addgene_83841), and pLL3.7m-mTurquoise2-SLBP (18-126)-IRES-H1-mMaroon1 (RRID:Addgene_83842) were transfected into HEK293T cells with 2mg pCMV-dR8.91, 0.2 mg pVSV-g and TransIT-LT1 reagent (Mirus Bio) as previously described to generate the FUCCI4 reporter system^86^. Supernatants containing the lentivirus were collected, filtered, and added to RD and Rh41 cells, in the presence of 4mg/mL polybrene (Millipore). Viral particle containing pLenti-CMV-GFP-puro was also added to cell lines for tracking cells *in vivo*. Viral particle containing pLKO.1-CMV-mKate2-Luc was added to RD and Rh41 and selected by FACS. Viral particle containing pLL3.7m-Clover-Geminin (1-110)-IRES-mKO2-Cdt (30-120) and pLL3.7m-mTurquoise2-SLBP (18-126)-IRES-H1-mMaroon1 was added to RD and Rh41 sequentially and selected by FACS.

### qRT-PCR

In figure 3, total RNA was harvested using Trizol (Life Technologies) in accordance with the manufacturer’s protocol. Complementary DNA synthesis was performed with Superscript II (Invitrogen). Quantitative PCR was performed using the Vii7 system (Life Technologies) with SYBR Green PCR Master Mix (Roche) and using validated primers specific to each target each gene. Two independent housekeeping controls were used to control for global effects on transcription. Primer sequences are displayed in **Supplementary Table 7**. For qRT-PCR performed in Figure 6, RNA was extracted from cells using Qiagen RNeasy Plus Micro Kit. Complementary DNA (cDNA) was then generated from purified RNA using the Bio-rad iScript reverse transcription kit (Bio-Rad). Quantitative reverse-transcriptase PCR (qRT-PCR) was performed using a Bio-rad CFX96 qPCR instrument in combination with the Biorad SsoFast Evagreen supermix and using validated primers specific to each target gene. Primer sequences are displayed in **Supplementary Table 7**.

### Co-Immunoprecipitation

Cells were grown to 70% confluency, prior to collection of pellets nuclear/cytoplasmic fractionation using the NE-PER Nuclear and Cytoplasmic Extraction Kit, as per the manufacturer’s instructions (Thermo Scientific). Nuclear lysates were dialyzed overnight at 4°C using the Slide-A-Lyzer MINI Dialysis 3.5K MWCO device (Thermo Scientific) in Buffer “A” (25 mM HEPES, 5 mM MgCl_2_, 25 mM KCl, 0.05 mM EDTA, 10% glycerol, 0.1% NP-40) with the addition of 1X protease inhibitor. Nuclear lysates were then mixed with Dynabeads M-270 beads conjugated to Rabbit polyclonal anti-SIX1 (RRID:AB_1079991) or Normal Rabbit IgG (AB_1031062) following the manufacturer’s instructions (Life Technologies). Nuclear proteins bound to the antibody linked Dynabeads M-270 beads were then eluted in 1X sample buffer and assessed by western blot analysis.

### Cloning, Expression, and Purification of Eya Proteins

The human Eya2-ED (residues 253–538) and equivalent residues for the Eya1-ED, Eya3-ED, and Eya4-ED were cloned into either pGEX-6P-1 (GE Healthcare) for crystallization studies or a pET28a vector (Novagen) for biochemical/phosphatase assays. For pET28a constructs, an N-terminal His₆-tag followed by a TEV cleavage site was introduced. For pGEX-6P-1, an N-terminal GST tag was used with a PreScission protease cleavage site. Constructs were transformed into *E. coli* BL21(DE3) cells. Cells were grown at 37 °C until OD₆₀₀ reached 0.6–0.8. Protein expression was induced with 0.3–0.4 mM IPTG at 18–20 °C for 16–20 h. Cells were harvested by centrifugation. Cell pellets were resuspended in lysis buffer [50 mM HEPES pH 7.5, 300 mM NaCl, 5% (v/v) glycerol, 1 mM DTT, 0.2 mg/mL lysozyme (ReadyLyse, Lucigen), 25 U/mL Benzonase (Sigma), and 1mL of 100x Xpert Protease Inhibitor Cocktail Solution (GenDEPOT) per 100mL buffer. Resuspended cells were lysed by sonication. After adjusting NaCl concentration to 500mM, lysates were clarified by centrifugation and filtered using a 0.45 µm filter. For His-tagged constructs, clarified lysates were applied to Ni²⁺-NTA agarose (Qiagen), washed with 20 CV of wash buffer [50 mM HEPES pH 7.5, 250 mM NaCl, 20 mM imidazole, 1 mM DTT], and eluted with 250 mM imidazole. For GST-tagged constructs, clarified lysates were incubated with glutathione sepharose beads (Cytiva) for 1 h at 4 °C, with flowthrough reapplied once to maximize binding. Beads were washed with lysis buffer, then incubated overnight at 4 °C with PreScission protease to cleave the GST tag. Cleaved protein was eluted with buffer [50 mM HEPES pH 7.5, 300 mM NaCl, 1 mM DTT], concentrated and applied to a Superdex 200 Increase 10/300 GL (Cytiva). Fractions corresponding to monomeric Eya2-ED were pooled, concentrated to 5–10 mg/mL, and flash frozen in aliquots at –80°C.

### In Vitro Phosphatase Assays

Phosphatase activity was quantified using 3-O-methylfluorescein phosphate (OMFP; Thermo Fisher) as described^87^. Assay buffer contained 50 mM HEPES (pH 7.5), 100 mM NaCl, 0.5 mM MgCl₂, 0.005% Tween-20, 0.05% BSA, and 1 mM DTT. For inhibitor assays, compounds were prepared as 10 mM DMSO stocks and serially diluted (100 µM-5nM) in DMSO. Compounds were added as 1% of the total assay volume to keep final DMSO concentration to ∼1%. Reactions (25 µL) were assembled in black low volume 384-well plates (Greiner), containing 50 nM Eya-ED protein and 100 µM OMFP. Reactions were allowed to proceed at room temperature protected from light for 60 minutes. Fluorescence was measured at 485 nm excitation/517 nm emission at 25 °C on a BioTek Synergy H1 plate reader.

### Crystallization

Purified Eya2-ED was pre-bound to compounds prior to crystallization. Protein (130 µM stock, 5 mg/mL) was diluted to 8 µM in optimized binding buffer [20 mM HEPES pH 7.5, 150 mM NaCl, 5% glycerol, 1.5 mM DTT, 5 mM TCEP, 0.5 mM EDTA]. Compound was added at a 2:1 molar ratio (protein:compound final concentrations 8 µM:16 µM). Complexes were incubated for 30 min on ice with intermittent gentle mixing, followed by 30 min at room temperature with occasional agitation. The complex was then reconcentrated to ∼130 µM (5 mg/mL) using Amicon centrifugal concentrators (10 kDa cutoff). This intermediate binding step was essential to achieve homogeneous complexes and reproducible crystallization.

Crystallization trays were set up using sitting-drop vapor diffusion method at 18 °C. 400 nL of Protein–compound complexes were mixed with 400 or 800 nL reservoir solution containing 0.1 M HEPES pH 7.5, 200 mM NaCl, and 20% (w/v) PEG-3350. 2 µL protein drops were equilibrated against 500 µL reservoir solution. Crystals appeared within 3–7 days. Optimal diffraction-quality crystals formed between 2.5–5 mg/mL of protein concentration while no crystals grew at ≥7.5 mg/mL.

### X-Ray Data Collection, Processing, and Structure Determination

Crystals were cryoprotected in reservoir solution supplemented with 20% glycerol and flash frozen in liquid nitrogen. Crystals were screened at the X-ray Crystallography Core Facility at the University of Colorado Anschutz Medical Campus. X-ray diffraction data were collected at beamline 8.2.1 of the Advanced Light Source (ALS, Lawrence Berkeley National Laboratory). Data were indexed and integrated with DIALS^88^ followed by scaling and merging with CARELESS^89^. Structures were solved by molecular replacement using the Phenix suite^90^ and the Eya2+9987 structure (5ZMA) as a search model. Iterative model building was conducted using both Phenix and Coot^91^, and model was refined in Phenix. Compound LG1-34 was fit into the electron density map using both the Phenix suite and Coot and validated against simulated annealing omit maps. Solvent molecules were added where justified by electron density. Final structures were validated with MolProbity^92^; all models contained >95% residues in favored regions and no residues in the disallowed regions of the Ramachandran plot (Supplemental Table 6). The final model of the Eya2 + LG1-34 structure is deposited in the Protein Data Bank with ID 9Y64.

### Animals

All animal studies were approved by local institutional review boards, including A485 vs. PBS zebrafish experiments (Massachusetts General Hospital), A485 vs. vehicle mouse experiments (Dana-Farber Cancer Institute) and A485 and LG1-34 combination experiments (University of Colorado Anschutz). Eight-week-old female and male NOD.Cg-Prkdc^scid^ H2-K1^b-tm1Bpe^ H2-Ab1^g7-em1Mvw^ H2-D1^b-tm1Bpe^ Il2rg^tm1Wjl^/SzJ (NSG) mice (The Jackson Laboratory, RRID:IMSR_JAX:025216) were used for mouse tumor xenograft experiments. For zebrafish tumor xenograft experiments, *rag2^Δ/Δ^;il2rga^Y91fs^*homozygous adult female and male AB and Casper (double mutant for *roy^a9/a9^* and *nacre^w2/w2^*)^93,94^ zebrafish were generated from incrossing *rag2^+/Δ^;il2rga^+/Y91fs^*heterozygous fish and outcrossing *rag2^+/Δ^;il2rga^+/Y91fs^* heterozygous fish with *rag2^Δ/Δ^;il2rga^Y91fs^*homozygous fish. Animals were identified following scale resection and genotyping at 2 to 3 months of age as previously described^93,95,96^. All immune compromised animals used in this study are kept at BL2 facilities with regular veterinary checks being performed.

### Rearing and Husbandry of *rag2^Δ/Δ^;il2rga^Y91fs^* Homozygous Zebrafish

For zebrafish tumor xenograft experiments, *rag2^Δ/Δ^;il2rga^Y91fs^* homozygous adult female and male AB and Casper (double mutant for *roy^a9/a9^* and *nacre^w2/w2^*)^93,94^ zebrafish were generated from incrossing *rag2^+/Δ^;il2rga^+/Y91fs^* heterozygous fish and outcrossing *rag2^+/Δ^;il2rga^+/Y91fs^* heterozygous fish with *rag2^Δ/Δ^;il2rga^Y91fs^* homozygous fish. Animals were identified following scale resection and genotyping at 2 to 3 months of age as previously described^93,95,96^ .Both males and females were used in these studies. Genotyped *rag2^Δ/Δ^;il2rga^Y91fs^* zebrafish were transferred to autoclaved, sterilized fish water containing penicillin G sodium salt (Sigma, 150 units/ml), streptomycin sulfate salt (Sigma, 100 mg/mL) and amphotericin b (Sigma, 2.5 mg/mL) for 4 to 7 days for recovery. After, fish were acclimated to 37°C by incrementally elevating the temperature over the course of 7 days in the presence of these same antibiotics (33°C on day 1, 34°C on day 2, 35°C on day 3, 36°C on day 4, and 37°C on day 5). Following acclimation to 37°C, animals were then held in individual 4L fish tanks containing autoclaved, sterilized fish water supplemented with the same antibiotics outlined above (2 L volume/tank). For animals raised in 4L fish tanks, water changes were performed every 2 to 3 days. Fish were fed two times daily. Once with 81g Medium 1640 (Powder, Gibco), 10g Gemma MiCro 300 (Adult fish food), 5 ml 1% PSG, 50 ml FBS (Gemini), concentrated in 5 ml then diluted into 50 mL of aforementioned antibiotic fish water, and another time with artemia (Brine Shrimp).

### Human Cancer Cell Transplantation into Zebrafish and Single Cell Imaging of Live Zebrafish Using Confocal Imaging

Single cell visualization of engrafted tumors in zebrafish was previously described, and similar procedure was followed for the current experiment^97^. RD FUCCI4 and Rh41 FUCCI4 reporter cell lines were grown to 90% confluence in culture dishes, harvested, and resuspended at 2.5 x 10^5^ cells/µL in Matrigel (Corning). Clondronate liposomes (Clodrosome) were then added to the injection mix to inhibit early macrophage ingestion of engrafted cells over the first 7 days (Encapsula nanoscience, 1 µg/µl). 10 to 20 µl of volume was injected into the retro-orbital space of recipient fish using a 0.36 mm insulin syringe (BD, 5 x 10^5^ cells/eye). Recipient zebrafish were then raised at 37 C in sterilized fish water containing the aforementioned antibiotics. Single-cell resolution imaging of retro-orbitally transplanted cells was performed using confocal microscopic imaging using 20X objective (Zeiss LSM980 inverted microscope), with Zen software platform (Zeiss). Engrafted zebrafish were anesthetized using low-dose 0.01% tricaine (western chemical) and placed into a 22 mm 6 well glass bottom plate (Cellvis, product P06-1.5H-N). Animals were imaged for only 2 to 3 min. FUCCI4 cell cycle imaging studies imaged mTurq2 (excitation = 458 nm, emission=472–508 nm), clover (excitation = 488 nm, emission = 493–586 nm), mKo2 (excitation = 561 nm, emission=575–703 nm) and mMaroon1 (excitation = 633 nm, emission = 599–689 nm). Cell numbers were counted in ImageJ using Particle Analysis plug-in. Total tumor cells (a measure of tumor volume) was determined by quantifying the individual cells at z-stacks and obtaining an average using Image J (NIH).

### Tumor Xenograft Mouse Studies

For all tumor xenograft mouse experiments we adhered to animal protocols approved by each institutions Institute Animal Care and Use Committee (IACUC). For all tumor studies 8-week old male and female NSG mice obtained from Jackson Laboratories had 3.0 x 10^6^ Rh4 cells in 50% Matrigel/DMEM engrafted into their flank. Mice were monitored every other day for tumor growth, and randomly enrolled to treatment when tumors reached 100-150 mm^3^. In experiments examining the effect of A485 alone, mice were randomized to vehicle (10% DMSO + 10% β-hydroxypropylcyclodextran (HPβCD) in water i.p. n = 7) or A485 (75 mg/kg daily i.p. n = 7). In experiments examining the effect of A485 and LG1-34 in combination, mice were randomized to vehicle (10% DMSO + 10% HPβCD in water i.p., 2% DMSO + 1% Carboxymethyl cellulose (CMC) in water p.o. daily n = 8), A485 (75 mg/kg daily i.p. daily n = 8), LG1-34 (50 mg/kg p.o. daily n = 8), or combination A485 + LG1-34 (75 mg/kg A485 i.p, 50 mg/kg p.o daily n = 7). Tumor growth was measured using calipers and mice were weighed every two days. Animals were euthanized in accordance with institutional guidelines when tumors reached 2000 mm^3^ in the A485 alone experiment, or at 1500 mm^3^ in the combination A485 and LG1-34 experiment. Tumor sizes and growth trajectory were compared through day 25 of treatment (A485 alone experiment) or day 24 of treatment (combination A485 and LG1-34 experiment) by fitting growth curves to an exponential growth model, and then comparing growth rate constants using an extra sum-of-squares F-test.

For each mouse xenograft study we performed a concurrent study to assess the effects of each compound on gene and protein expression. In the A485 only experiment 6 mice were xenografted with 3.0 x 10^6^ Rh4 cells as previously. When tumors reached 100-150 mm^3^ three additional mice were treated with vehicle or A485 (75 mg/kg i.p.) for 10 days, prior to euthanization and tumor extraction for IHC. In studies investigating the combination of A485 and LG1-34, 12 additional mice were engrafted in the flank with 3.0 x 10^6^ Rh4 cells and treated with vehicle (n = 3), A485 (75 mg/kg i.p. n = 3), LG1-34 (50 mg/kg p.o. n = 3), or A485 + LG1-34 (75 mg/kg i.p. A485, 50 mg/kg p.o. LG1-34 n = 3) for 7 days. Animals were then euthanized, tumors were extracted, and divided for IHC or RNA-seq analysis.

### Pharmacokinetics Analysis

For pharmacokinetic profiling, CD-1 mice (n = 3 per group) were administered a single dose of LG1-34 either intravenously (IV) via the tail vein at 2 mg/kg or orally (PO) via gavage at 8 mg/kg. Blood samples were collected at designated intervals [e.g., 0.083, 0.25, 0.5, 1, 2, 4, 8, and 24 h] post-dose. Plasma was isolated, and LG1-34 concentrations were quantified using liquid chromatography-tandem mass spectrometry (LC-MS/MS) by Pharmaron. Pharmacokinetic parameters, including area under the plasma concentration-time curve (AUC0-∞), half-life (t1/2), clearance (CL), and volume of distribution (Vss), were calculated using non-compartmental analysis (NCA). Absolute oral bioavailability (F) was determined as follows:

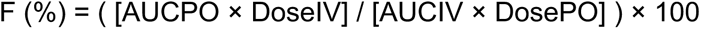

### ChIPseq

Chromatin immunoprecipitation sequencing was performed as previously described^17,82^. Briefly, for each ChIP, 10 μg of antibody was added to 3 ml of sonicated nuclear extract, derived from 50×10^6^ fixed cells. We used the following antibodies for ChIP (IgG, H3K27ac, c-MYC, MYCN, SIX1, SOX8, ZEB2, TCF12, MYOD1) (RRIDs:AB_2118291, AB_1903938, AB_2536872, AB_10988426, AB_669818, AB_1944273, AB_1079991, AB_2890928, AB_1848334).

### CUT&RUN Sequencing

CUT&RUN sequencing was performed by standard methodology and using reagents from Epicypher Inc. H3K27Ac (Millipore # MABE647) and IgG (Epicypher #13-0042) antibodies were used. Briefly, 500,000 live cells per sample were permeabilized and processed by CUT&RUN according to the CUTANA CUT&RUN protocol. Samples were internally controlled by spiking in exogenous *E.coli* DNA, added according to the input cell number, as per the manufacturer’s protocol. DNA was quantified using the Quant-iT PicoGreen ds DNA assay (ThermoFisher). Libraries were prepared with HyperPrep Library Preparation Kit (Roche PN 07962363001) with modified PCR conditions: Step 1 98C for 45s, Step 2 98C for 15s, Step 3 60C for 10s, Step 4 72C for 1min, Repeat steps 2-4 12 times for > 10 ng input, 13 cycles for input between 5 and 10 ng, and 15 cycles for input < 5 ng), Step 4 72C for 1min. Libraries were analyzed for insert size distribution, then libraries were sequenced on a NovaSeq 6000 with 10 million paired-end 75-bp reads per sample.

### Non-Cell-Number-Normalized ChIP-seq/CUT&RUN Analysis

Raw ChIP-Seq/CUT&RUN reads were aligned to the hg19 version of the human reference genome using bowtie v1.2.2^98^ with parameters -k 2 -m 2 –best and -l set to the read length. Paired-end reads were aligned in single-end mode as above and per-read alignments were merged. Coverage tracks were generated from aligned reads using MACS v1.4.1^99^ with parameters -w -S –space=50 –nomodel –shiftsize=200, normalized by the millions of mapped reads, converted into bigwig format using wigToBigWig v4, and visualized in the Integrative Genomics Viewer v2.19.4^100^. Genome browser coverage snapshots at A485+LG1-34-sensitive genes were generated in regions 10kb upstream from the transcription start site to 10kb downstream the transcription end site of the longest isoform of the gene with trackplot (https://github.com/PoisonAlien/trackplot/) atop bwtool (https://github.com/CRG-Barcelona/bwtool) v1.0 with parameters binsize=2, y_min = 0, groupAutoScale = FALSE, collapse_txs = FALSE. Peaks of transcription factors were identified using MACS v1.4.1 using input control and -p 1e-9.

Super-enhancers were identified from aligned H3K27ac and input reads and peaks identified using them as previously^17,101^. Briefly, peaks were identified twice using MACS v1.4.2 with input control and parameters -p 1e-9 –keep-dup=auto and MACS v1.4.2 with input control and parameters -p 1e-9 –keep-dup=all; the collapsed union of these output peaks were used as input into ROSE with parameters -s 12500 -t 2000, which set stitching size to 12.5kb and exclude from stitching peaks fully contained within a 2kb region centered on an annotated transcription start site. Stitched enhancer outputs from ROSE were assigned to the single expressed transcript whose transcription start site was nearest the center of the stitched enhancer after promoter-excluded stitching. Expressed genes were determined for this purpose as being in the top 2/3 when ranking by H3K27ac promoter coverage, identified using bamToGFF [https://github.com/BradnerLab/pipeline/blob/master/bamToGFF.py] -m 1 -r -d -e 200.

### ATAC-seq Analysis

Raw ATAC-Seq reads were aligned to the hg19 revision of the human reference genome using bowtie v1.2.2 in single-end mode with parameters -k 2 -m 2 –best and -l set to the read length. Coverage tracks were generated from aligned reads using MACS v1.4.1 with parameters -w -S – space=50 –nomodel –shiftsize=200, normalized by the millions of mapped reads, converted into bigwig format using wigToBigWig v4, and visualized in the Integrative Genomics Viewer v2.19.4.

### Cell-Number-Normalized CUT&RUN Analysis

H3K27ac CUT&RUN data from Rh4 cells treated with small molecules were aligned and BAMs were normalized using PerCell (https://github.com/lextallan/PerCell/) with default parameters, i.e. --skip_trimming true, --experimental human, --spikein ecoli, --macs2_cutoff 1.301 --skip_downsample false, --skip_motif true, --skip_idr true, --skip_consensus true, -override_spikeinfail false, --skip_bamCoverage false and the spike-in genome set to Escherichia_coli_K_12_MG1655/. Downsampled BAMs were used for downstream coverage analyses.

### Omic coverage analysis

For promoter heatmap, boxplot, and scatterplot analysis, 4kb regions centered on the transcription start sites of the same genes used for RNA-seq analysis were defined. For heatmap analysis specifically of EYA2 and/or SIX1 significant binding sites, peaks of these two factors were collapsed using bedtools (v2.31.0)^102^ merge, filtered against the ENCODE ignore-list (https://www.encodeproject.org/files/ENCFF001TDO/) using bedtools intersect -v, and converted into 4kb windows centered on the middle of collapsed peaks. For enhancer heatmap, boxplot, and scatterplot analysis, the collapsed union of the five non-MYC transcription factor ChIP-Seq peak sets identified as above within one cell line was used. Collapsed regions were defined using bedtools merge, and regions that overlapped ENCODE ignore-list regions were filtered using bedtools intersect -v, and surviving regions that overlapped a promoter were filtered out to minimize ambiguity. Surviving regions were converted into 4kb windows centered on the middle of collapsed peaks. To find enhancers assignable to specific genes, we used bedtools (v2.31.0) closest -t first and the single-bp transcription start site from each transcript.

Using bamToGFF, reads-per-million-normalized (-r) read coverage was quantified in in 100 bins (-m 100) using -f 1 and -e 200, separately for promoters and each cell line’s enhancers in heatmap analysis. Regions were ordered by the row sum of coverage of the specified ranking factor in each figure, which was calculated from the resulting bamToGFF matrix and plotted in R using heatmap.3 (https://gist.github.com/nachocab/3853004). Metagenes were constructed using the column means from each bamToGFF-compueted matrix.

Using bamToGFF, reads-per-million-normalized (-r) read coverage was quantified in in 1 bin (-m 1) using -f 1 and -e 200, separately for promoters and each cell line’s enhancers in scatterplot and boxplot analysis.

### RNA-seq Analysis

Anonymized human RMS tumor data was extracted from SRA: SRP048220. Primary tumor RNA-seq data was aligned to hg38 and expression TPM data was produced using the GTEx pipeline (https://github.com/broadinstitute/gtex-pipeline/tree/master/rnaseq), as in^103^.

RNA-seq data analyzing the effects of A485 on cell lines *in vitro* were initially analyzed as previously^104^. Raw reads from cell lines were aligned to the hg19 revision of the human reference genome to which the sequences of the ERCC spike-in probes were added as pseudochromosomes using hisat v2.1.0^105^. A gene set comprising the RefSeq genes downloaded 7/5/2017 to which the coordinates of the ERCC spike-in probes was used for per-gene quantification with htseq-count^106^ with parameters -i gene_id –stranded=reverse -m intersection-strict. For statistical differential expression analysis, counts were used as input for DEseq2^66^ with default parameters for all options except using the spike-in probe values as the controlGenes in estimateSizeFactors. The threshold used for determine significance were adjusted p value <0.05 and log_2_fold change >|0.5|. Genes were integrated from all four cell lines, and those identified as downregulated in >3/4 cell lines were used as input to StringDB analyses. In parallel, on a per-cell-line basis, DEseq2-normalized log_2_ fold-changes were identified for all CRC genes, and compared against the top 10% of genes on a per-cell-line-basis defined by DEseq2-defined basemean values.

RNA-seq data analyzing the effects of various treatments on xenografted Rh4 cells were first aligned to the mm10 revision of the mouse reference genome to remove contaminating mouse reads using hisat v2.1.0 in paired-end mode. Non-mouse-aligned reads were then aligned to the hg19 revision of the human reference genome to which the sequences of the ERCC spike-in probes were added as pseudochromosomes using hisat v2.1.0. A gene set comprising the RefSeq genes downloaded 7/5/2017 to which the coordinates of the ERCC spike-in probes was used for per-gene quantification with htseq-count with parameters -i gene_id –stranded=reverse -m intersection-strict. For pair-wise differential expression analysis, counts were used as input for DEseq2 with default parameters for all options except using the spike-in probe values as the controlGenes in estimateSizeFactors. Significantly differentially expressed genes had absolute value log2 fold-changes > 0.5 and adjusted p value <0.05 between conditions. For four-condition display, counts were converted to transcripts per million using the standard strategy:

normterm = sum of (readcount * readlength/exonlength) across all genes.

TPM = readcount * readlength/exonlength. * 1e6/normterm.

Per-gene total exon sizes were generated by collapsing all exons of each isoform of each gene into a single set of regions using bedtools^107^ merge, then quantifying the numbers of unique base pairs in these collapsed exons.

### Single Cell RNAseq Analysis

Single cell RNA sequencing (scRNAseq) data from rhabdomyosarcoma tumors was obtained as fully processed Seurat object from FigShare https://figshare.com/projects/RMS_consensus_analysis/194417. The original analysis of this data was performed as described by the original authors^50^. For the current analysis we examined scRNAseq data for expression of EYA1-4 transcripts. No further raw data processing was performed.

### IHC Methods

Tumors were dissected, fixed in 10% Neutral Buffered Formalin, and processed using the HistoCore PEGASUS Tissue Processor (Leica Biosystems, Deer Park, IL). Processed tissues were embedded in paraffin and sectioned at a thickness of 4 µm using a HistoCore AUTOCUT fully automated rotary microtome (Leica Biosystems, Deer Park, IL). Tumor sections were stained with hematoxylin and eosin (H&E) using the HistoCore SPECTRA ST Stainer (Leica Biosystems, Deer Park, IL). Simplex immunohistochemistry was performed to detect cleaved caspase 3, c-MYC, Ki67, and H3K27ac. Pertinent details are summarized in **Supplementary Table 8**.

### Gene Set Enrichment Analysis (GSEA)

GSEA (RRID:SCR_003199) was performed using whole transcriptomes recovered from RD, JR, Rh4 and Rh28 cells treated with A485 at the cell line-specific day 3 IC_50_ value of A485, compared with DMSO, using the MSIGDB Hallmarks dataset, specifically focusing on the Hallmarks “Myogenesis” and “MYC Targets V1.” GSEA was also performed on RNA-seq data from Rh4 xenograft tumors treated for 7 days with the A485 + LG1-34 combination, compared with vehicle-treated tumors. Genes were pre-ranked in descending order by the DESeq2 Wald statistic and analyzed against the MSigDB Hallmarks dataset, matched by gene symbol, with minGSSize = 15 and maxGSSize = 500. Gene sets with a BH-adjusted p-value < 0.05 were considered significantly enriched.

### STRING Database Analysis

Genes downregulated by 2h of treatment with A485 at the cell line-specific day 3 IC_50_ value, compared to DMSO were compared across four cell lines (RD, Rh4, Rh28, JR). Gene dependencies found downregulated (adjusted p<0.05) in >3/4 cell lines (n=61) were used as input into the String database (RRID:SCR_005223;^108^). Network edges reflect evidence of interactions. Color was used to indicate CRC member (red) or not (blue), or alternatively, membership in the Gene Ontology dataset 006357, “Regulation of Transcription by RNA polymerase II” (green) or not (grey).

## Data and Code Availability

RNAseq and ChIP-Seq data generated in this study have been deposited in the Gene Expression Omnibus (GEO) database under SuperSeries accession number GSE307452. Eya2 with LG1-34 structures are deposited in the Protein Data Bank under the ID 9Y64. No custom code was developed for use in the manuscript.

## Acknowledgements

This work was supported by NIH grants T32-GM008497 (A.L.G.), T32-GM136444 (A.L.G.), R01-CA275187 (H.L.F., K.B.A., A.D.D.), R01-CA301267 (H.L.F., R.Z.), K08-CA245251 (A.D.D), R37-CA286444 (A.D.D), R01-CA296556 (A.D.D, B.J.A, J.Q), and P30-CA021765 (to A.D.D., B.J.A), R35-GM145289 (R.Z), R41-CA291231-01A1 (H.L.F, R.Z, X.W), R01-CA289761 (A.L.H), R01-CA276116 (D.M.L.), R01-CA269213 (D.M.L.), R24-OD031955 (D.M.L.), R35-CA283977 (K.S), T32-CA190216 (L.W), F31-CA275314-01A1 (A.W), K00-CA245552 (S.R.R). B.E.G. was supported through the DOD’s Convergent Science Virtual Cancer Center (W81XWH-21-1-0298) This work was supported by the American Lebanese Syrian Associated Charities, and the St. Jude Children’s Research Hospital Transcription Collaborative. This work was also supported by the Pediatric Cancer Dependencies Accelerator of the Broad Institute, Dana-Farber Cancer Institute, and St. Jude Children’s Research Hospital: peddep.org. B.E.G. A.D.D, J.Q., and N.A.M.S are supported by the Alex’s Lemonade Stand Foundation. A.D.D. and N.A.M.S. were supported by the Rally Foundation for Childhood Cancer Research. A.L.G. was supported and would like to acknowledge the Victor W. Bolie and Earleen D. Bolie Graduate Scholarship Fund. A.D.D. and J.Q. were supported by the Curing Kids Cancer Foundation. A.D.D was supported by CureSearch for Children’s Cancer, the V Foundation for Cancer Research and the Hyundai Hope on Wheels Foundation. K.S. and A.D.D. were supported by Team Sciarappa Strong. B.E.G. is supported by Reign in Sarcoma and the B+ Foundation. Spark and ABNexus helped provide support to R.Z, H.L.F, and X.W. We would also like to acknowledge the Walter and Marina Bornhorst family for their contributions to this work (N.V.D, A.L.H, G.K, J.S.B, J.N.R, F.V, K.S, A.D.D).

The authors would also like to acknowledge Meifen Lu, IHC Team, Comparative Pathology Core, Department of Pathology at St. Jude Children’s Research Hospital for her contributions to the *in vivo* IHC staining slide and protocol generation, and Dr. Todd R. Golub (Broad Institute of MIT and Harvard) for his early contributions to the Pediatric Dependency Map. The authors thank investigators and laboratories that provided RMS cell lines for study, including Drs Beat Schafer (University Hospital Zurich), David Malkin (Sickkids Hospital Research Institute) Paul Jedlicka (University of Colorado, Anschutz Medical Campus), and the Children’s Oncology Group Cell Line Repository (www.cogcells.org). The authors would also like to acknowledge the X-ray Crystallography Core Facility at the University of Colorado Anschutz Medical Campus which is supported by the University of Colorado Cancer Center Support Grant (NIH P30CA046934). Beamline 8.2.1 of the Advanced Light Source, a DOE Office of Science User Facility under Contract No. DE-AC02-05CH11231, and is supported in part by the ALS-ENABLE program funded by the National Institutes of Health, National Institute of General Medical Sciences, grant P30 GM124169-01.

