## Supplementary Figures for "Synergistic targeting of EP300/CBP and EYA co-activators collapses the rhabdomyosarcoma core regulatory circuit"

Supplementary Information includes 25 Supplementary Figures and 8 Supplementary Tables

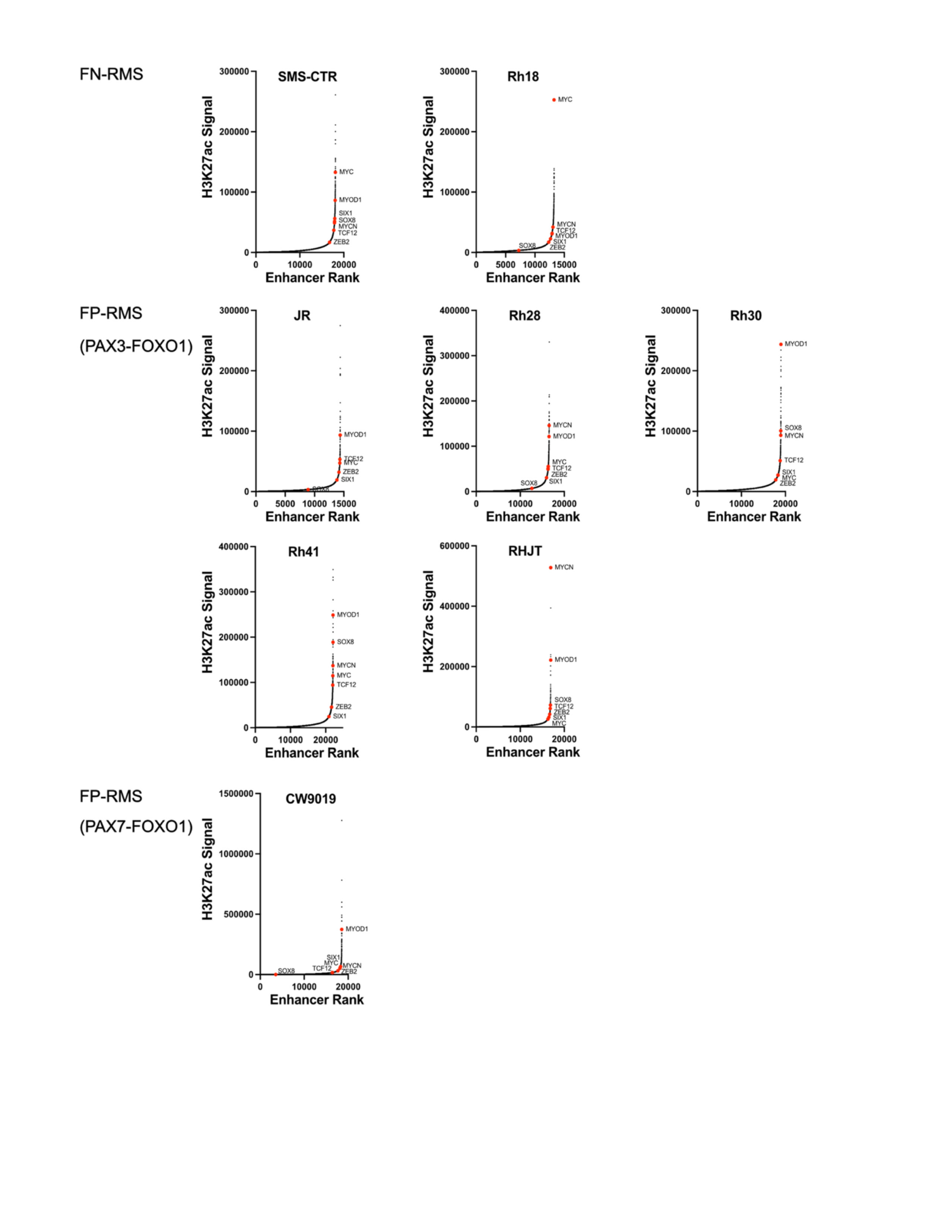

**Supplementary Figure 1. A subset of super-enhancer regulated genes include transcription factors found across 10 RMS cell lines.**

Plots of H3K27ac signal, compared to enhancer rank, in FN-RMS (SMS-CTR and Rh18) and FP-RMS PAX3-

FOXO1 (JR, Rh28, RHJT, Rh30, and Rh41) and FP-RMS PAX7-FOXO1 (CW9019). Highlighted are common

transcription factors identified in >6/9 cell lines displaying super-enhancers.

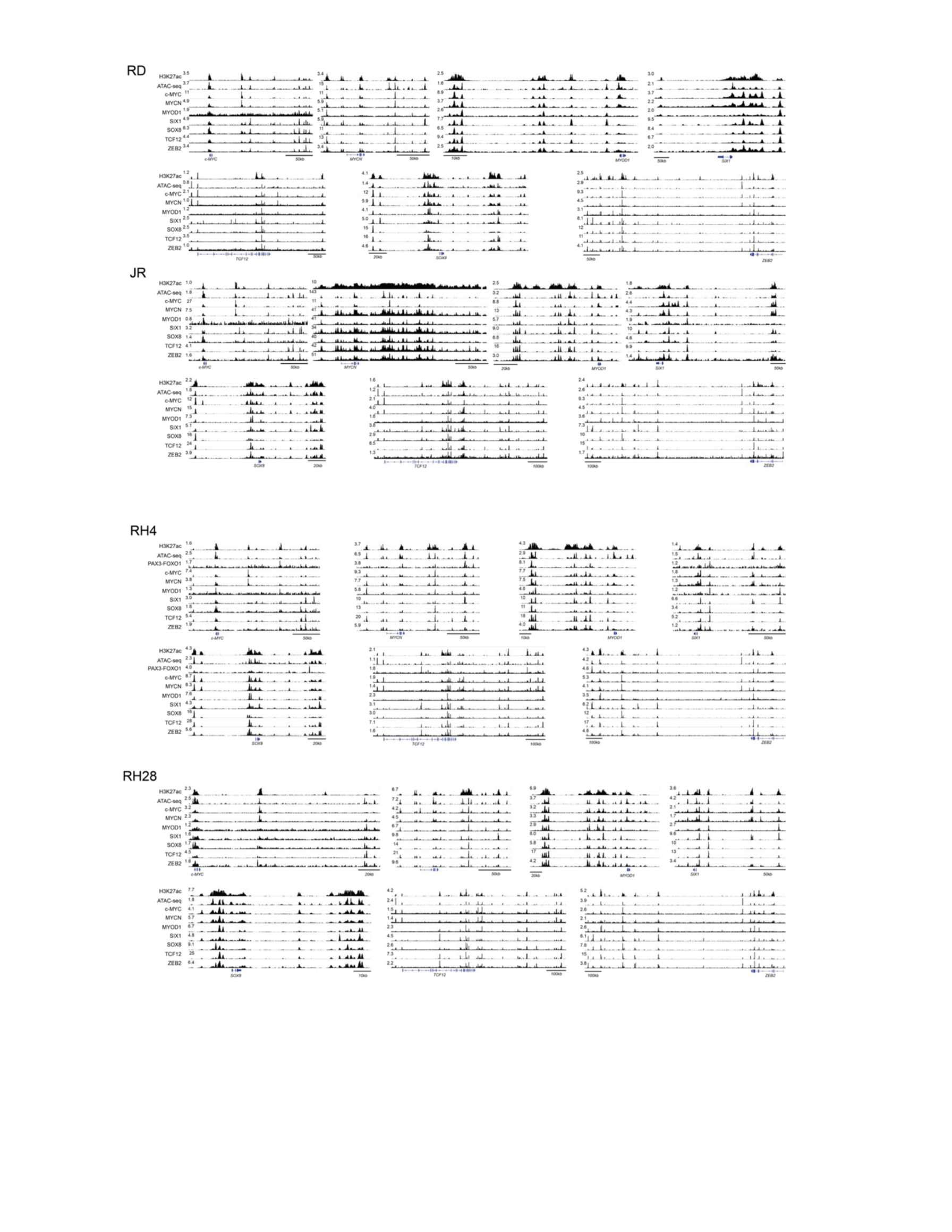

**Supplementary Figure 2. Core regulatory circuit mTFs bind at SEs to regulate their own and each others**

**transcription.**

ChIP-seq gene tracks at all CRC mTF loci in FN-RMS RD cells, and FP-RMS JR, Rh4, and Rh28 cells. Tracks

show co-binding at cis-regulatory elements of c-MYC, MYCN, MYOD1, SIX1, SOX8, TCF12, and ZEB2 at

regions of open chromatin resolved by ATAC-sequencing, and within super-enhancers, resolved by H3K27ac

ChIP-seq.

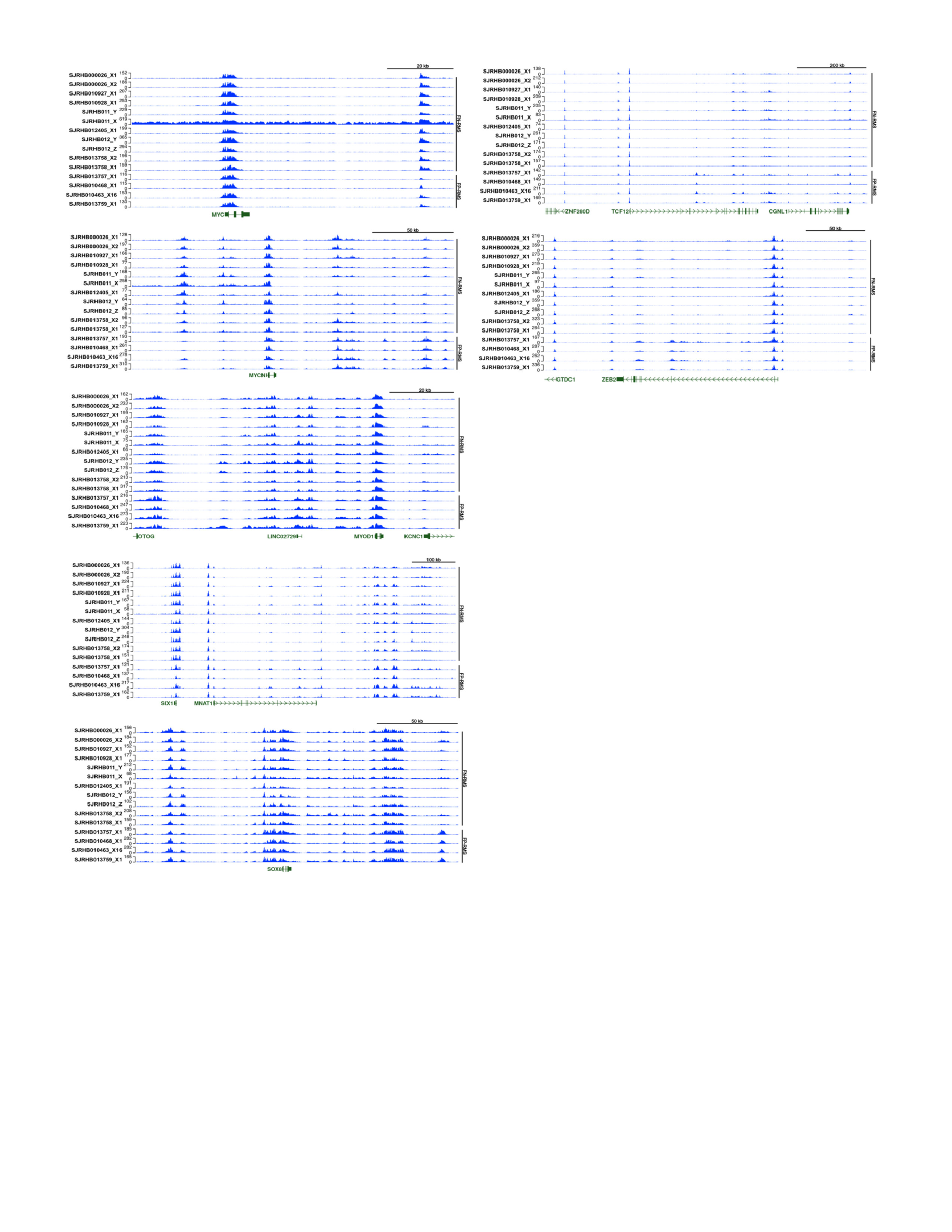

**Supplementary Figure 3. Super-enhancer regulated genes identified in RMS cell lines are also marked by large stretches of H3K27ac in RMS PDXs.**

ChIP-seq gene tracks for H3K27ac at all CRC mTF loci in RMS PDXs. Data obtained from https://www.stjude.cloud/, originally deposited in Stewart, E. et al. Cancer Cell (2018)16. Sample accessions: fusion negative tumors - SJRHB000026_X1_Rec3, SJRHB000026_X2_Rec4, SJRHB010927_X1_Prim, SJRHB010928_X1_Rec1, SJRHB011_Y_Prim, SJRHB011_X_Rec1, SJRHB012405_X1_Rec1,

SJRHB012_Y_Rec1_SiteA, SJRHB012_Z_Rec1_SiteB, SJRHB013758_X2_Rec1, SJRHB013758_X1_Prim. Fusion positive tumors: SJRHB010468_X1_Prim, SJRHB010463_X16_Rec1, SJRHB013757_X1_Prim,SJRHB013759_X1_Rec1.

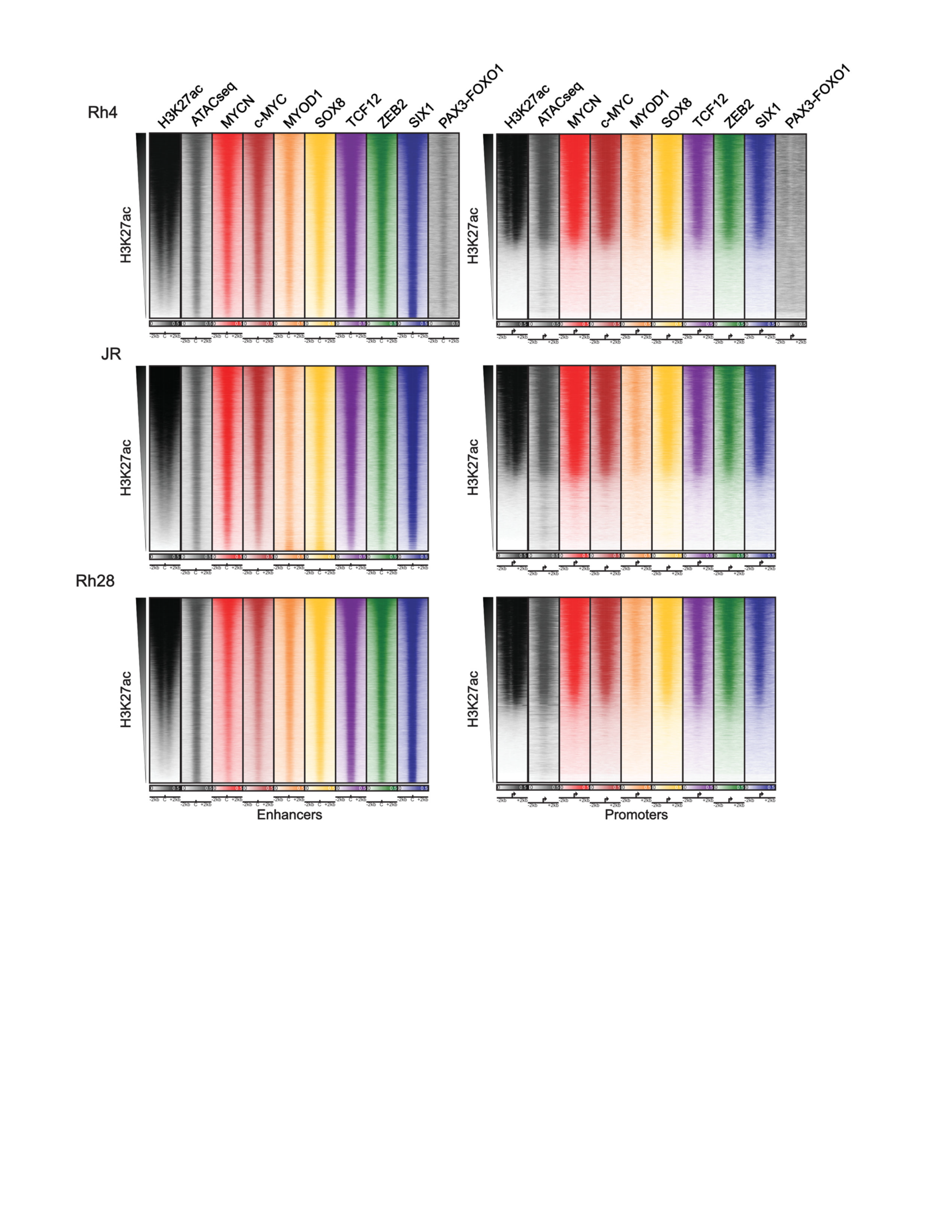

**Supplementary Figure 4. Core regulatory circuit mTFs bind at overlapping sites genome-wide.**

Genome-wide heatmaps ranked by H3K27ac signal at enhancers (left) and promoters (right) for the union of peaks bound by MYOD1, SIX1, SOX8, TCF12 and ZEB2. Binding of c-MYC and MYCN are also shown, and accessible chromatin is shown by ATACseq. Cell lines demonstrated are fusion positive JR, Rh4 and Rh28.

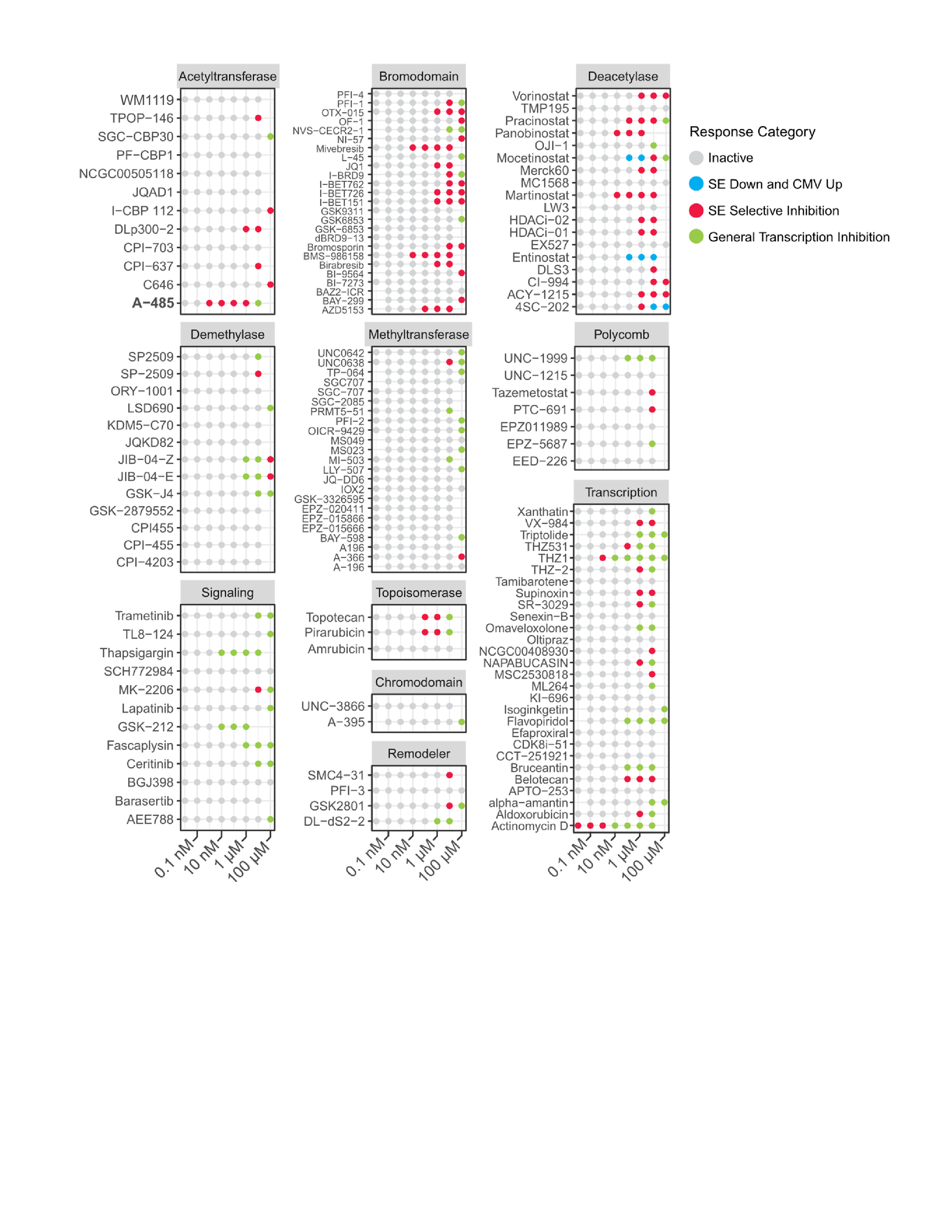

**Supplementary Figure 5. Individual compound results from epigenetic targeted compound screen.**

Individual compound results from super-enhancer-luciferase reporter screen. n=147 total compounds tested.

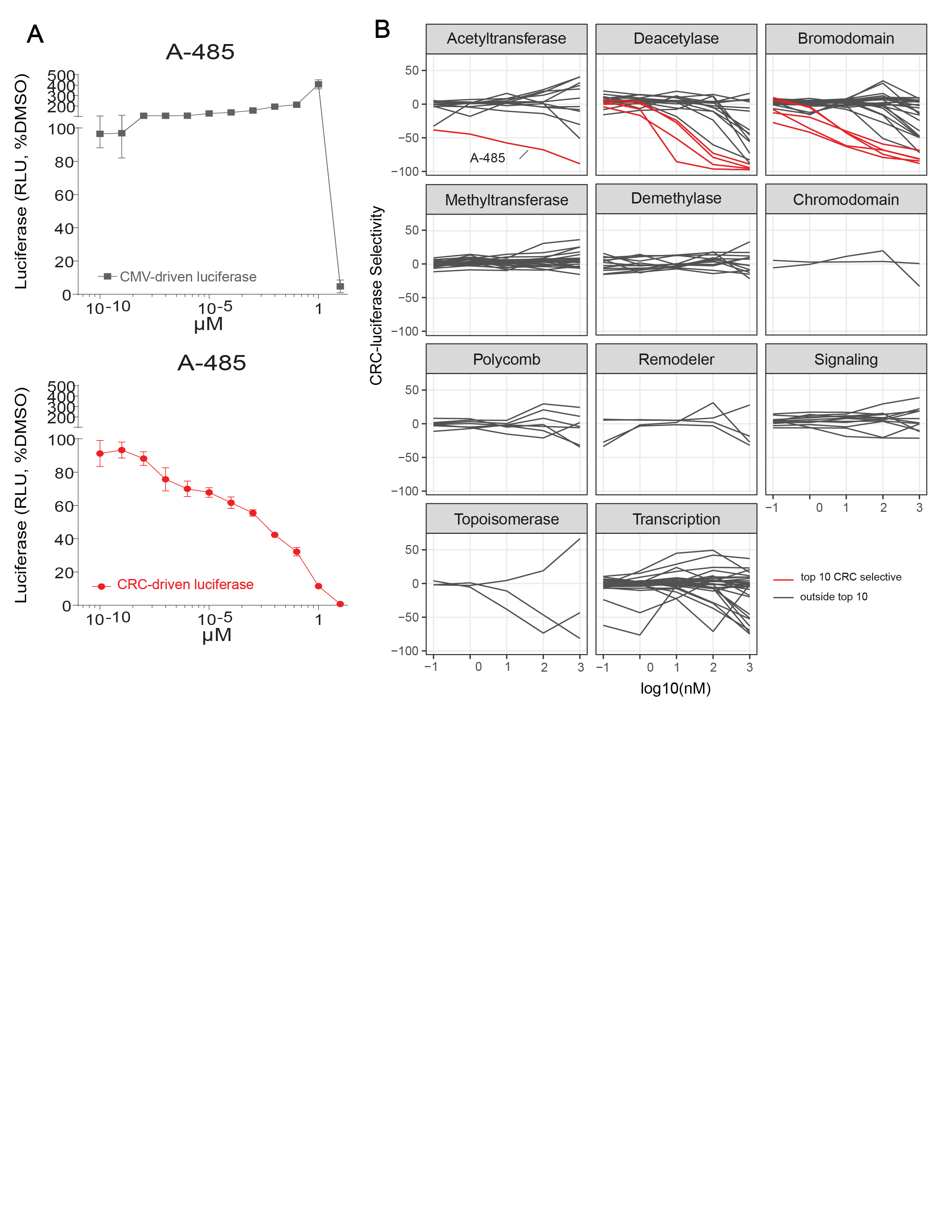

**Supplementary Figure 6. Dose curves for compounds tested in epigenetic-targeted compound screen.**

A. Luciferase signal driven by either a CMV-promoter or driven by a CRC-SE controlled-promoter in Rh4 cells treated with A485 at doses ranging from 0.1 nM to 100 µM. Luciferase signal normalized to the DMSO negative control is plotted in both the CMV-driven luciferase control and the CRC-driven luciferase.
B. CRC-luciferase selectivity in all compounds tested in the drug screen grouped by compound target. The top 10 CRC selective compounds are highlighted in red.

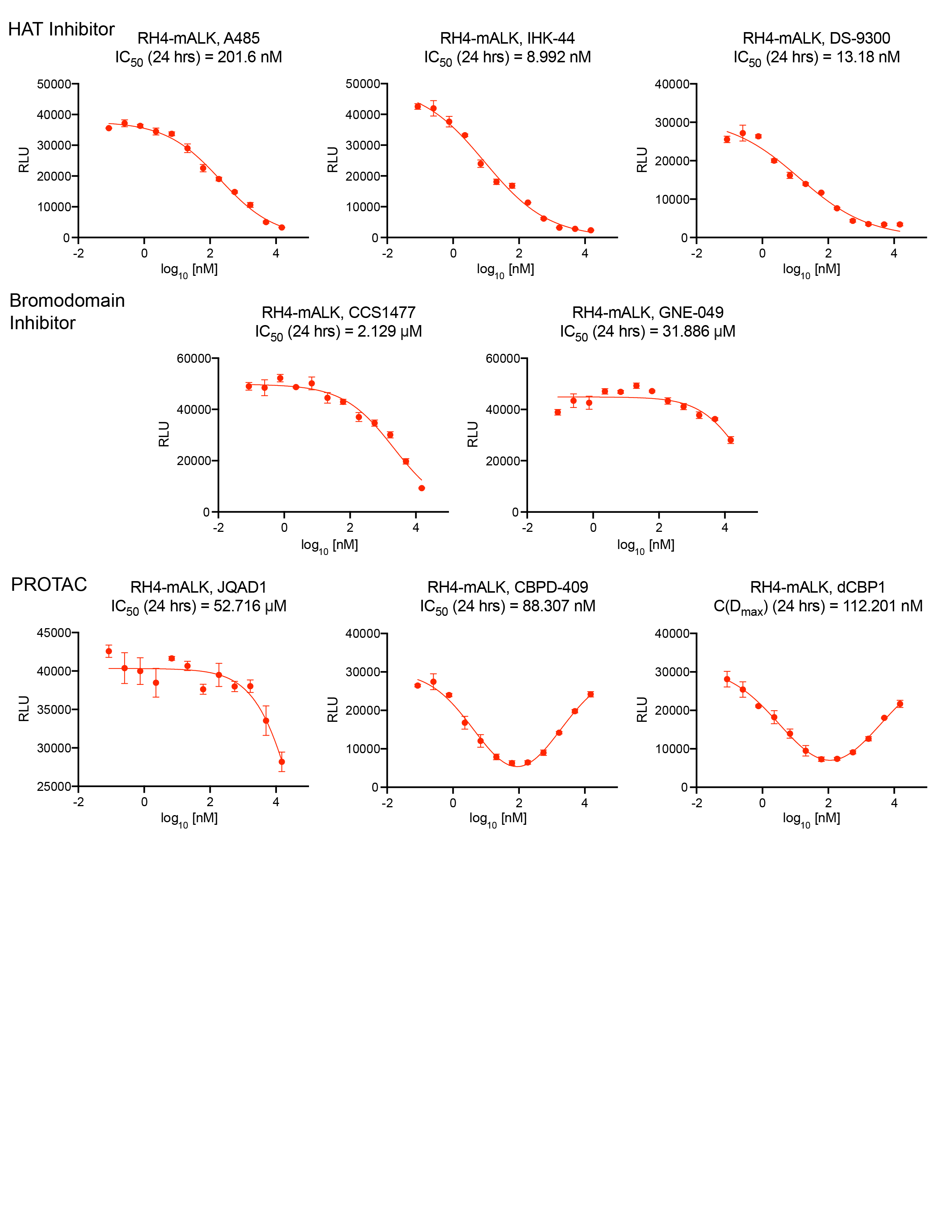

**Supplementary Figure 7. Luciferase signal in Rh4 ALK-SE cells treated with EP300/CBP inhibitors**

Luciferase signal measured in Rh4 ALK-SE-luciferase cells treated with EP300/CBP HAT inhibitors (A485, IHK-44, and DS-9300), CBP/EP300 bromodomain inhibitors (CCS1477 and GNE-049), and CBP/EP300 degraders (JQAD1, CBPD-409, and dCBP1). IC50 values are measured at 24 hours of treatment.

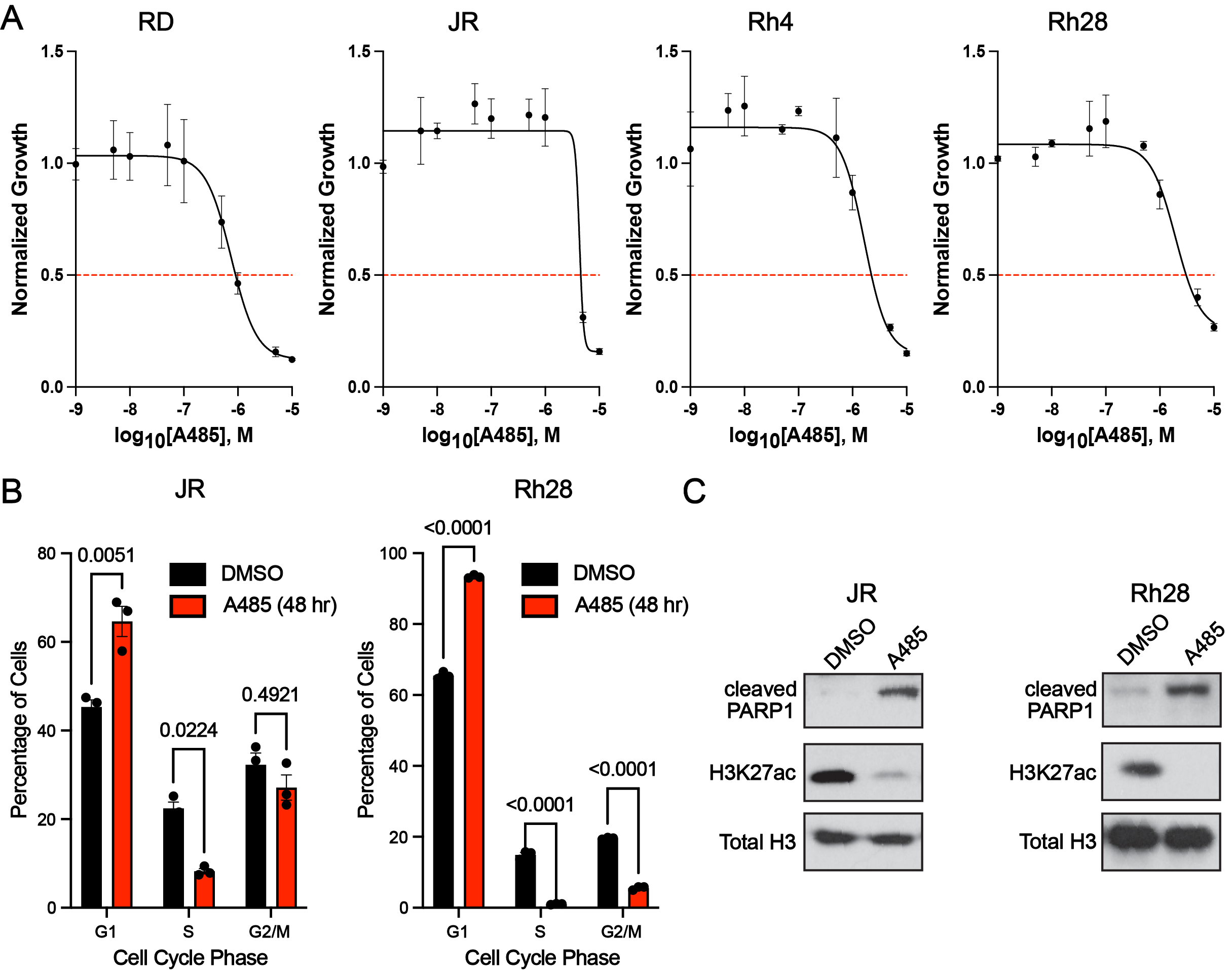

**Supplementary Figure 8. A485 causes cell cycle arrest and apoptosis in RMS cells.**

A. RMS cells were treated with a range of doses of A485 for 7 days, prior to resolution of cell growth by

CellTiter-Glo assay. Data plotted is average of n=3 independent experiments per cell line. Bars = S.E.M.

B. RMS cell lines were treated with the IC50 value of A485 as determined in A, for 48h prior to analysis for cell

cycle by propidium-iodide flow cytometry. Cells were treated with DMSO as a vehicle control. Data shown

represents average of n=3 per treatment.

C. RMS cell lines were treated for 48h with DMSO or A485 at 500nM (RH4), 1 μM (JR), 2 μM (RD) and 3 μM

(RH28), prior to lysis and protein extraction for western blotting. Cleaved PARP1 and H3K27ac are shown, with

total histone H3 shown as a loading control. Data is representative of 3 independent treatments, lysates and

immunoblots for each cell line.

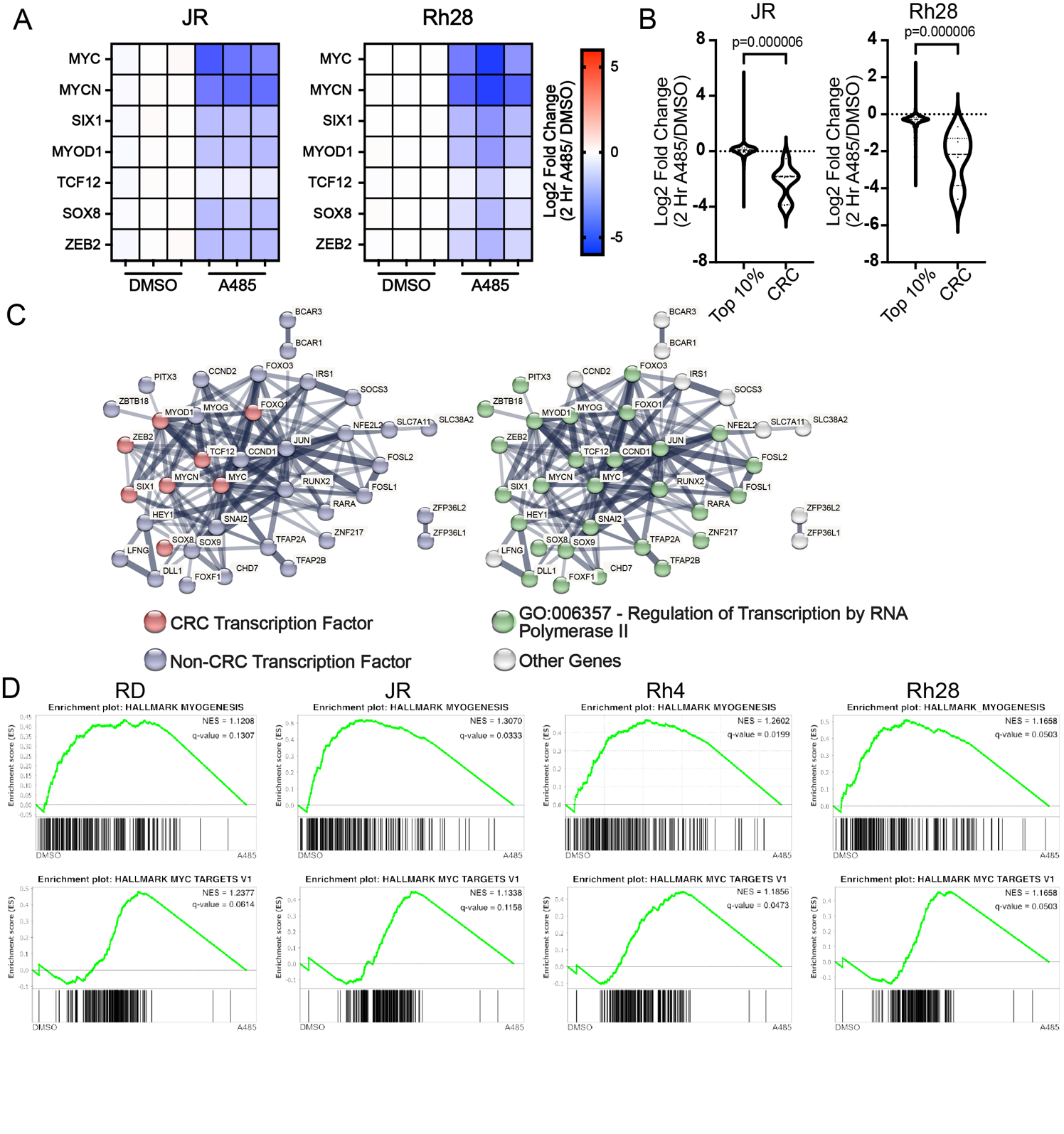

**Supplementary Figure 9. EP300/CBP inhibition reduces CRC expression leading to loss of myogenic**

**and MYC-driven transcriptional hallmarks.**

A. JR and Rh28 cells were treated with 1μM (JR) or 3 µM (Rh28) A485 for 2hr prior to RNA extraction and

external spike-in normalized RNA-seq analysis. Shown are heatmaps demonstrating log2(fold change) at 2hr,

compared with DMSO treatment, in CRC mTF transcript abundance.

B. Analysis of the same RNAseq data from Fig S7A, examining the response of the top 10% highest

expressed genes in vehicle controls after 2h A485 treatment, compared to CRC gene response at the same

time point. Statistical significance was assessed by a two-sample Kolmogorov-Smirnov test with p<0.05

considered significant. n=3 treatments and RNA isolation/sequencing per time point, treatment and cell line.

C. STRING database analysis of genes down-regulated (Log2FC<-0.5 in 3/4 RMS lines) after 2 hours of A485

treatment compared with DMSO treatment, that are also dependencies in at least 1 RMS line. Left: RMS CRC

members are highlighted (red) vs other (blue). Right: Genes that are members of Regulation of Transcription

by RNA Polymerase II (GO:006357) are highlighted in green (right) vs. other (grey).

D. Gene Set Enrichment Analysis in RMS cell lines treated for 2 hours with A485 at 500nM (RH4), 1 μM (JR),

2 μM (RD) and 3 μM (RH28), or DMSO demonstrate an enrichment for gene sets HALLMARK MYOGENESIS,

and HALLMARK MYC TARGETS V1. NES = normalized enrichment score. n=3 treatments and RNA

isolation/sequencing per time point, treatment and cell line, comparing vehicle control against 2 hour A485

treatment.

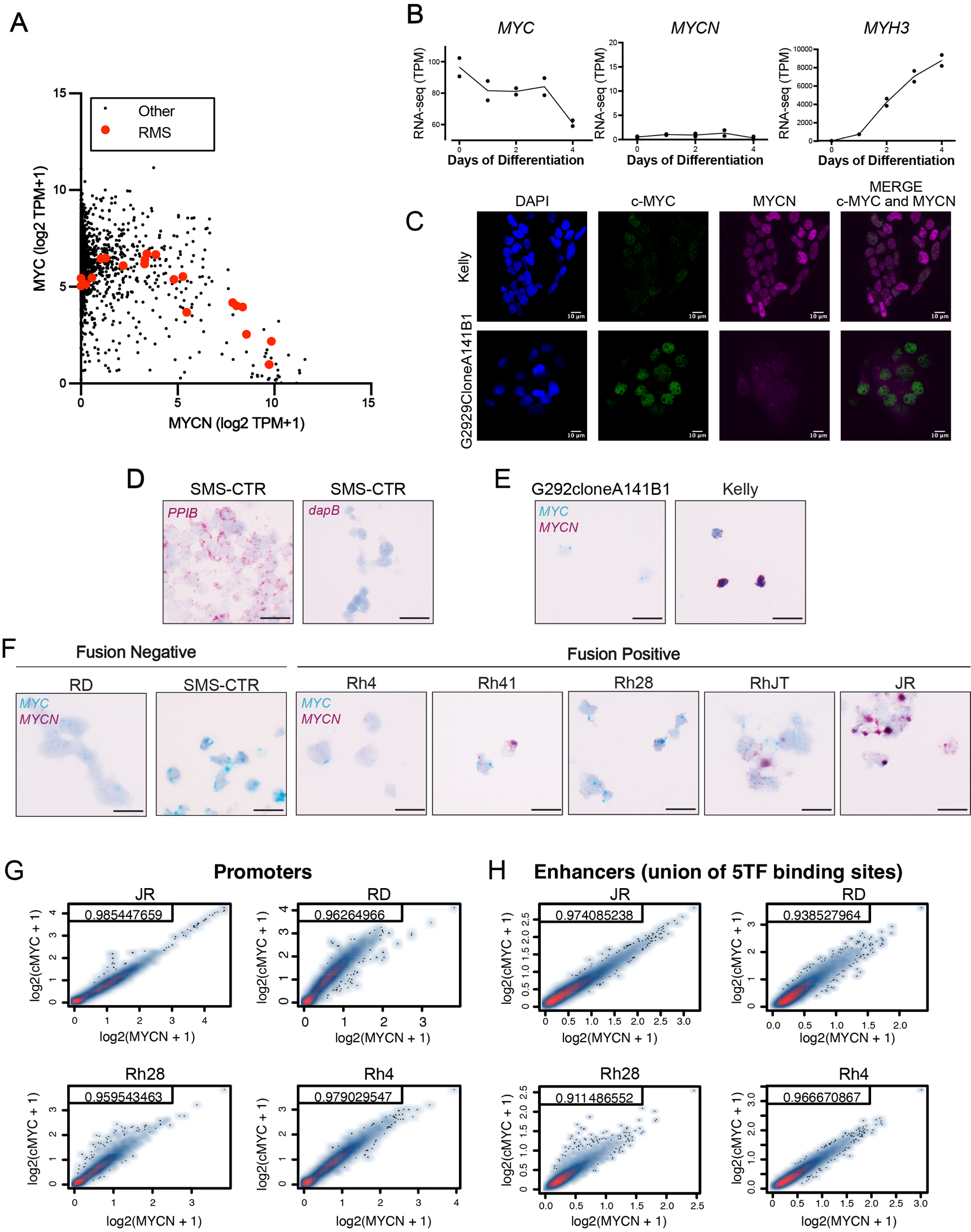

**Supplementary Figure 10. c-MYC and MYCN are both expressed in RMS cells.**

A. Cancer Cell Line Encyclopedia RNAseq expression analysis of c-MYC and MYCN. RMS cell lines display expression of both c-MYC and MYCN, in contrast to the majority of other cell lines. n=29 (RMS), 2274 (other) cell lines. Data retrieved from www.depmap.org, CCLE data release 25Q2.
B. Expression of cMYC and MYCN during staged skeletal muscle development as measured by RNA-seq. Differentiation of human skeletal muscle myoblasts over 4 days is shown by skeletal muscle differentiation marker MYH3. Data retrieved from r2.amc.nl.

C. Immunofluorescent detection of c-MYC and MYCN was performed on a MYCN-amplified neuroblastoma cell line (Kelly) and a c-MYC-amplified osteosarcoma cell line (G292CloneA141B1). c-MYC is stained in green, MYCN in magenta, and DAPI in blue. Images are representative of 3 images and independent staining reactions. Bar = 10 µm.

D. RNAscope for positive control PP1B (pink chromogen) and negative control dapB (pink chromogen) in SMS-CTR cells. Images are representative of 3 images. Bar = 20µm.
E. RNAscope performed for c-MYC (teal chromogen) and MYCN (pink chromogen) in G292cloneA141B1

(osteosarcoma) and Kelly (neuroblastoma) cell lines. Images are representative of 3 images. Bar =20 µm.

F. RNAscope performed for c-MYC and MYCN (pink chromogen) in FN-RMS cell lines (RD and SMS-CTR) and FP-RMS cell lines (Rh4, Rh41, Rh28, RhJT, and JR). Images are representative of 3 images. Bar = 20 µm.

G. Heatmaps generated demonstrating signal from ChIPseq for c-MYC and MYCN in four different RMS cell lines specified to enhancers and promoters. Promoters from all genes genome-wide were queried for average c-MYC and MYCN occupancy.
H. For enhancer analysis, the collapsed union of 5 mTFs, MYOD1, SIX1, SOX8, TCF12, and ZEB2 identified in at least one cell lines were used. Spearman correlation coefficient values are represented in the upper-left corner of each plot and demonstrate the degree in overlap between c-MYC and MYCN binding sites.

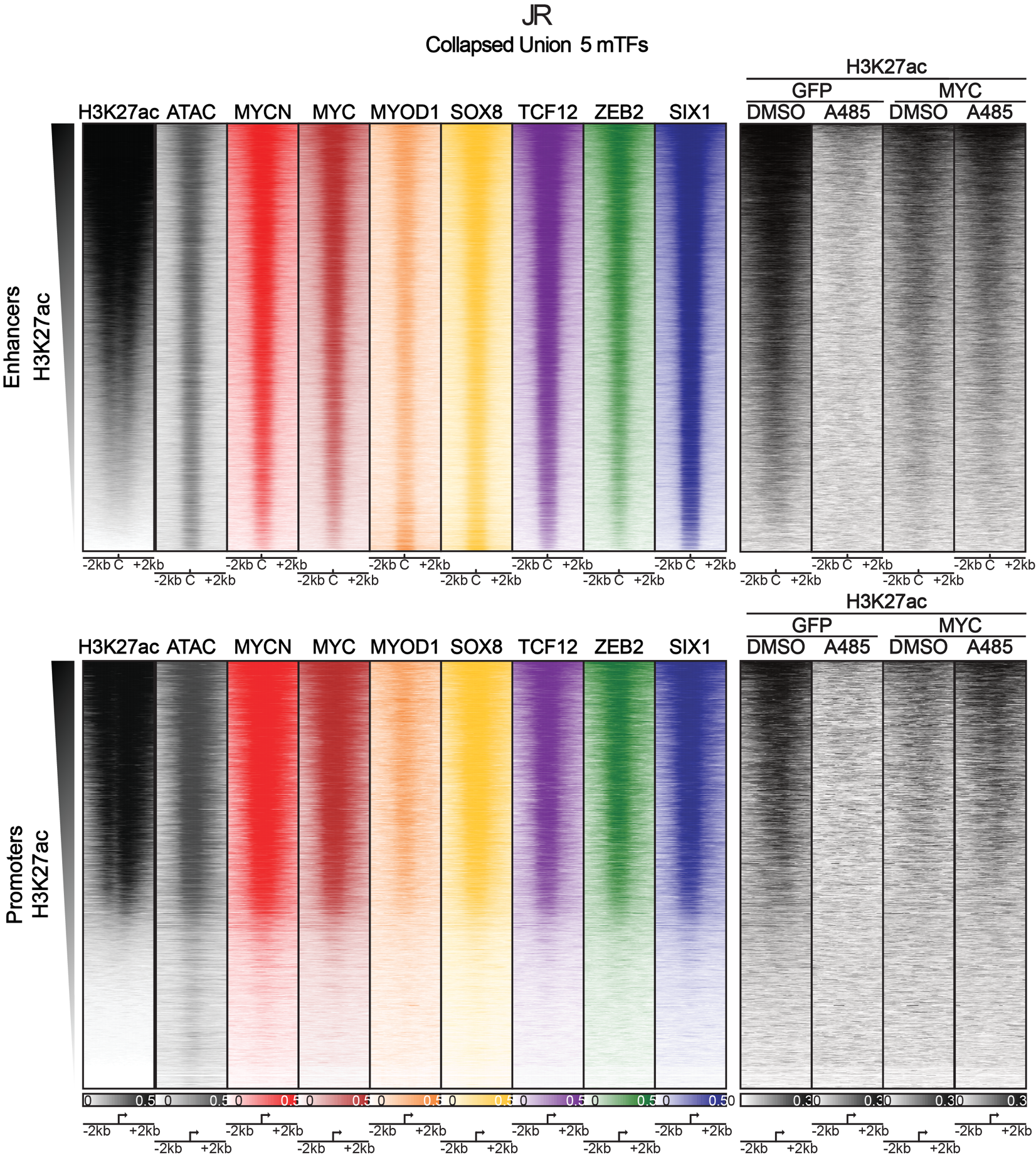

**Supplementary 11. MYC overexpression partially rescues A485-induced loss of H3K27ac.**

A. Reads-per-million-normalized CUT&RUN coverage heatmaps at enhancers (above, defined by master TF

binding peaks) or promoters (below, transcription start sites +/-2kb). GFP overexpression as control for MYC

overexpression. DMSO as control for A485 treatment with 1 µM for 2 hours.

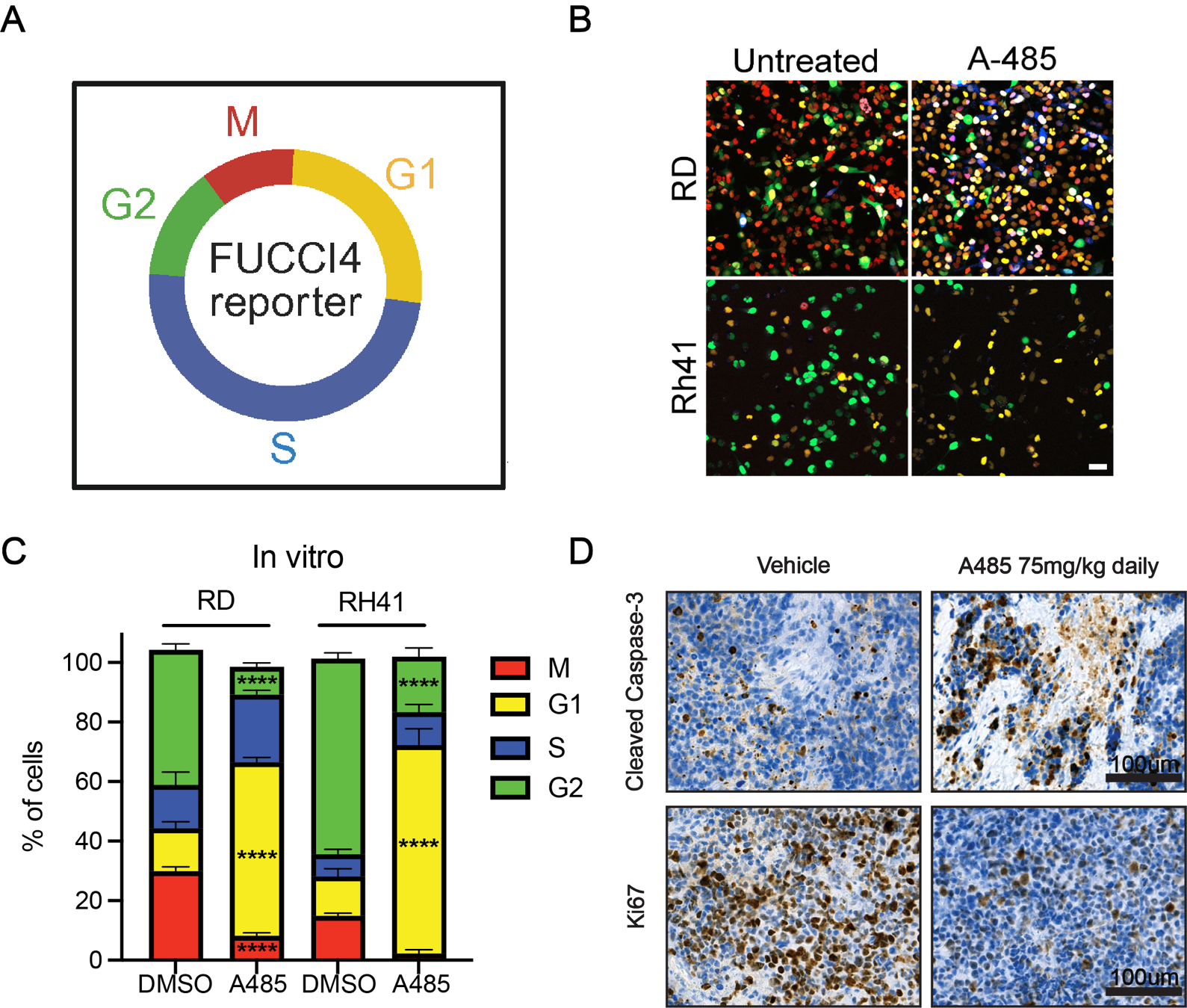

**Supplementary Figure 12. A485 causes RMS cell cycle arrest and cell death.**

A. Diagram showing FUCCI4 reporter system.

B. Fluorescent images showing FUCCI4 reporter system in RD and Rh41 cells treated with DMSO and A485

at 2 µM and 1 µM respectively for 24h. Images are representative of three independent treatments. Bar=20 µm

C. Quantification of cell cycle phases from cells treated in A. n=100 cells per time point analyzed from 3

independent images. **** p<0.0001 by two-way ANOVA comparing across cell cycle phases.

D. Immunohistochemistry for Ki67 and cleaved caspase-3 in Rh4 xenograft tumors. Images are representative

of three separate tumors taken from individual mice. Bar=100 µm.

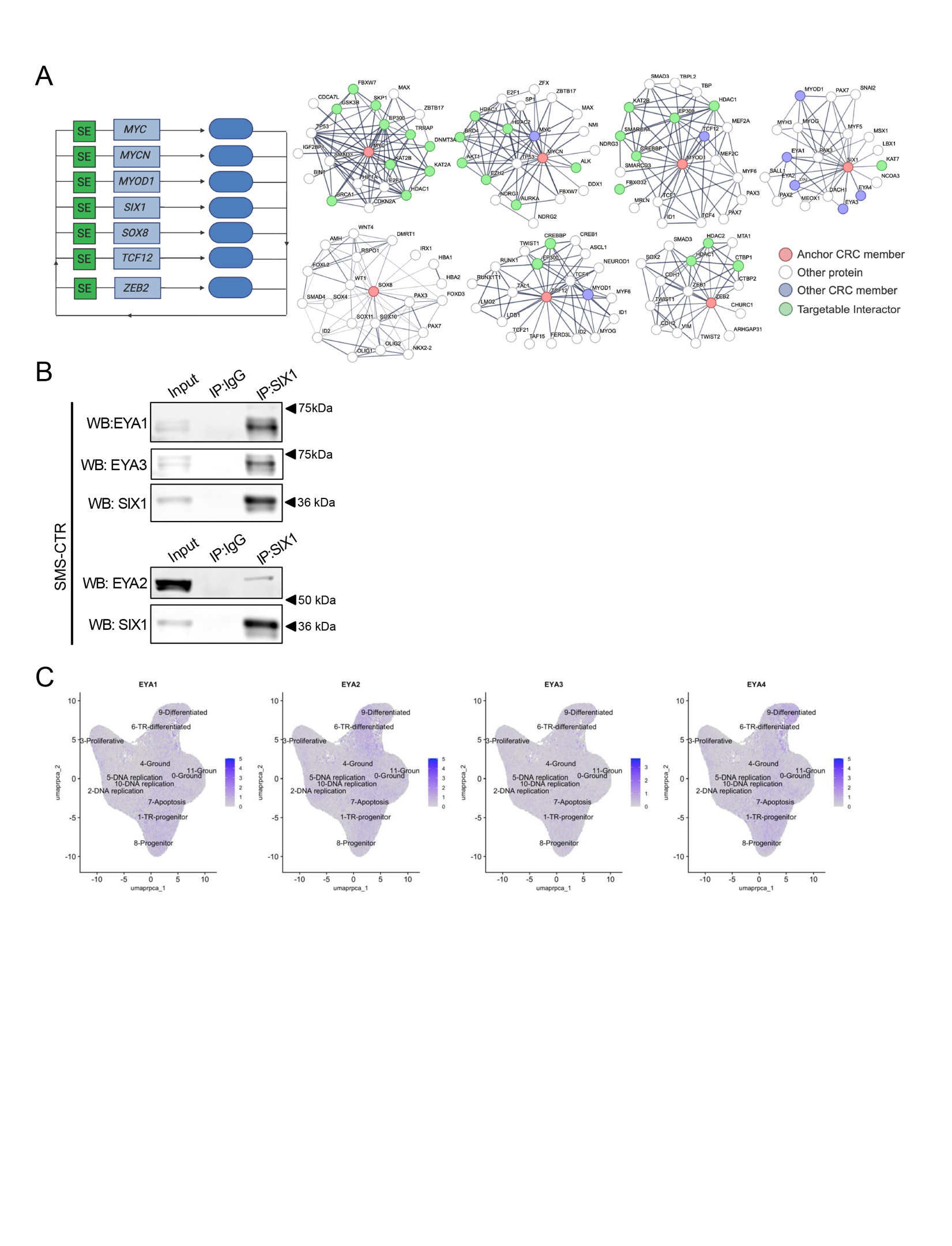

**Supplementary Figure 13. EYA transcriptional co-activators interact with SIX1 and are expressed in primary human RMS.**

A. STRING database analysis of CRC members in the pan-RMS CRC. The anchor CRC member is in red,

targetable proteins are in green, and other CRC members are in blue.

B. Co-immunoprecipitation for SIX1 in SMS-CTR FN-RMS cells followed by western blot analysis where blots

were probed for SIX1, EYA1, EYA2, and EYA3. Data is representative of three independent co-

immunoprecipitation analyses and blots.

C. UMAP plots of integrated cells/nuclei (n = 72 datasets, 107,523 cells), cells are labeled by assigned cell

state. EYA1, EYA2, EYA3, and EYA4 expression are shown. Data retrieved from https://figshare.com/projects/

RMS_consensus_analysis/194417.

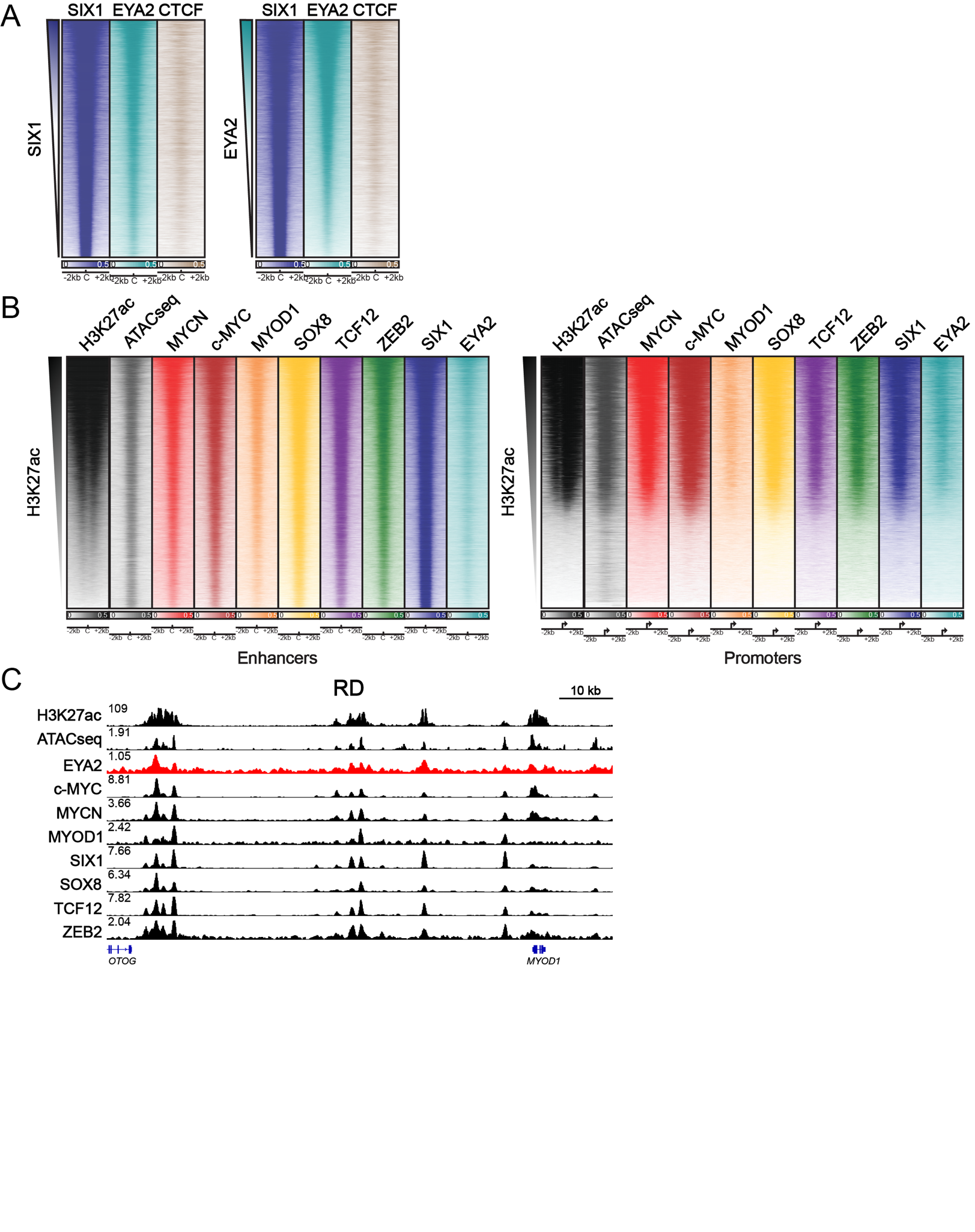

**Supplementary Figure 14. EYA2 is a co-factor associated with the pan-RMS core regulatory circuit.**

A. MACS-derived peaks for either SIX1 or EYA2 were collapsed, and SIX1, EYA2, and CTCF binding were

quantitated at this set of regions. The same set of regions is displayed on the left and right, but are differentially

ranked: left ranks rows by SIX1 signal, right ranks rows by EYA2 signal. High quantitated signal across virtually

all rows indicates binding of SIX1 and EYA2 occurs at the same sites. CTCF is included as negative control

unlikely to bind most activating cis-regulatory elements.

B. Genome-wide heatmaps ranked by H3K27ac signal at enhancers (left) and promoters (right) for the union

of peaks bound by MYOD1, SIX1, SOX8, TCF12, and ZEB2 in RD cells. Binding of c-MYC, MYCN, and EYA2

are also shown, and accessible chromatin is shown by ATACseq. Data is representative of two independent

RMS cell lines.

C ChIP-seq gene tracks at the MYOD1 locus in RD FP-RMS cells, demonstrating co-binding at cis-regulatory

elements of EYA2, c-MYC, MYCN, MYOD1, SIX1, SOX8, TCF12, and ZEB2. Binding occurs in regions of

open chromatin resolved by ATAC-sequencing, and within a super-enhancer, resolved by H3K27ac ChIP-seq.

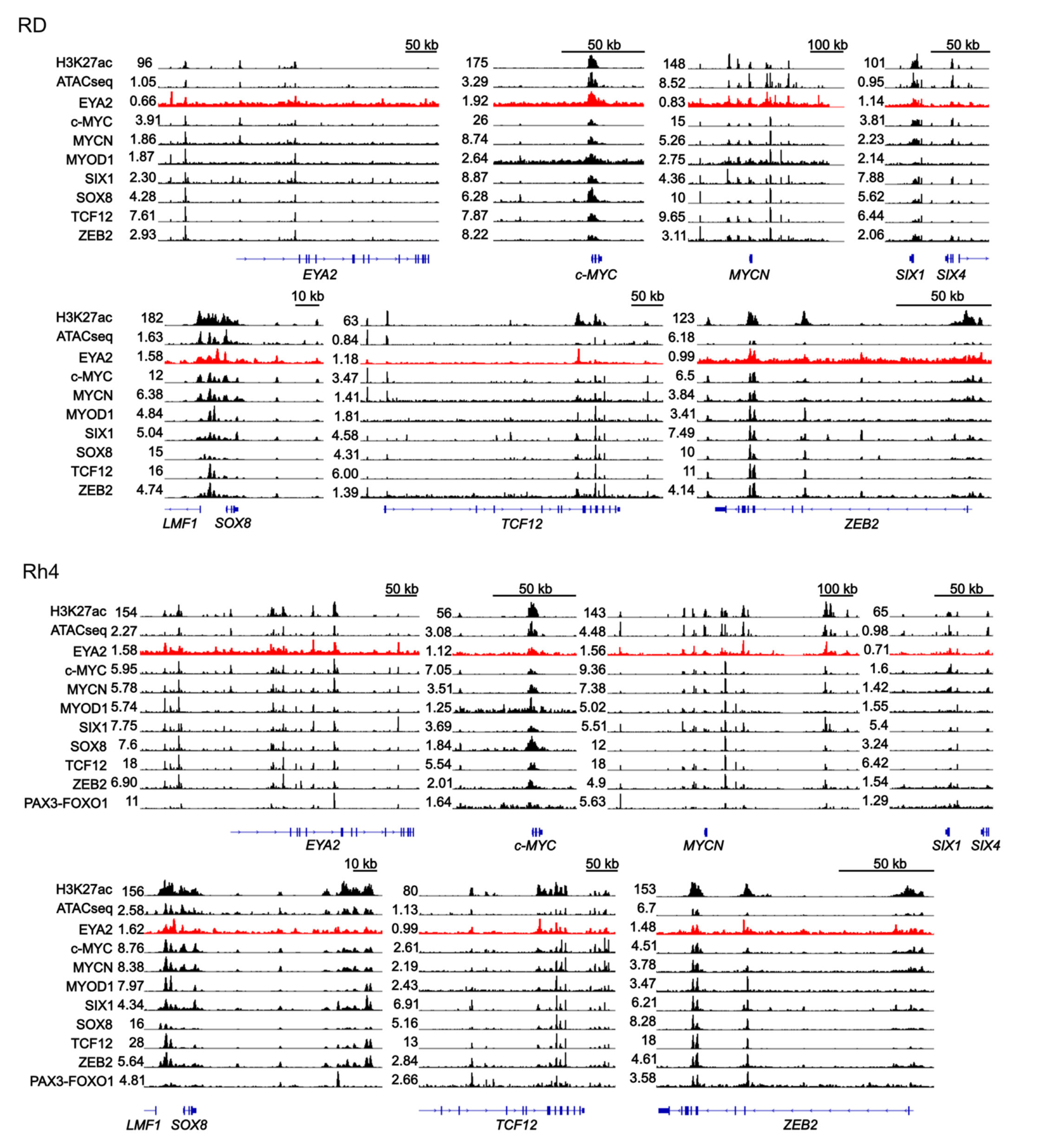

**Supplementary Figure 15. EYA2 is bound at super-enhancers regulating core regulatory circuitry mTFs.**

ChIP-seq gene tracks at CRC mTF loci in FN-RMS RD cells, and FP-RMS Rh4 cells. Tracks show co-binding at cis-regulatory elements of EYA2 and RMS CRC members (MYOD1, SIX1, SOX8, TCF12, ZEB2, MYCN, c-MYC) at regions of open chromatin resolved by ATAC-sequencing, and within super-enhancers, resolved by H3K27ac ChIP-seq.

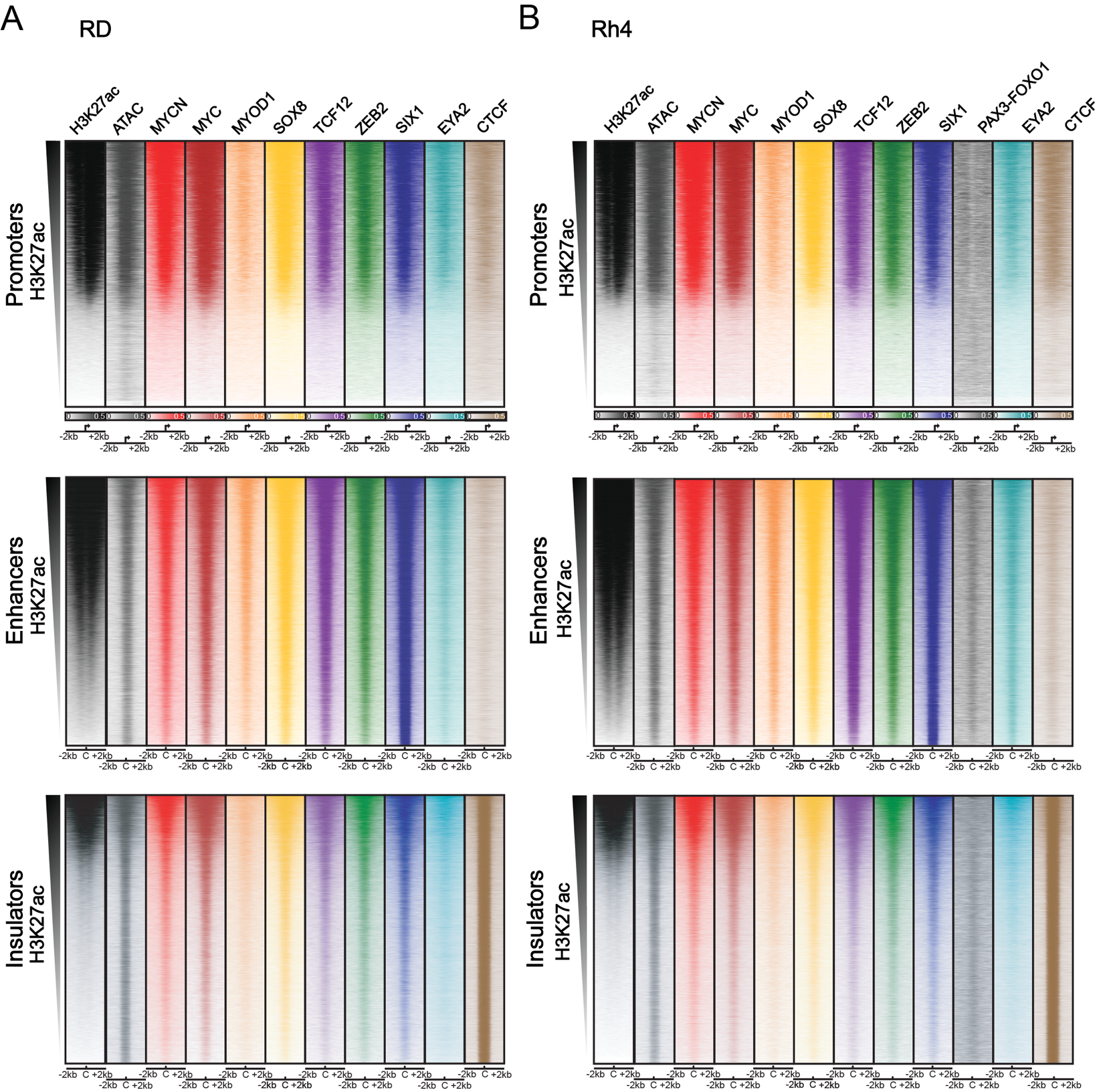

**Supplementary Figure 16. EYA2 is a co-factor associated with the pan-RMS core regulatory circuit.**

ChIP-Seq binding at individual promoters, enhancers, and insulators in RD (left) and Rh4 (right).

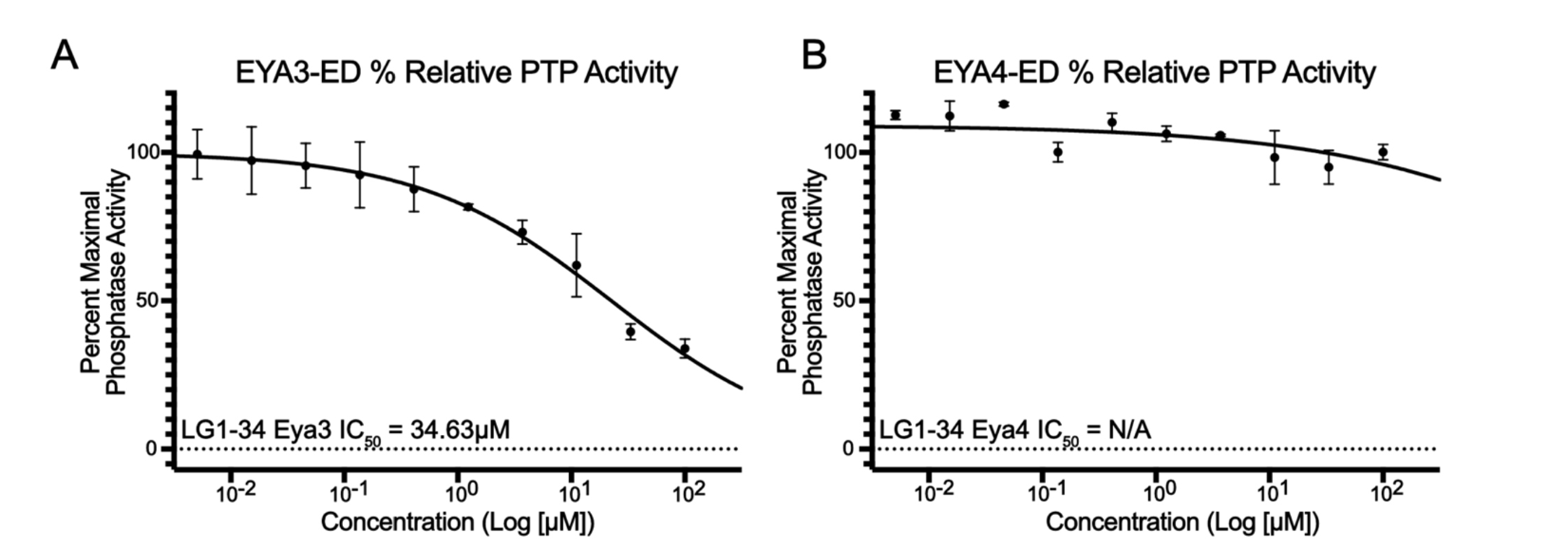

**Supplementary Figure 17. LG1-34 inhibits EYA3, but does not inhibit EYA4 tyrosine phosphatase activity.**

Tyrosine phosphatase activity of purified EYA3-ED (A) and EYA4-ED (B) in the presence of increasing

concentrations of LG1-34. Enzymatic activity was measured by fluorescent 3-O-methylfluorescein phosphate

(OMFP) assay; data are plotted as percentage of activity relative to vehicle control. Data are representative of

three independent experiments (mean ± S.E.M).

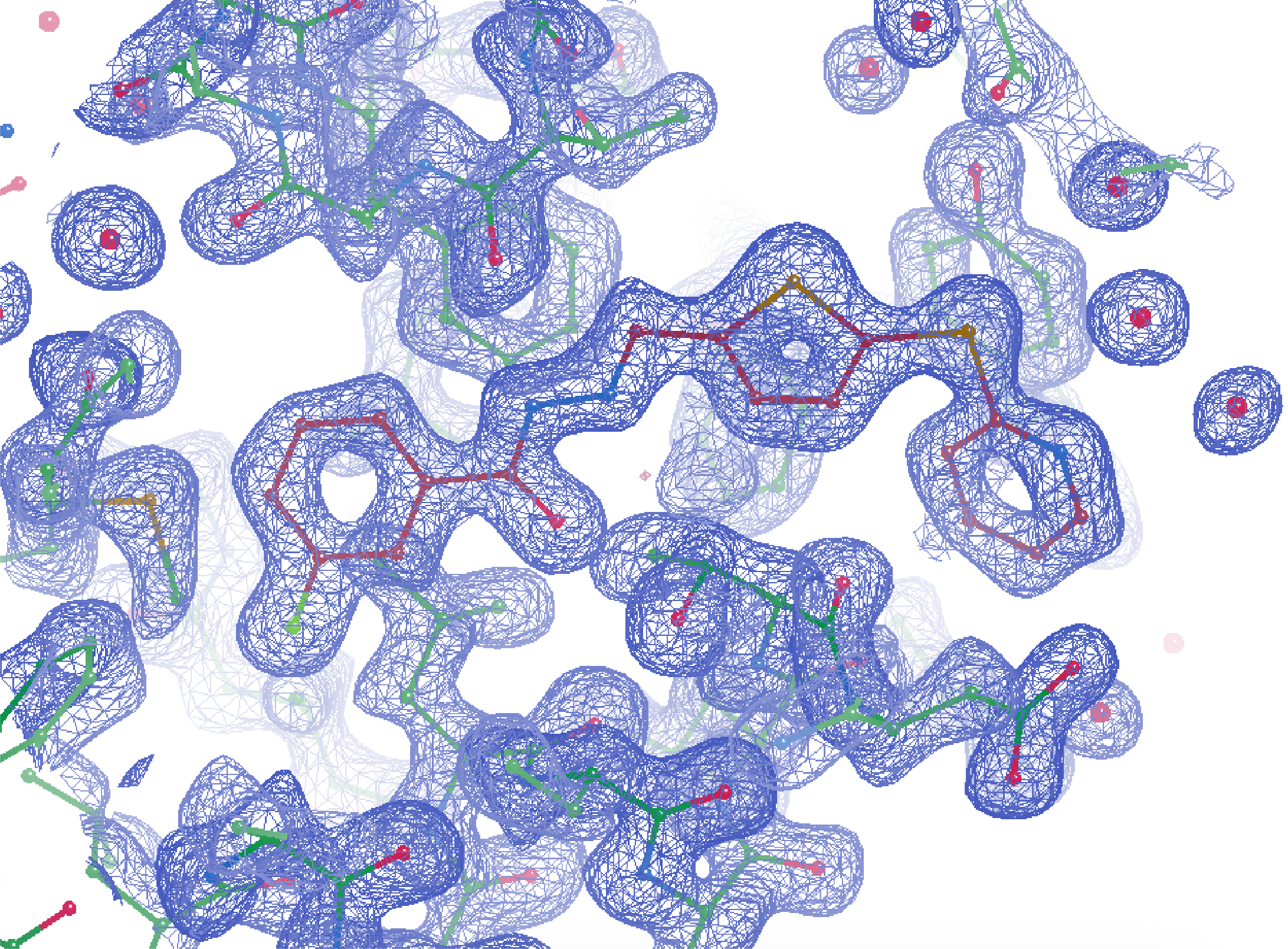

**Supplementary Figure 18. Electron density map of LG1-34 bound to EYA2.**

Electron density map of LG1-34 and surrounding EYA2 protein residues.

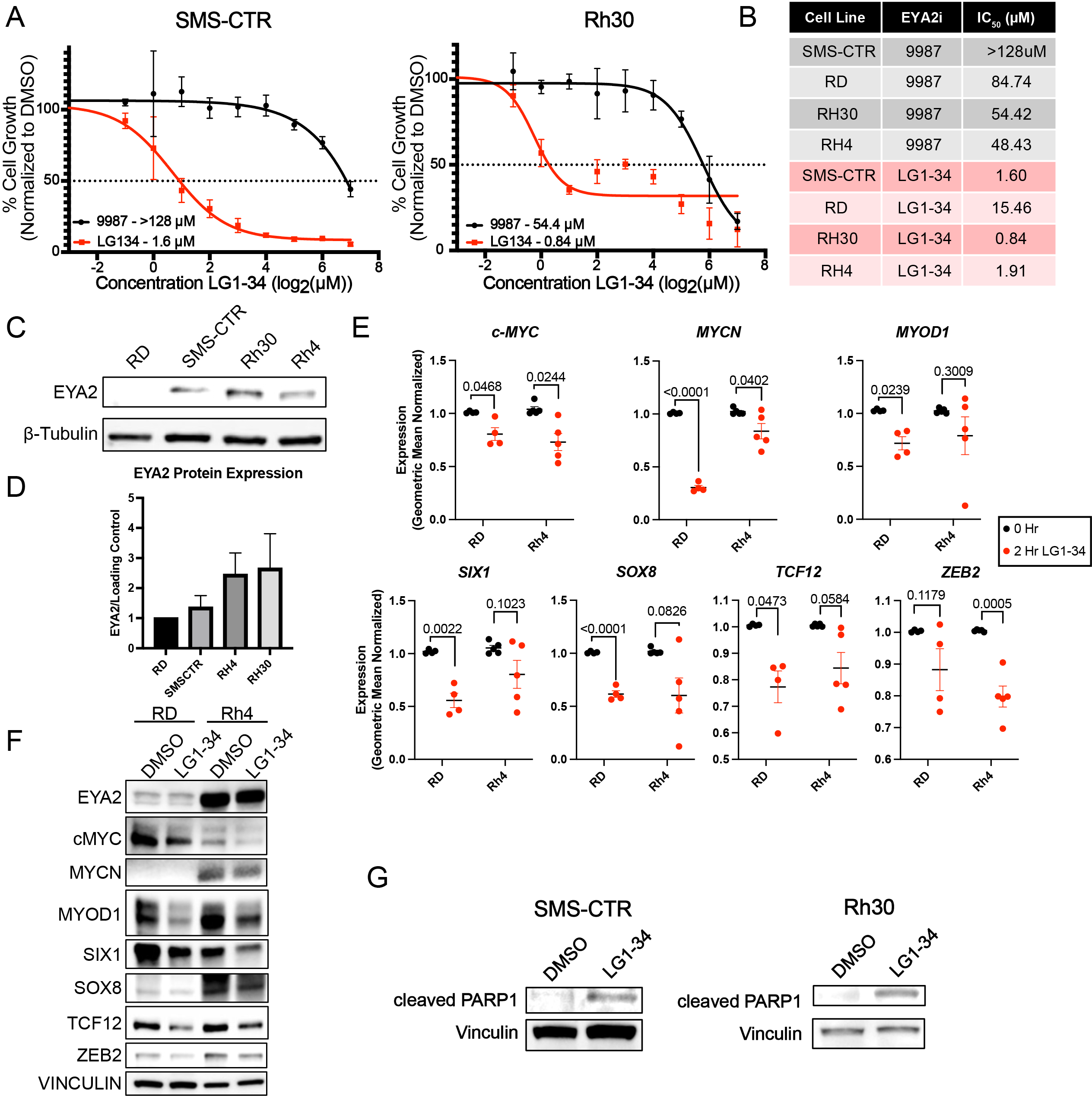

**Supplementary Figure 19. LG1-34 inhibits the RMS core regulatory circuit resulting in cell death.**

A. SMS-CTR and Rh30 cells treated with increasing doses of LG1-34 (500nM – 128µM) for 3 days. Cell

growth was measured by CellTiter-Glo. Data points are representative of three independent experiments

(mean ± SD).

B. Inhibitory concentration 50% (IC50) values for RMS cell lines treated for 3 days with 9987 or LG1-34.

C. Western Blot analysis for EYA2 across a panel of RMS cell lines. Data is representative of three

independent blots.

D. Quantification of western blots in parental RMS cell lines. Signal was normalized to loading control and fold

change over RD was calculated. Data is representative of 4 independent western blots of RMS cell lines

probed for EYA2 (mean ± S.D.).

E. Individual qRT-PCR plots reflecting summary data presented in Fig 6F. CRC gene expression determined in

RD and Rh4 treated with LG1-34 at 15 µM (RD) and 2 μM (Rh4) (72h IC50 values) for 2 hours. Data is

normalized to the geometric mean of *β-Actin, GAPDH,* and *HPRT*. Data represents > four independent

treatments (mean ± S.E.M.), and qRT-PCR reactions. Statistical differences were calculated using lognormal t-

test.

F. Western blot analysis of RMS cell lines treated with LG1-34 for 24 hr. Cell lines: RD (15µM) and Rh4 (2µM).

Blotting was performed for RMS CRC members and EYA2, with vinculin as a loading control. Images shown

are representative of three independently performed experiments.

G. SMS-CTR and Rh30 cells were treated with 2 μM (SMS-CTR) and 1 µM (Rh30) LG1-34 for 48 hr prior to

protein extraction for western blotting. Data is representative of three independent protein isolations,

treatments and immunoblots.

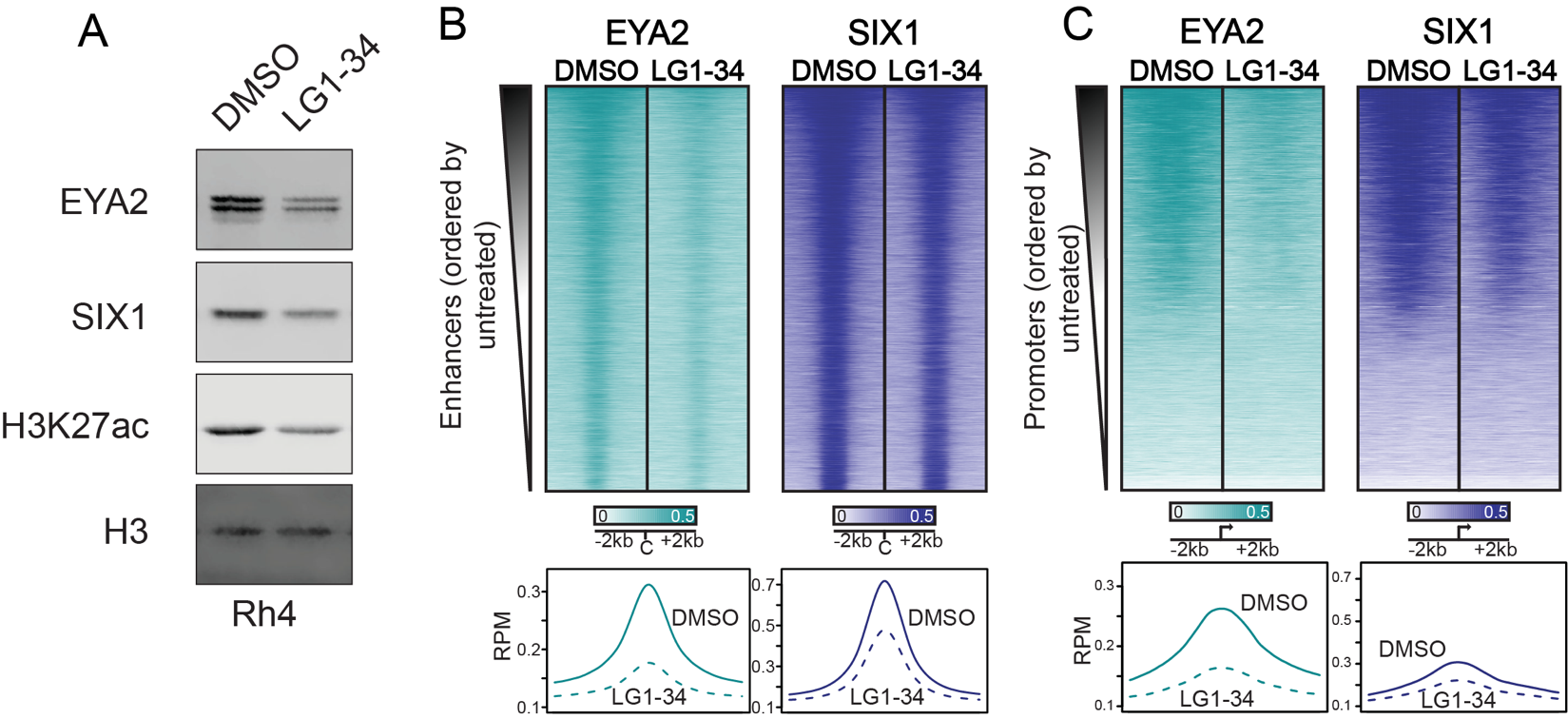

**Supplementary Figure 20. Treatment of Rh4 cells with LG1-34 decreases association of EYA2 with chromatin.**

A. Histone extracts followed by western blot of RD and Rh4 cells demonstrating decreased H3K27ac

deposition and SIX1/EYA2 chromatin binding with 24 hr treatment of 2 µM LG1-34. Blots are representative

of 3 different independent histone extractions and western blots

B. Reads-per-million-normalized ChIP-Seq of EYA2 and SIX1 at enhancers (sites bound by master TFs) or C. promoters (transcription start sites +/- 2kb) in Rh4 cells treated with DMSO or LG1-34 at 2 µM. Rows ordered by untreated H3K27ac displayed in **Figure 1E**. Metagenes show average across all heatmap rows.

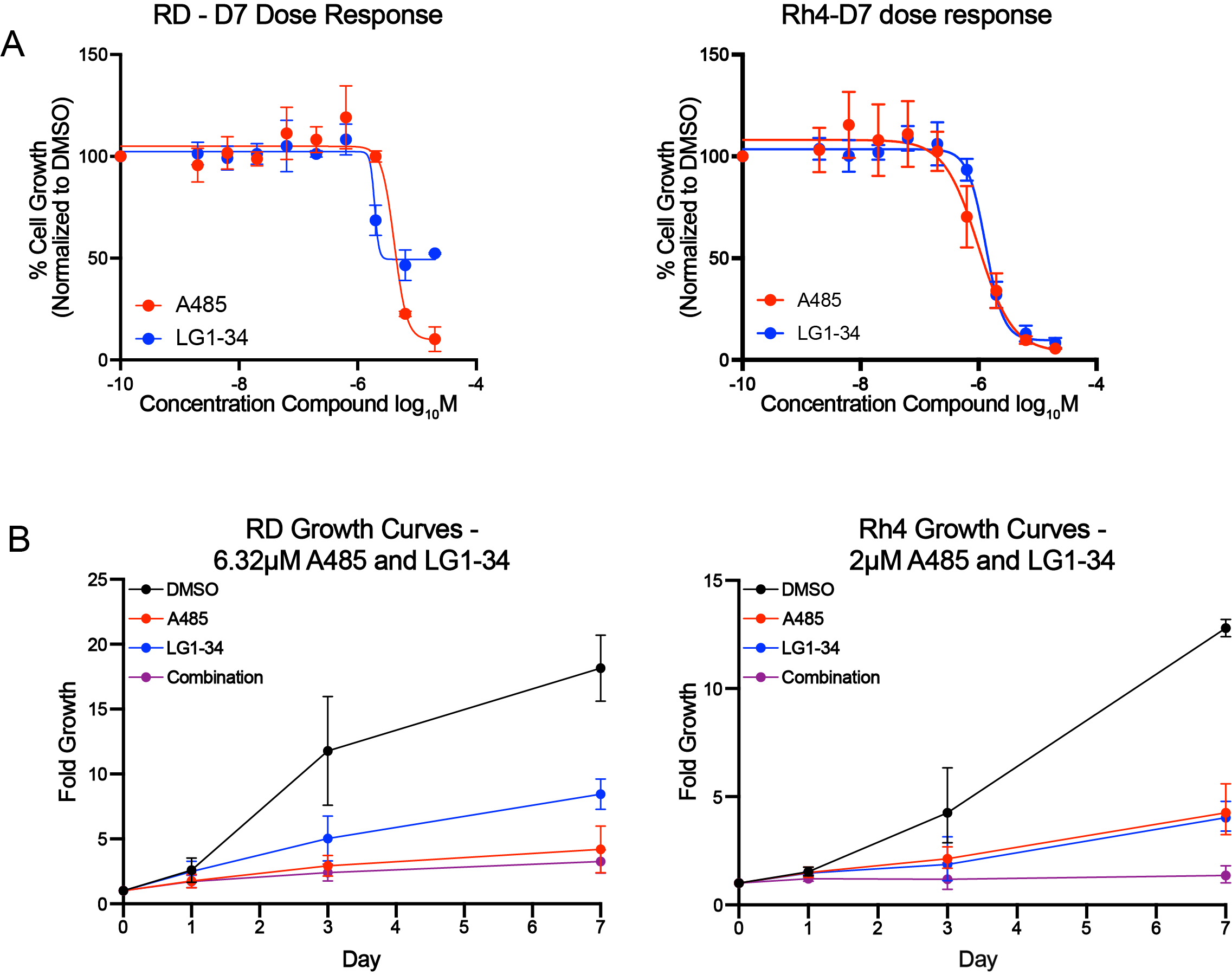

**Supplementary Figure 21. A485 and LG1-34 are synergistic *in vitro*.**

A. Dose–response curves for A485 and LG1-34 across a range of doses (2 nM–20 µM) in RD and Rh4 cells

treated for 7 days, measured by CellTiter-Glo. Data represents average of three independent experiments,

bars = S.E.M.

B. Fold growth over 7 days in cells treated with DMSO, A485, LG1-34, or the combination. RD cells were

treated with 6.32 µM A485 and 6.32 µM LG1-34; Rh4 cells were treated with 2 µM A485 and 2 µM LG1-34.

Data represents average of three independent experiments, bars = S.E.M.

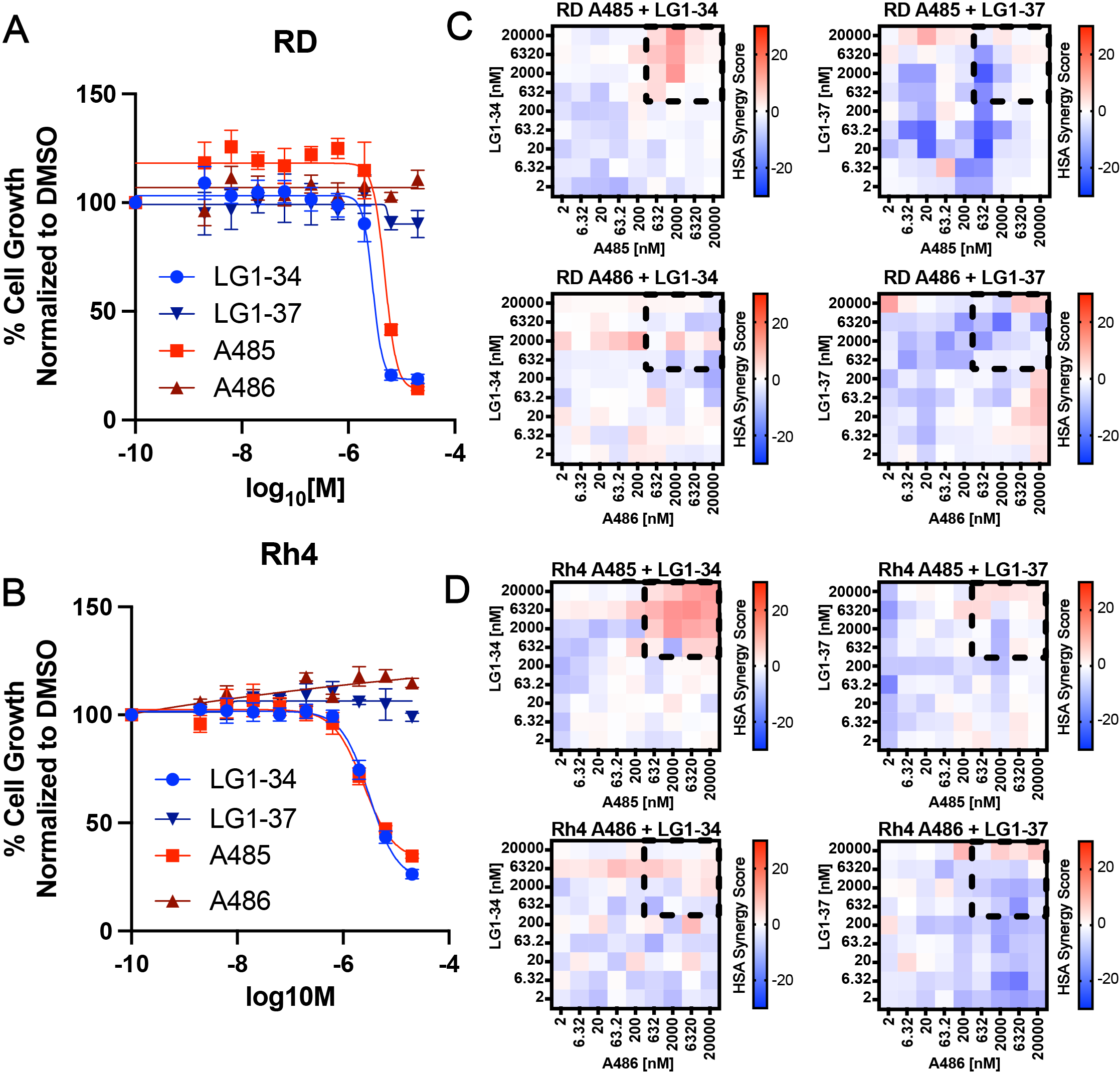

**Supplementary Figure 22. Synergy between LG1-34 and A485 is specific to active compounds.**

A. Dose–response curves for A485, A486, LG1-34 and LG1-37 across a range of doses (2 nM–20 μM) in RD

And B. Rh4 cells treated for 3 days, measured by CellTiter-Glo. Data represents average of two independent

experiments, bars = SEM.

C. Heatmaps of HSA Synergy scores calculated for A485 + LG1-34, A486 +LG1-34, A485 + LG1-37, and A486 + LG1-37 in RD cells and D. Rh4 cells. Scores were calculated using the average of two independent experiments performed in technical duplicate.

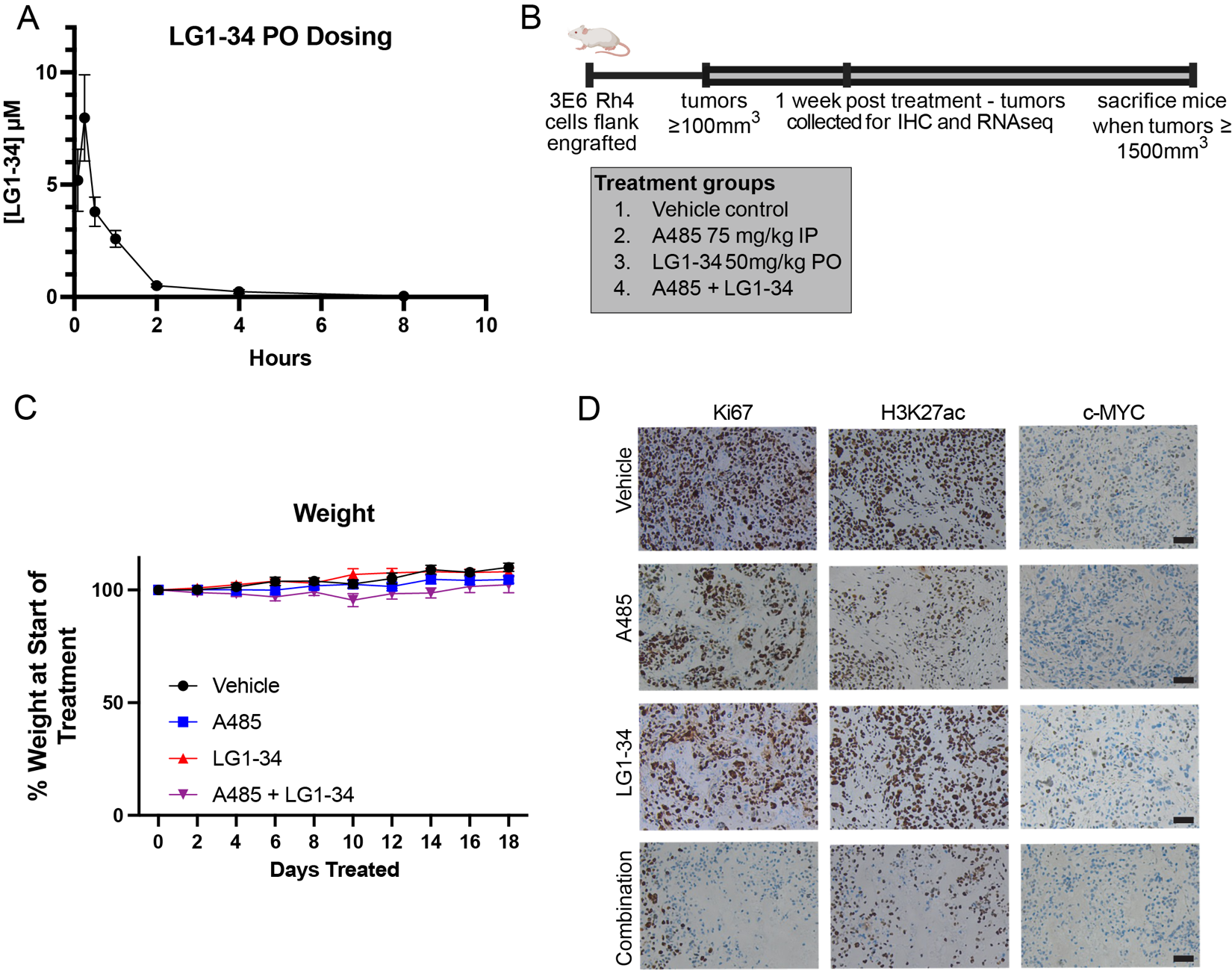

**Supplementary Figure 23. The combination of A485 and LG1-34 *in vivo* suppresses tumor growth.**

A. Estimated serum molarity of LG1-34 following QD PO dosing of 50 mg/kg based on linear extrapolation of

PO 8mg/kg PK data.

B. Diagram illustrating experimental strategy for in vivo combination of A485 and LG1-34.

C. Plot of mouse weights for duration of experiment where A485 and LG1-34 were combined. n= 8 per group.

Bars = S.E.M.

D. Immunohistochemistry for Ki67, H3K27ac, and c-MYC performed on Rh4 xenografts treated for 7 days with

vehicle, A485, LG1-34, or combination. Data is representative of three separate tumors per group, stained by

IHC. Bar = 50 μm.

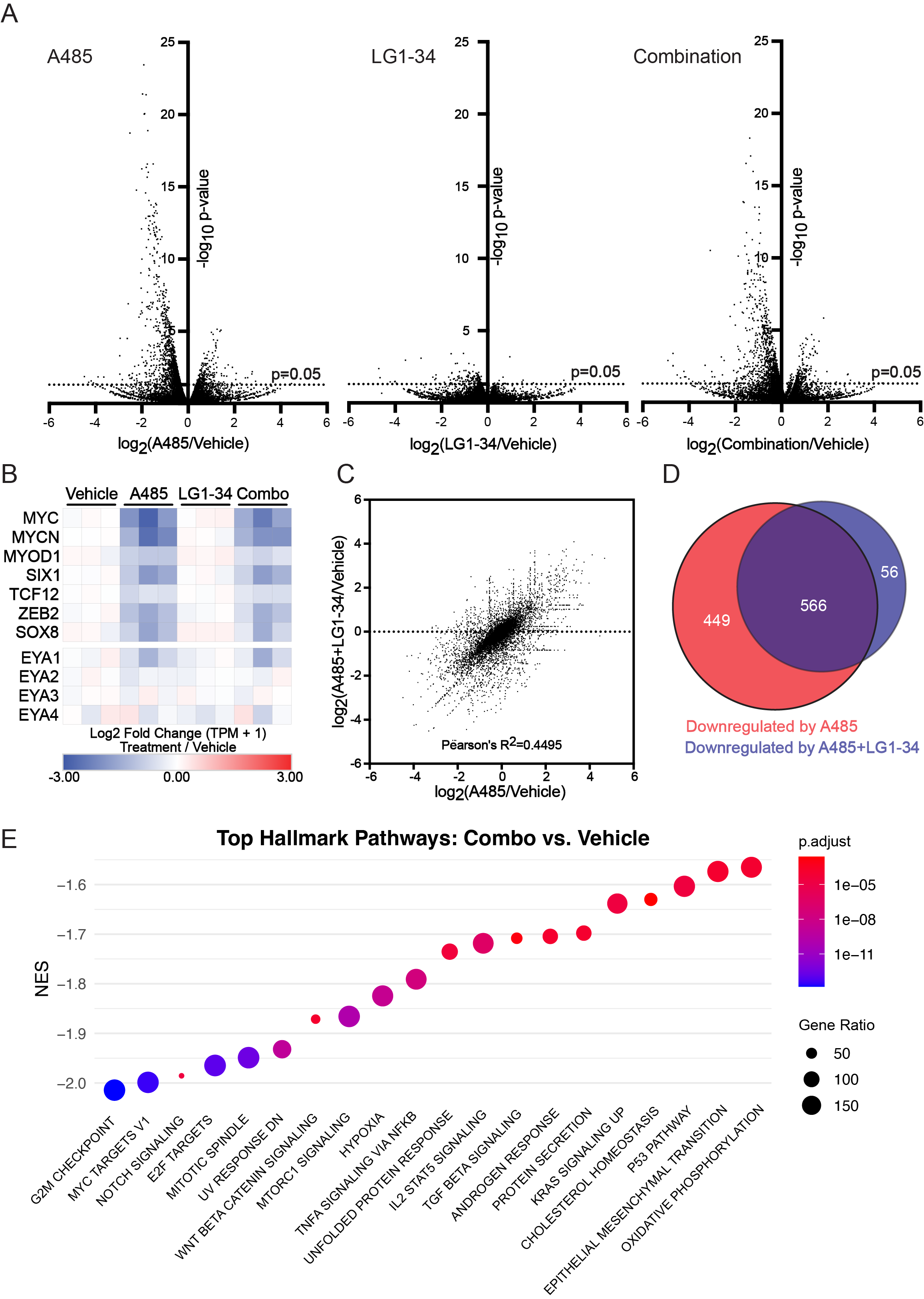

**Supplementary Figure 24. The combination of A485 and LG1-34 *in vivo* results in transcriptional dysregulation.**

A. Rh4-bearing mice were treated with vehicle control, A485, LG1-34 or A485+LG1-34 for seven days prior to

sacrifice and tumor extraction. Three tumors per group were extracted and processed for RNAseq. Demonstrated are volcano plots comparing the log2 fold change in gene expression for A485 vs. vehicle

(log2(A485/Vehicle)), LG1-34 vs. vehicle (log2(LG1-34/Vehicle)), and combination vs. vehicle (log2(A485+LG1-34/Vehicle)) plotted against adjusted p-values from a DEseq2 differential expression analysis.

B. log2 Fold Change TPM+1 in treatment groups normalized to vehicle controls heatmaps of CRC and EYA1-4

genes from RNA sequencing of Rh4 xenografts treated with vehicle, A485, LG1-34, or A485 + LG1-34 for 7d.

C. Correlation of gene expression changes between A485 vs. vehicle and combination vs. vehicle (log2(A485/

Vehicle) vs. log2(A485+LG1-34/Vehicle)), with Pearson’s R² = 0.4495.

D. Venn diagram showing overlap in downregulated genes (log2 fold change < –0.5, adjusted p-value <0.05),

comparing A485 vs. DMSO genes, and A485+LG1-34 vs. DMSO genes. 449 genes were selectively

downregulated with A485 alone, 56 genes with the combination, and 566 genes in both conditions. CRC

genes were identified in both conditions.

E. GSEA Analysis performed on DEseq2 differential expression analysis comparing vehicle treated Rh4

xenografts A485 + LG1-34 treated xenografts.

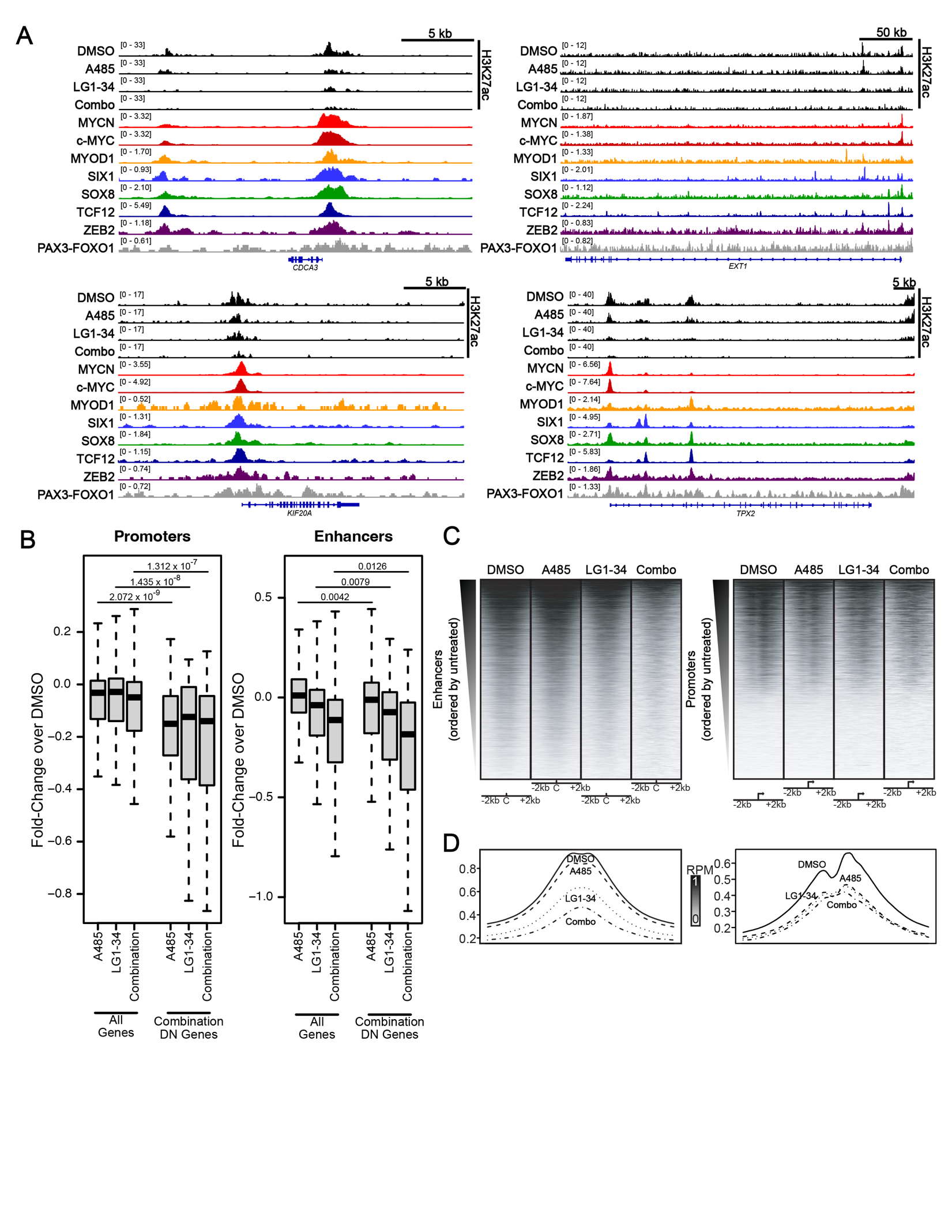

**Supplementary Figure 25. Combination treatment with A485 plus LG1-34 results in more profound loss of H3K27ac than either treatment in isolation.**

A. H3K27ac CUT&RUN tracks following 24 hr treatment with DMSO, LG1-34, A485, or the combination alongside ChIPseq tracks for members of the RMS CRC. Shown are representative genes that are dependencies in RMS and selectively downregulated with combination treatment of LG1-34 and A485.

B. Fold-Change in H3K27ac signal following 24 hr treatment with A485, LG1-34 or combination A485 + LG1-34

across (Left) all promoters or promoters of the 56 genes selectively down regulated with combination

treatment as shown in Figure 7D or (Right) all enhancers or enhancers of the 56 genes selectively down

regulated with combination treatment as shown in Figure 7D. Student T-test comparing fold-change H3K27ac

at all promoters versus 56 down regulated gene promoters with treatment of A485 (p-value = 2.072 x 10-9 ),

LG1-34 (p-value = 1.435 x 10-8), or combination (p-value = 1.312 x 10-7). Student T-test comparing fold-change H3K27ac at all enhancer versus 56 down regulated gene enhancers with treatment of A485 (p-value = 0.0042), LG1-34 (p-value = 0.0079), or combination (p-value = 0.0126).

C. Spike-in-normalized CUT&RUN coverage heatmaps at enhancers (above, defined by master TF binding

peaks) or promoters (below, transcription start sites +/-2kb). Row ordering by untreated H3K27ac shown in

Figure 1E.

D. Metagene averages across all sites in heatmaps.

**Supplementary Tables:**

**Supplementary Table 1. DepMap 25Q2 analysis of 12 rhabdomyosarcoma cell line selective dependencies, compared with 1171 other cancer cell lines.**

**Supplementary Table 2. DepMap rhabdomyosarcoma cell line selective dependency genes.**

**Supplementary Table 3. Super-enhancer-associated genes in 10 rhabdomyosarcoma cell lines.**

**Supplementary Table 4. STRING database-defined interactors of each core regulatory circuitry master transcription factor.**

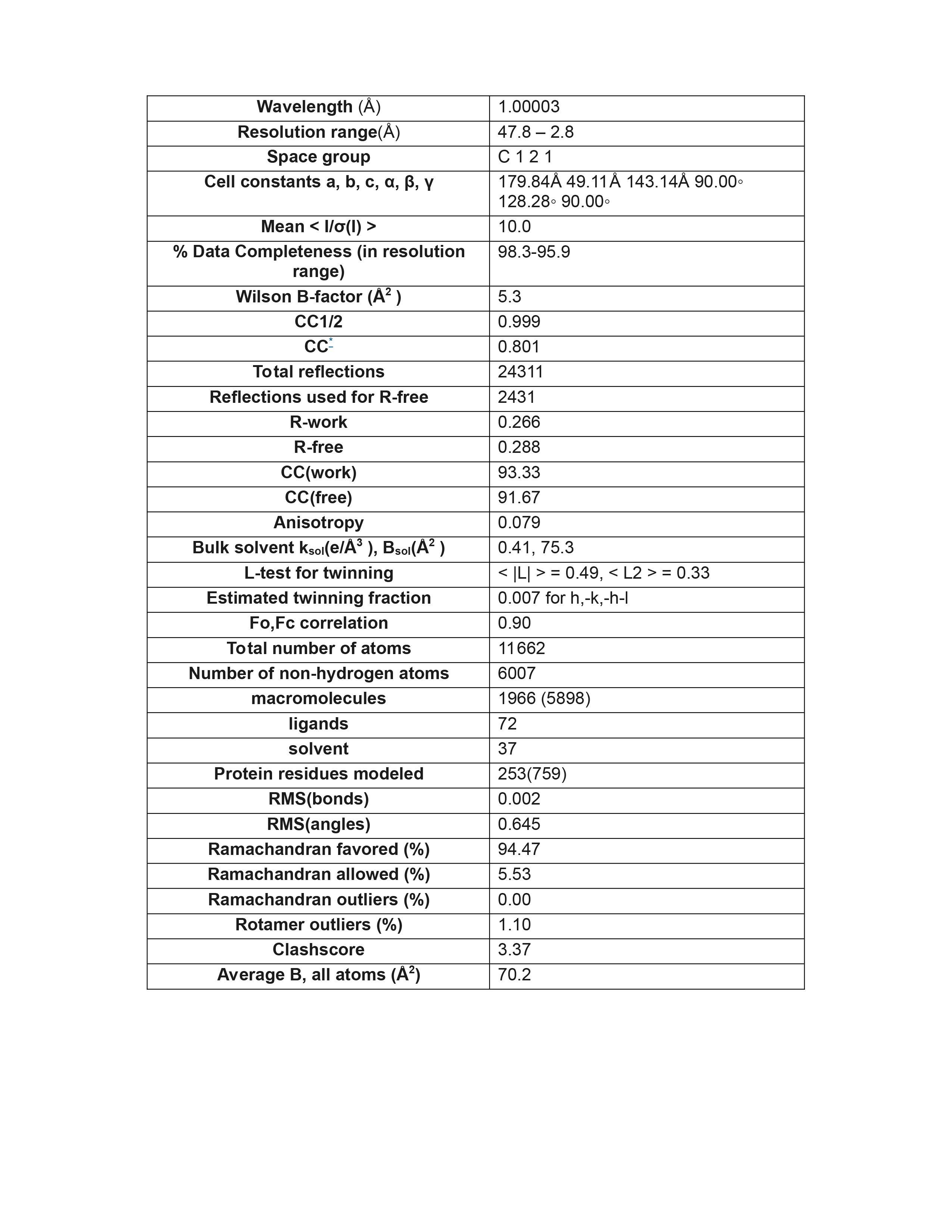

**Supplementary Table 5. Crystal structure parameters for EYA2-ED bound to LG1-34.**

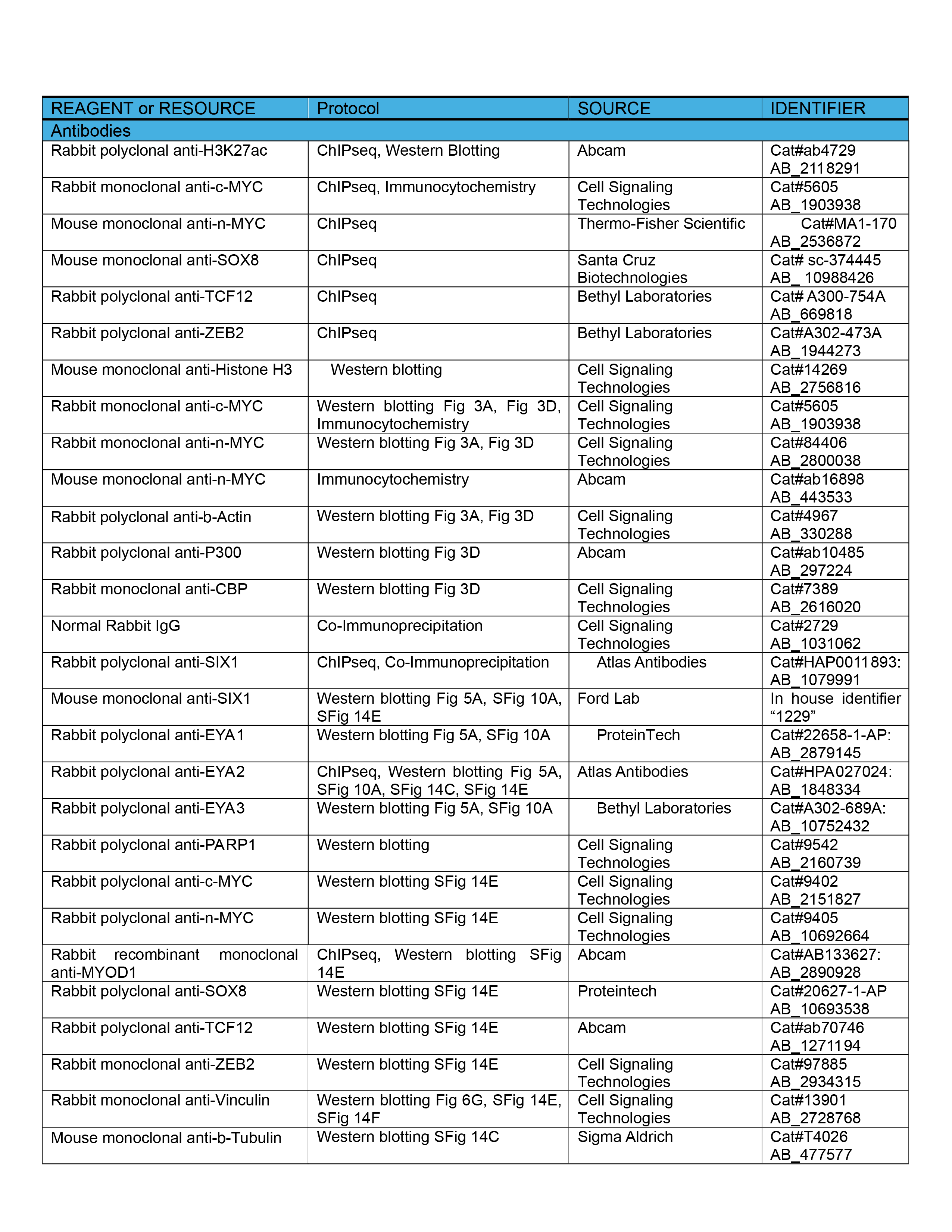

**Supplementary Table 6. Antibodies used in this study.**

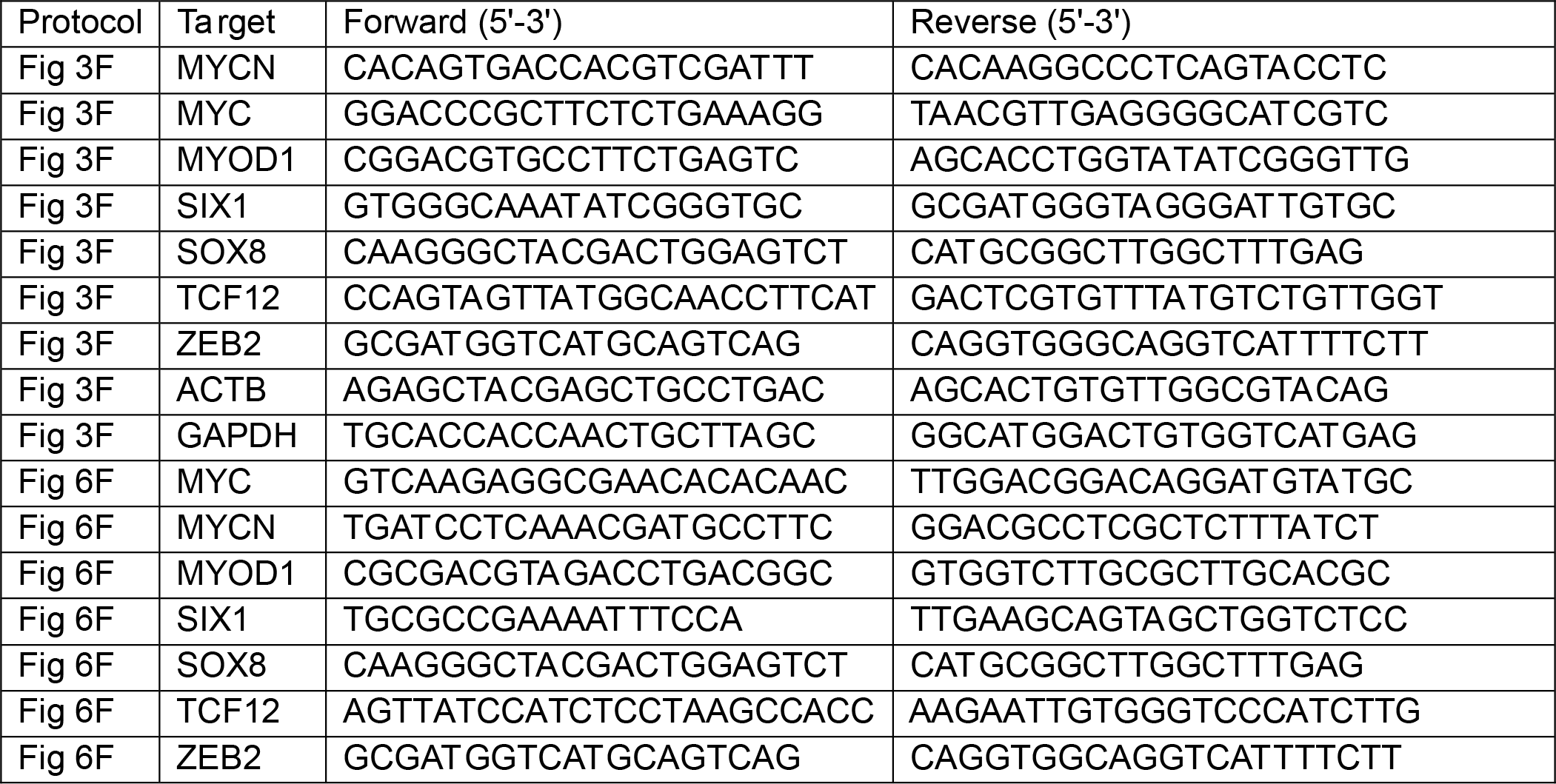

**Supplementary Table 7. Primers used for q-RT-PCR.**

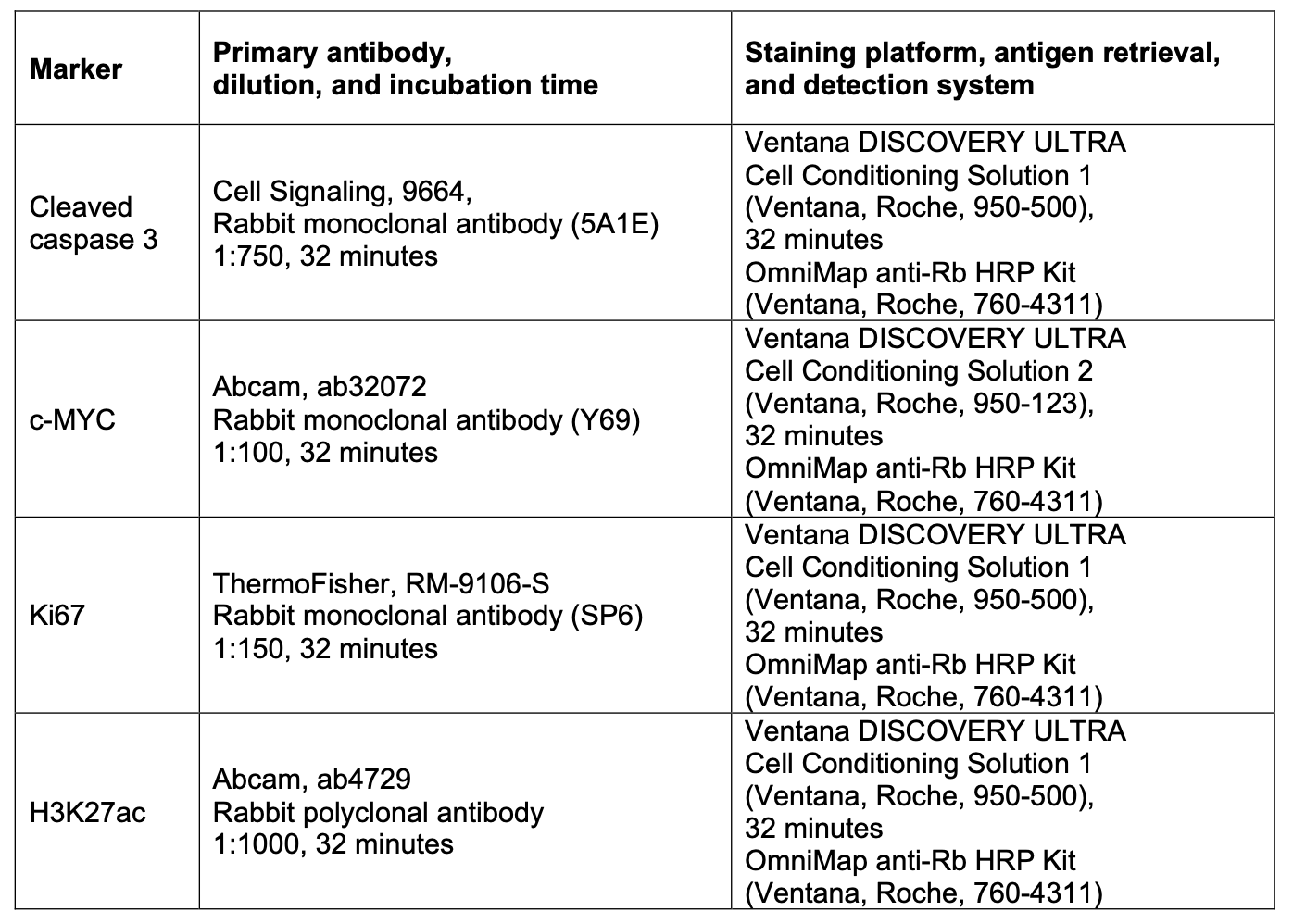
**Supplementary Table 8. Conditions and antibodies used for immunohistochemistry.**
